# Decoding the reproductive system of the olive fruit fly, *Bactrocera oleae*

**DOI:** 10.1101/481523

**Authors:** M-E Gregoriou, M Reczko, K.T Tsoumani, K.D Mathiopoulos

## Abstract

A great deal of behavioral and molecular interactions between male and female insects takes place during insect reproduction. Here, we comprehensively analyze the reproductive system of the olive fruit fly. Specifically, transcriptomic and genomic analysis of the reproductive tissues from virgin and once mated insects were performed. Genes encoding proteins implicated in immune response, mucins, antigen 5 proteins, proteases inhibitors and proteins with putative secretory activity were identified. Comparison of the transcriptomes between virgin and mated insects resulted in the identification of genes that are up- or down-regulated after mating. In testes 106 genes were up-regulated and 344 genes were down-regulated, whereas in male accessory glands with ejaculatory bulb 1,607 genes were up-regulated and 384 genes were down-regulated in mated male insects. Respectively, in mated females 1,705 genes were up-regulated and 120 genes were down-regulated in mated insects. To get a deeper insight, the expression profiles of selected genes throughout sexual maturation for the male tissues and throughout different time points after mating for the female reproductive tissues were determined. Identification of genes that take part in the mating procedure not only gives an insight in the biology of the insects, but it could also help the identification of new target genes in order to disturb the reproductive success of the olive fly and thus develop alternative pest control method.

## 1. Introduction

In all arthropods, insects are the most divergent and abundant group, equipped with high reproductive rates and numerous behavioral and physiological adaptations that are critical to the maintenance of their populations. During insect reproduction, sperm and seminal fluid that are produced in testes and male accessory glands (MAGs) respectively are delivered to the female reproductive system ^1, 2, 3^.

Male seminal fluid proteins have been characterized mainly as proteases, peptidases, serpins and protease inhibitors ^4, 5, 6^. Although the functional classes of these proteins are conserved across species, the genes that encode them rarely are. Genes expressed in the accessory glands, as it was previously shown in Drosophila species, exhibit rapid evolutionary change and gene expansion because of their critical role in encoding products that underlie striking, fitness-related phenotypes ^7^. Until today, male seminal fluid proteins have been identified in different Drosophilidae ^8, 9^, in the major disease vectors *Anopheles gambiae* ^10^ and *Aedes aegypti* ^11^, and in Tephritid fruit flies like *Ceratitis capitata* ^12, 13^ and *Bactrocera cucurbitae* ^14^.

Female accessory glands also produce a secretory material that serves a number of functions. It may act as a lubricant for egg passage, as a protective oothecal cover, or as a glue to attach eggs to various substrates ^15^. To date, female reproductive genes have been comprehensively studied in very few insects like the sandfly *Phlebotomus papatasi* ^16^, the house fly *Musca domestica* ^17^ and the Mediterranean fruit fly *C. capitata* ^18, 19^.

After mating, the presence of sperm and seminal fluid in female insects induces multiple physiological and behavioral changes such as repression of sexual receptivity to further mating ^20, 21, 22, 23^, egg-laying stimulation, stimulation of immune responses and reduced longevity ^24, 25, 12, 26^. These post-mating responses have been addressed in genome-wide studies in species such as *Drosophila melanogaster* ^27, 28^, the honeybee queen *Apis mellifera* ^29, 30, 31^, *C. capitata* ^32^, *An. gambiae* ^33, 34^ and *Ae. aegypti* ^35^. Such studies have revealed that post-mating responses differ between different insect species.

For the olive fruit fly, *Bactrocera oleae*, the major pest of olive cultivation, the morphology and ultrastructure of the male accessory glands with ejaculatory bulb tissues have been analyzed by Marchini et al ^36^. The first molecular analysis of the reproductive system was presented in 2014 by Sagri et al ^37^. That study focused on the identification of sex differentiation genes, i.e. differentially expressed genes either in male testes or female accessory glands and spermathecae ^37^. In order to go a step further and understand the molecular processes that underlie reproduction of the olive fly, one needs to identify genes that are over- or under-expressed during mating of the flies. The usefulness of the knowledge of such genes and mechanisms in insect control is obvious: if one could somehow impair the function of these genes, one could also disturb the reproductive success of the olive fly and lead to population control.

In this study, an extensive transcriptomic and genomic analysis of the reproductive tissues from virgin and once-mated olive flies was performed. Comparison of the different conditions resulted in the identification of differentially expressed genes. To investigate their contribution in reproduction, we analyzed the expression profiles of selected genes throughout the sexual maturation for the male tissues and throughout different time points after mating for the female reproductive tissues.

## 2. Results and Discussion

The tissues selected for the transcriptomic analysis of the olive fly’s reproductive system were: 1. the male accessory glands from virgin (V_MALE) and mated (M_MALE) male insects, 2. the testes from mated insects (M_TESTES) and 3. the lower female reproductive tract, comprising of the spermathecae, the uterus and the female accessory glands from virgin (V_FEMALE) and mated (M_FEMALE) female insects.

In order to obtain tissues from sexually mature virgin flies, dissections were performed on DAY-7 after eclosion. The sexual maturation of the insects was determined by their ability to mate and give offspring. The olive fly laboratory strain used for the experiments is sexually mature and can mate successfully at the selected day. Dissection of the tissues from mated olive flies was performed 12 hours after mating. Specifically, on DAY-7 after eclosion virgin male and female flies were mixed and allowed to mate. A successful mating lasts for at least one hour ^38^. When the insects completed mating (i.e., they were voluntarily separated from each other), they were kept in different cages for 12 hours before dissections. In *D. melanogaster* the highest post-mating gene expression occurs after 6 hours ^28^ and in *C. capitata* there is a general increase in the transcriptional activity only after three repeated matings ^39^. As there is no such evidence for the reproductive tissues of *B. oleae*, two pieces of information guided our decision for the determination of the appropriate time-point after mating to analyze. Firstly, male olive flies can remate at least 24 hours after a previous mating ^40^ and, secondly, oviposition of the mated females also starts 24 hours after mating.

### 2.1. Transcriptome sequencing assembly

A single representative *de novo* assembly for *B. oleae* was generated from a concatenation of the libraries obtained with the Illumina platform ^41^ using the Trinity pipeline ^42^. After assembly, transcript and unigene level expression values were calculated using RSEM ^43^ for the four libraries (V_MALE, V_FEMALE, M_MALE, M_FEMALE) sequenced using the Ion Proton. The average read length was 669.78bp and the totally sequenced 203,690,146 bp gave 255,077 genes. From the libraries V_MALE and M_MALE 11,452 transcripts were obtained whereas from the libraries V_FEMALE and M_FEMALE 10,478 transcripts were obtained.

### 2.2. Before vs after mating differential expression

To identify the differentially expressed genes between virgin and mated insects, the edgeR algorithm ^44^ with a stringent cutoff (q value < 0.05) was used. Specifically, in the reproductive tissues from the mated female insects, 1,705 genes were up-regulated and 120 genes were down-regulated. On the other hand, in male accessory glands with ejaculatory bulb 1,607 genes were up-regulated and 384 genes were down-regulated, whereas in testes 106 genes were up-regulated and 344 genes were down-regulated. Less differentially expressed genes were detected in testes, compared to the other two reproductive tissues, showing a limited transcriptional activity. Indeed, in *D. melanogaster*, it was shown that spermatozoa are generally metabolically quiescent and transcriptionally silent in adult insects ^45, 46^. The entire lists of all significantly (q<0.05) differentially expressed genes in all analyzed tissues are given in supplementary Tables S1, S2, S3. Moreover, volcano plots were created for the visualization of the distribution of the transcripts; the red dots indicate differentially expressed genes between virgin and mated insects (Figure 1).

**Figure 1:**
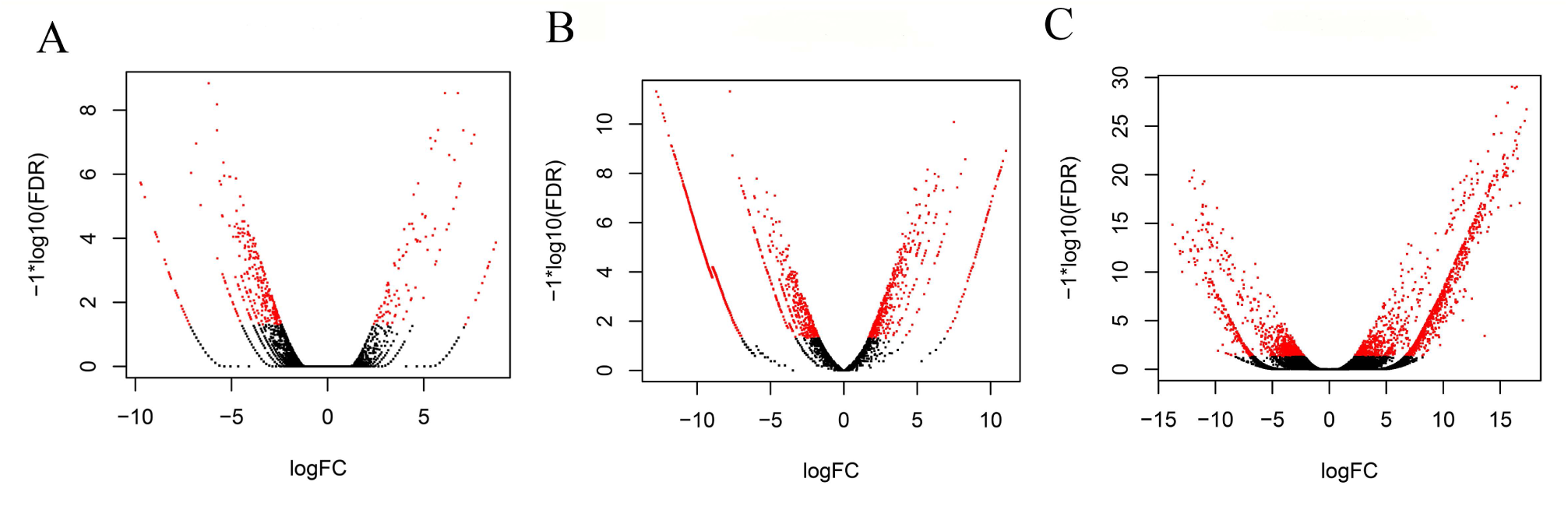
Volcano plots represent the differentially expressed genes between virgin and mated flies in the three dissected tissues: A. testes, B. male accessory glands with ejaculatory bulb, C. lower female reproductive tract. The Y axis represents significance and the X axis represents logarithmic fold change. The red dots represent differentially expressed genes (p value < 0.05).

Functional annotation of the top100 over-expressed transcripts in the tissues of the mated flies was performed based on the Gene Ontology (GO) categorization level II using BLAST2GO tool ^47^. The GO categorization is presented in Figure 2 and involves biological process (BP) and molecular function (MF). The Gene Ontology (GO) terms of the testes transcriptome, with the most abundant hits on the GO functional annotation based on categorization level II, were similar to those obtained from the *Bactrocera dorsalis* respective testes-transcriptome ^48^. Regarding the “molecular function” classification, the identified genes mostly fell in the main groups “binding” and “catalytic activity”. These groups were also identified as the most abundant in similar studies in *B. dorsalis* 48 and *C. capitata* ^32, 39^, two closely related insects to the olive fly, showing a conservation of the functions that alter during mating in these insects. Regarding the male accessory glands and ejaculatory bulb tissues, the GO analysis in response to biological processes and molecular function showed enrichment of “metabolic processes” and “biological regulation” as in *C. capitata* ^32^. As sexually mature males are actively involved in pheromone response and female courting, they show significant enrichment of these GO terms, indicating the high energy investment required in mating. Finally, as it was extensively analyzed in many Drosophila species, the female reproductive genes encode proteases, protease inhibitors and genes related to immune response and energy metabolism ^28, 27, 49, 50^. In consistence with those data, homologous genes were also observed in the GO annotation of the upregulated genes in the lower female reproductive tract of *B. oleae*. This transcriptional activity of mated olive flies is characterized by rapid cell proliferation and secretory activity, as supported by the categorization of the transcripts in functional classes related to biological regulation, metabolic and cellular processes. As it comes to olive fly, indeed the general transcriptomic profiles of the analyzed tissues were similar to other dipteran reproductive systems such as *C. capitata* ^32, 39^. However, a more detailed analysis of the transcripts showed that there is diversity in the mating response among species. Specifically, compared to *C. capitata,* there were two distinct differences. Firstly, there was a profound increase in the number of transcripts in once mated *B. oleae* insects. This rise in the number of transcripts was detected only in three times mated *C. capitata* insects. Secondly, in *B. oleae* there was a modification of the immunity response of the reproductive tissues while in *C. capitata* there were not ^32^. Similarly, comparison of *D. simulans* male accessory gland proteins with their orthologues in its close relative *D. melanogaster* demonstrated rapid divergence of many of these reproductive genes ^51^. The divergence of the reproductive genes is based on the important role that they play in ensuring the successful mating and fertilization ^52, 53^.

**Figure 2:**
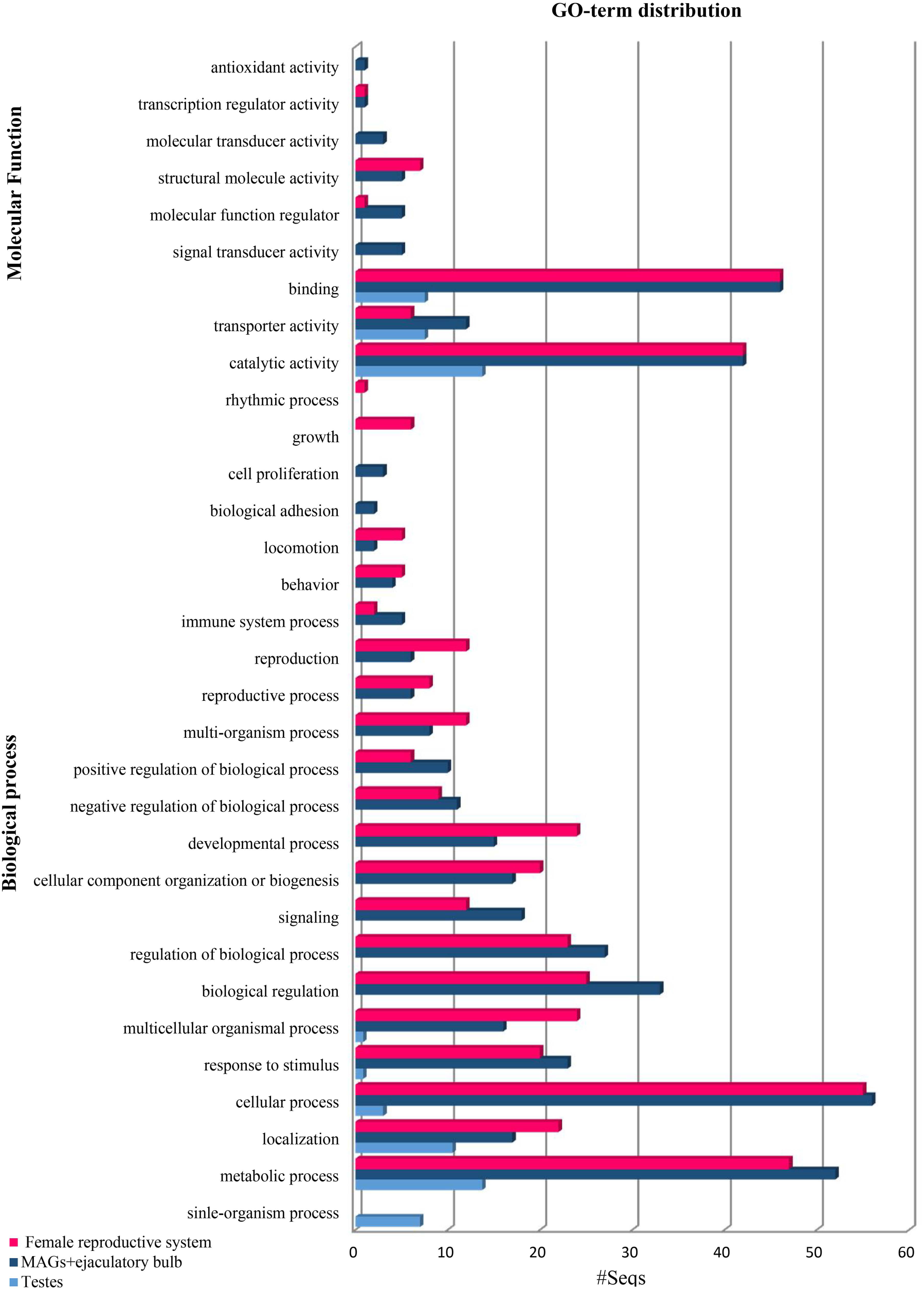
Functional annotation of the top 100 over expressed genes in *B. oleae* reproductive tissues from mated insects showing top 20 hits of different category for molecular function (MF) and biological process (BP).

For the validation of the differential expression of various genes observed after the RNAseq analysis of reproductive tissues before and after mating, further functional analysis was performed for 5 loci in testes and 6 loci in male accessory glands and female accessory glands, respectively. These genes were selected based on their known involvement in the reproductive system and their differential expression between virgin and mated insects. In testes of mated flies, significant overexpression was confirmed for the genes *c58283*, *c37552*, *hemolectin*, *mucin* and *cation transporter* and downregulation for *scribbler* gene. qRT-PCR did not confirm the expression profile of the genes *c15699* and *c52071* obtained from RNAseq, whereas *c42528* showed very low expression (Figure 3). In male accessory glands of mated flies, six reproductive loci were analyzed and their overexpression was confirmed through qRT-PCR: *timeless*, *c52416*, *c57257*, *c52655*, *yellow-g* and *c53574* (Figure 4A). In female accessory glands, qRT-PCR results showed that *troponin C* had high expression (18 fold) in virgin flies, in contrast to the RNAseq results that showed overexpression in mated female flies. The *glutathione S-transferase epsilon class* gene showed very low expression. Overexpression of the other four genes, *lingerer*, *yolk protein-2, bestrophin-2* and *ornithine decarboxylase antizyme* observed in RNAseq was also confirmed in qRT-PCR (Figure 4B).

**Figure 3:**
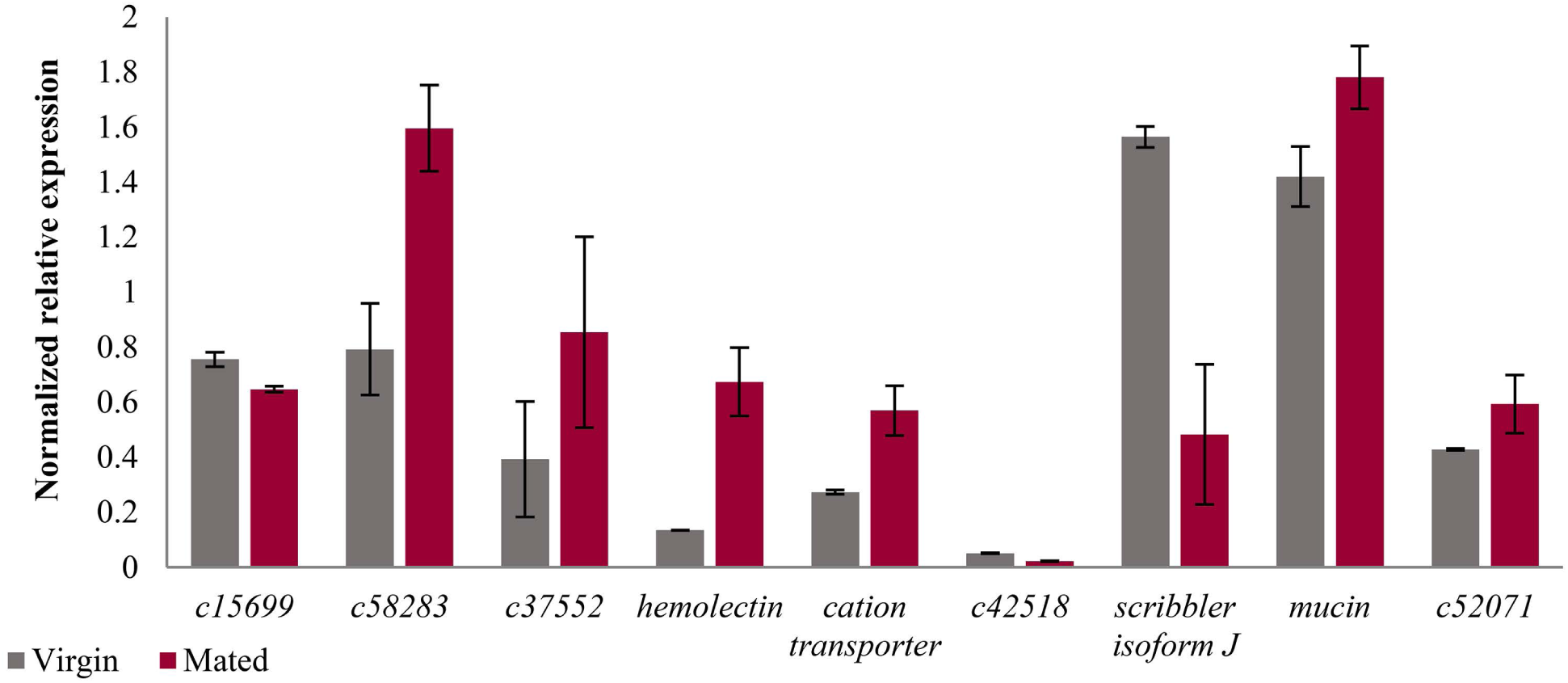
Relative expression profiles of genes expressed in the reproductive tissue of testes from virgin and mated male insects. Mean values ± standard error of triplicate data from three biological replicates are shown. No statistical significance as determined by t-test (p< 0.05).

**Figure 4:**
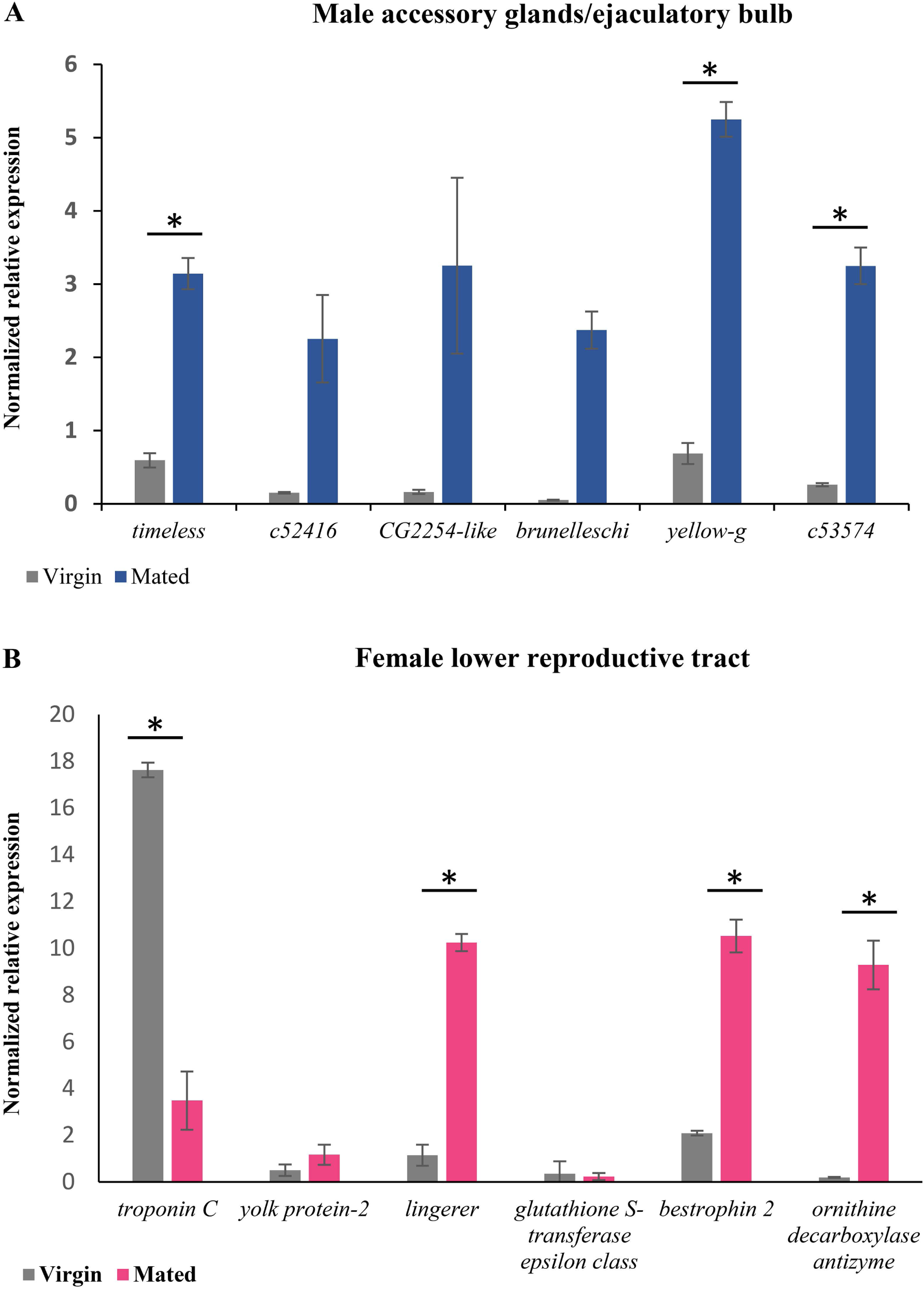
Relative expression profiles of genes expressed in virgin and mated insects from: A. male accessory glands with ejaculatory bulb and B. flower female reproductive tract between virgin and mated female insects. Mean values ± standard error of triplicate data from three biological replicates are shown. The * indicates significantly different, as determined by t-test (p< 0.05).

### 2.3. Genomic annotation of reproductive genes

The 100 highly differentially expressed genes were annotated to the recently sequenced genome of the olive fly (https://i5k.nal.usda.gov/Bactrocera_oleae) (Table S2). In an attempt to annotate more seminal fluid proteins (SFPs), the genome scaffolds were queried (tBLASTn, e-value <10^−10^) using the amino acid sequences of the 139 characterized *D. melanogaster* SFPs ^8^. Only 43 of the Drosophila genes gave significant hits in the olive fly genome. The homologous genes were grouped into 17 functional classes based on the categories defined for *D. melanogaster* ^9^ and *C. capitata* ^13^ seminal fluid proteins (Table S4). The annotated genes obtained from this procedure encode proteins that belong to the conserved functional classes such as proteases and protease inhibitors, lipases, sperm-binding proteins and antioxidants ^54^. Four sperm protein genes including the testes-specific protein *betaTub85D* and one odorant binding receptor, *or82a*, were also annotated.

### 2.4. Transcriptional analysis of the differentially expressed genes

In order to get a closer look into the transcriptional changes of the reproductive genes, a more detailed follow-up expression profile analysis was performed for the male accessory glands with ejaculatory bulb tissues and the female lower reproductive tract.

For the male accessory gland genes, we determined the expression profiles of the selected genes from DAY-0 (first day of insect eclosion) to DAY-7 (sexually mature insects). Presumably, a gene encoding a seminal fluid protein should be expressed before mating, so that the protein will be present at the time of mating. In agreement with this hypothesis, the highest expression of most genes was detected before DAY-7 (Figure 5). The genes *timeless*, *c52416* and *c53574* showed highest expression on DAY-0, while their expression dropped to lower levels and remained stable until DAY-7. The genes *brunelleschi* and *yellow-g* showed their highest expression on DAY-5 and gene *CG2254-like* on DAY-6.

**Figure 5:**
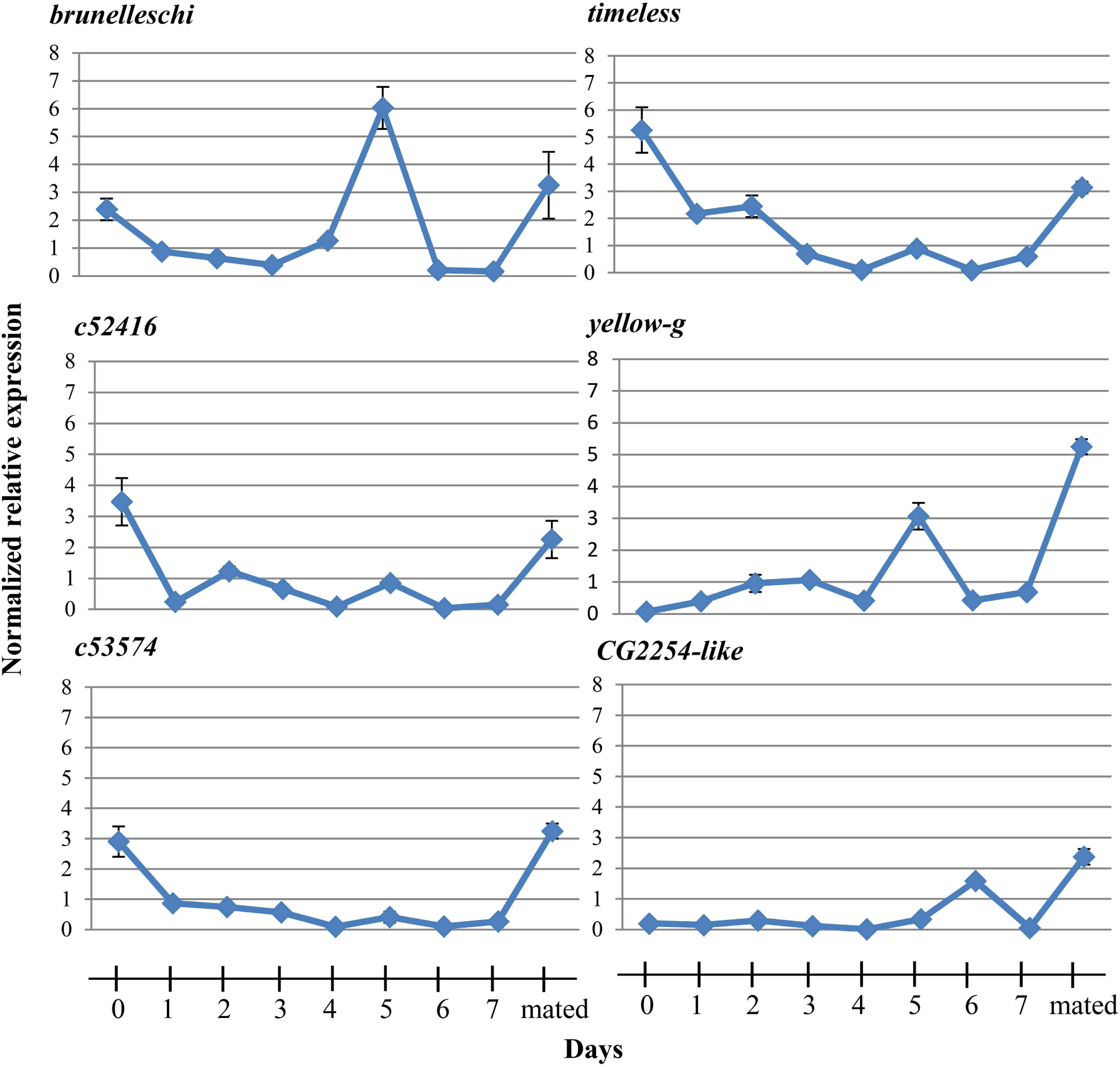
Expression profiles in female reproductive tissues. Expression profiles of the selected genes from the first day of eclosion (DAY-0) until DAY-7. The error bars show the standard error of the mean between the three biological samples.

*Timeless* expression increased after mating, indicating possibly a role in the rhythmic cycle. *Timeless* along with *per* (*period*) regulate the circadian cycle of insects. Knockout of *timeless* in male *D. melanogaster* showed a change in mating time ^55^. In *Spodoptera litorralis* it has been demonstrated that the sperm release rhythm is controlled by an intrinsic circadian mechanism located in the reproductive system ^56^. *Brunelleschi* encodes a protein that belongs to the TRAPII complex which is involved in vesicle trafficking in the secretory pathway ^57^. As male accessory glands are the secretory tissues of the reproductive system, *brunelleschi* is involved in the maturation of accessory glands in order to produce the secretory proteins of the seminal fluid. *Yellow-g* belongs to the MRJP/YELLOW family that includes the major royal jelly proteins and the yellow proteins. The *yellow* gene family has been associated with behavior ^58, 59, 60, 61, 62, 63^, pigmentation ^64, 65^, and sex-specific reproductive maturation ^62^ in *D. melanogaster* and *A. mellifera*.

*CG2254-like* encodes a dehydrogenase that is localized in the lipid droplets, organelles that store lipids and have a significant role in metabolism and membrane synthesis ^66^. The ejaculatory bulb is a muscle tissue and its contractions help to transfer the seminal fluid to the female flies during mating. During mating, the tissue has high energy demands and the presence of lipid droplets give them an alternative source of energy. Moreover, these lipid organelles could serve as a source of substrate for steroid hormone synthesis, such as ecdysteroid hormone that plays a significant role in reproduction.

With regard to the female reproductive tract, the expression profile of the selected genes was determined in DAY-7 old virgin females and at five time points (0, 3, 6, 9, 12, 24 h) after mating (Figure 6). This was based on the hypothesis that if a gene codes for a protein which is induced by mating, it should be expressed some time after mating. The obtained expression profiles of the 6 genes were variable. *Troponin C* showed limited expression after mating. *Ornithine decarboxylase antizyme* showed an increasing expression with the highest expression at 24 hours after mating, while *lingerer* (10-fold) and *bestrophin-2* (10-fold) showed highest expression at 12 hours. *Yolk protein-2* showed 2-fold overexpression 9 hours after mating and *glutathione S-transferase* showed highest expression immediately after mating (0 Hours).

**Figure 6:**
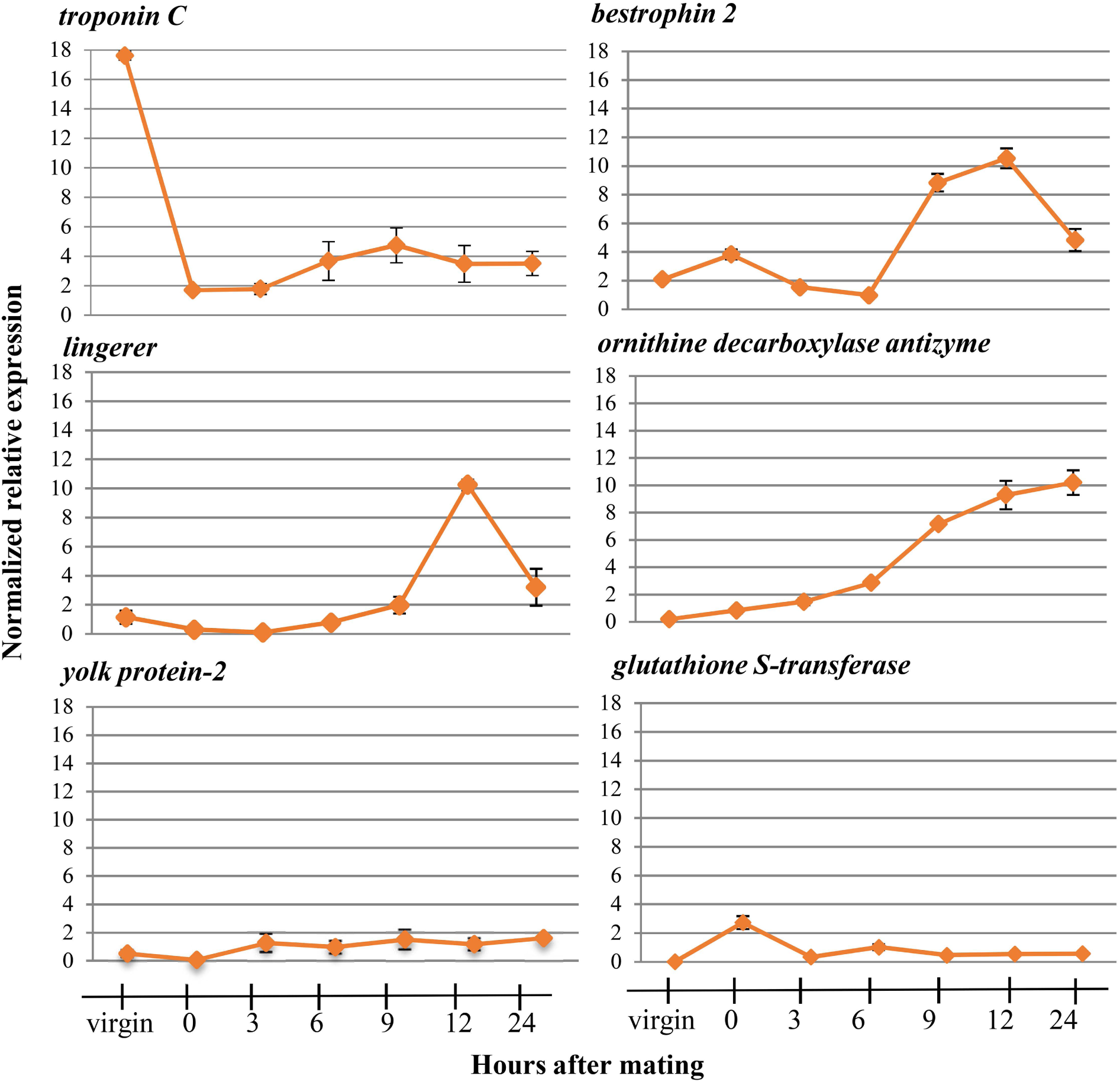
Expression profiles in female reproductive tissues. Expression profile of the selected genes from the virgin flies and several time points after mating (0, 3, 6, 9, 12, 24 hours). The error bars show the standard error of the mean between the three biological samples.

Troponin C protein plays a significant role in muscle contractions. In *Pieris rapae*, the small cabbage white butterfly, it was identified as a component of the bursa copulatrix female reproductive tissue that is responsible for the digestion of the nutrient-rich spermatophore produced by the male accessory glands ^67^. The overexpression of this gene in the female reproductive tract of virgin flies indicates its involvement in muscle contraction, probably aiding the digestion of the seminal fluid proteins that are transferred to the female during mating.

An upregulation of *Ornithine decarboxylase antizyme* (ODC-AZ) was observed in mated females. ODC-AZ binds and destabilizes the ornithine decarboxylase (ODC), a key enzyme in polyamine synthesis ^68^. Correlative changes between hormone levels and polyamine metabolism were described in several insects. For example, 20-hydroxyecdysone increases ODC activity in silk moth pupal tissues ^69^ and juvenile hormone stimulates ODC activity during vitellogenesis in *D*. *melanogaster* ^70^. Ornithine decarboxylase antizyme is an inhibitor of ODC. Inhibition of ODC activity causes impaired vitellogenesis in *Ae. aegypti* ^71^ and oviposition delay in the silkmoth, *Hyalophora cecropia* ^69^. The observed upregulation of their inhibitor indicates that ODC-AZ is probably involved in the control of ODCs levels in mated female olive flies.

*Lingerer* showed upregulation 12 hours after mating. Mutations of *lingerer* in male *D. melanogaster* result in abnormal matings and the “stuck” phenotype where males cannot be separated from females after the end of mating. It has been also identified as a maternal gene expressed in *D. melanogaster* early embryos ^72^. A similar expression profile has been demonstrated for the *bestrophin 2* gene. In *D. melanogaster* it encodes an oligomeric transmembrane protein that is thought to act as chloride channel ^73^. It facilitates the transportation of small molecules that are transferred into the female flies as part of the seminal fluid during mating.

*Yolk protein-2* homologue in *D. melanogaster* is expressed almost exclusively in females and it was associated with a female-sterile mutation ^74, 75^. *Yolk protein-2* gene encodes for a precursor of the major egg storage protein, the vitelline. There are three main factors that regulate vitellogenesis in *D. melanogaster*: a brain factor, an ovarian factor that stimulates fat body vitelline synthesis and a thoracic factor that is involved in the uptake of the vitelline by the ovaries ^76^. Moreover, vitellogenins have also been implicated in the transportation of various molecules such as sugars, lipids and hormones in insects ^77^.

Glutathione S-transferase epsilon class is a predicted intracellular or membrane-bound protein ^78^. Predicted intracellular proteins have been reported in the reproductive system of *D. melanogaster* ^4^, *A. mellifera* ^79^ and *Ae. aegypti* ^80^. For both *A. mellifera* and *Ae. aegypti* these proteins are suggested to be secreted through apocrine and holocrine secretion, non-standard secretion routes ^81, 82^. Macro apocrine secretion has been reported in the *B. oleae* male reproductive system ^36^.

The transcriptional profiles of the aforementioned genes identified in the reproductive tissues of *B. oleae* showed changes associated with the production of the proteins with relation to the sexual maturation of the males and the induction of several post-mating responses for the females. However, the very complex transcriptional profile of several of these genes necessitates further investigation. A key focus for future studies is a more detailed functional analysis and characterization of the processes that are involved in the reproduction of olive fly. Further experiments, including transient silencing through RNA interference (RNAi) of the aforementioned genes, would be helpful towards the clarification of the specific role of the genes in the mating procedure or the post-mating response of the female insects.

## 3. Conclusions

Undoubtedly, the reproduction system is one of the major systems of living organisms, primarily responsible for any species’ existence on earth. The present study sheds light on various aspects of the reproduction system of a major agricultural pest in an effort to understand its basic characteristics but also help in the development of specific and more ecologically sound approaches to insect control. During the last sixty years, agrochemical companies have been in a constant quest for all the more specific, effective and environmentally friendly approaches to contain populations of innocuous insects. Recently, RNA interference (RNAi) has been explored as a strategy for pest control by administering insect-targeted double-stranded RNA (dsRNA) to specifically block the expression of essential genes ^83, 84^. In that regard, inhibition of reproductive genes could lead to impairments in courtship behavior, fertility, oviposition, reproductive physiology or direct death of the developing embryo. Whatever the reason, the effect would be a dramatic drop in population size of the targeted insect. Alternatively, one can envision a CRISPR-based gene drive system in which a Cas9 (with its guide RNA) targets an essential reproductive gene ^85, 86^. The result would be a rapid substitution of the particular reproductive gene in a population with an impaired copy (or the complete deletion of the gene), thus damaging the reproductive ability of the pest and leading to its population collapse.

## 4. Methods

### 4.1 Ethics statement

For these experiments no specific permissions are required. This study did not interfere with any endangered or protected species as it was carried out on laboratory reared olive flies.

### 4.2 Fly culture and stock

The laboratory strain of the olive fly is part of the original stock from the Department of Biology, “Democritus” Nuclear Research Centre, Athens, Greece, and has been reared in our laboratory for over 20 years. The flies are reared at 25°C with a 12h light/12h dark photoperiod and humidity 65% in laboratory cages with diameter 30*30*30cm^3^ with wax cones inside for oviposition as described by Tzanakakis 1967 ^87^.

### 4.3 Tissue collection

To obtain the male adult flies used to generate the testes and accessory glands libraries as well as the female adult flies for the lower reproductive tract libraries, a standard laboratory rearing cage was set up with about 50 males and 50 females, one day after eclosion. The insects were maintained separately, in order to obtain tissues from two distinct groups:1. Virgin flies and 2. Mated flies. For the group of the virgin insects, males and females were removed from the cages 7 days after eclosion. For the group of the mated flies, we mixed virgin male and female flies to mate once. When the mating procedure was completed, the insects were kept in different cages separated for 12 hours before we perform dissections. The testes and MAGs with ejaculatory bulb from the male insects and the lower reproductive tract from the female insects were dissected in ddH2O. The dissected material was immediately immersed in the TRIzol Reagent (Ambion-Invitrogen). The RNA isolation was performed based on the TRIzol Reagent following the manufacturer’s instructions with slight modifications. An additional DNA removal using the TURBO DNA-free Kit (Ambion-Invitrogen), was performed, according to manufacturer’s instructions. RNA’s integrity was assessed by 1% agarose gel electrophoresis. The purity of all RNA samples was evaluated at Fleming Institute (Greece) with the use of (Agilent 2100 Bioanalyzer) and NanoDrop (2000).

For the validation of the differentially expressed genes, RNA was extracted from three pools each one containing 10 pairs of the lower reproductive tract (FAGs/ spermathecae, uterus), 10 pairs of testes and 10 pairs of ejaculatory bulb/ MAGs (three biological pool replicates) before and 12 hours after one mating of the aforementioned laboratory strain. To determine the expression profiles of the selected genes: 1. mRNA was extracted from 10 pairs of ejaculatory bulb/ MAGs from virgin males at DAY-0 (first day of eclosion) until DAY-7 and 2. RNA was extracted from three pools of 10 pairs of the lower reproductive tract from mated once females immediately after mating (0 hours) and 3 hours, 6 hours, 9 hours, 12 hours and 24 hours after mating for the female tissues. The RNA isolation was performed as described above.

### 4.4 RNA isolation for library preparation and functional analysis

mRNA transcripts from the samples were used to construct cDNA libraries for sequencing analysis. The cDNA libraries of accessory glands and ejaculatory bulb from virgin and mated male flies, the lower reproductive tract from virgin and mated female flies and the gut tissue from virgin male flies were sequenced on the Ion Proton™ system for Next-Generation. Sequencing was performed at the Fleming Institute (Greece) using the Ion Torrent™ Ion Chef™ automated platform. The cDNA library obtained from the testes of mated male insect (M_TESTES) was sequenced by Illumina Hi-Seq 2000 using the Illumina TruSeq RNA Sequencing protocol at the Genome Quebec in Canada.

### 4.5 Expression analyses of selected gene

Following extraction, the RNA was treated with 1.0 unit of DNase I (Invitrogen) according to manufacturer’s instructions. The total amount of DNA-free RNA obtained from each tissue was converted into cDNA using 300ng Random hexamer primers (equimolar mix of N5A, N5G, N5C and N5T), 200units MMLV Reverse Transcriptase (NEB), 5x Reaction buffer, 40mM dNTP mix and 40 units RNAse Inhibitor (NEB) according to the manufacturer’s instructions. Reverse transcription was conducted at 42°C for 50 minutes and 70°C for 15 minutes. The resulting cDNA was used in the followed qPCR reactions.

Specific primers to amplify genes identified on the transcriptomic analysis were designed by Primer-BLAST (http://www.ncbi.nlm.nih.gov/tools/primer-blast) (Table S5). Normalized relative quantitation was used to analyze changes in expression levels of the selected genes using a Real-time PCR approach. Expression values were calculated relatively to the housekeeping genes. *Rpl19* were used as reference in male reproductive tissues while *GAPDH* in FAGs/spermathecae ^41^.

The qRT-PCR conditions were: polymerase activation at 50 °C for 2 min, DNA denaturation step at 95 °C for 4 min, followed by 50 cycles of denaturation at 95 °C for 10 s, annealing/ extension and plate read at 55 °C for 20 s and finally, a step of melting curve analysis at a gradual increase of temperature over the range 55 °C to 95 °C. In this step, the detection of one gene-specific peak and the absence of primer dimer peaks were guaranteed. Each reaction was in a total volume of 15 μl, containing 5 μl from a 1:10 dilution of the cDNA template, 2X SYBR Select Master Mix (Applied Biosystem) and 300nM of each primer. The reactions were performed on a Bio-Rad Real-time thermal cycler CFX96 (Bio-Rad, Hercules, CA, USA) and data were analyzed through the CFX Manager™ software. All qRT-PCRs were performed in triplicate (i.e., three technical replicates).

### 4.6 Statistical tests

One-way ANOVA followed by Dunnett’s multiple comparisons test was performed using GraphPad Prism version 7.00 for Windows, GraphPad Software, La Jolla California USA, www.graphpad.com. Values were stated as mean ± standard error and a p value of < 0.05 was considered statistically significant.

## Supporting information

## Authors’ contributions

**Maria-Eleni Gregoriou**: Conceptualization; Methodology; Validation; Formal analysis; Resources; Data curation; Validation; Investigation. **Martin Reczko**: Data curation; Software; Formal analysis. **Konstantina Tsoumani:** Validation; Resources; Investigation. **Kostas Mathiopoulos:** Conceptualization; Supervision; Project administration; Funding Acquisition. All authors participated in drafting the manuscript and read and approved the final document.

## Acknowledgements

This research was implemented by the “ARISTEIA” Action of the “Operational programme Education and Life Long Learning” and the two postgraduate programs of the Department of Biochemistry and Biotechnology of the University of Thessaly (“Biotechnology – Quality Assessment in Nutrition and the Environment” and “Applications of Molecular Biology -Genetics –Diagnostic Biomarkers”).

## Supplementary material

Supplementary table S1: Genes differentially expressed in testes between virgin and mated males. Transcript_id, p value and fold values (for all the transcripts) and gene_id and annotation name (for the top100) are shown. Positive fold values indicate upregulation while negative fold values indicate downregulation in mated flies compared to virgin flies.

Supplementary table S2: Genes differentially expressed in male accessory glands and ejaculatory bulb between mature virgin and mated males. Transcript_id, p value and fold values (for all the transcripts) and gene_id and annotation name (for the top100) are shown. Negative fold values indicate upregulation in mated flies while positive fold values indicate downregulation compared to virgin flies.

Supplementary table S3: Genes differentially expressed in female lower reproductive tract between mature virgin and mated males. Transcript_id, p value and fold values (for all the transcripts) and gene_id and annotation name (for the top100) are shown. Negative fold values indicate upregulation in mated flies while positive fold values indicate downregulation compared to virgin flies.

Supplementary table S4: Annotated genes of *Bactrocera oleae* based on the homology of known seminal fluid proteins in *D. melanogaster* ^54^.

Supplementary table S5: Primers used for the qRT-PCR. The *RPL19* was used for the normalization of the values obtained from qRT-PCR for male tissues and *GAPDH* for the normalization of the values obtained from qRT-PCR for female tissues.

