## Supplementary material for "Decoding the reproductive system of the olive fruit fly, *Bactrocera oleae*"

| | | *B. oleae* transcriptome<br>Tissue: testes | | | |
|---|---|---|---|---|---|
| N | transcript_id | Annotation name | gene_id | logFC | PValue |
| 1 | c15699_g1 | *dikar* | NW_013581252.1.3 | 8.750 | 3.0E-07 |
| 2 | c582833_g1 | *not predicted* | | 8.639 | 5.4E-07 |
| 3 | c11986_g1 | *Octopamine receptor in mushroom bodies* | NW_013581214.1.89 | 8.422 | 1.7E-06 |
| 4 | c38051_g1 | *mucin-2* | NW_013581220.1.46 | 8.354 | 2.5E-06 |
| 5 | c52158_g4 | *not predicted* | | 8.318 | 2.9E-06 |
| 6 | c97454_g1 | *not predicted* | | 8.167 | 6.3E-06 |
| 7 | c57629_g2 | *not predicted* | | 8.041 | 1.2E-05 |
| 8 | c128061_g1 | *not predicted* | | 7.856 | 2.8E-05 |
| 9 | c52274_g1 | *not predicted* | | 7.699 | 5.8E-05 |
| 10 | c123143_g1 | *CG10911-like* | NW_013581220.1.59 | 7.699 | 5.8E-05 |
| 11 | c44387_g1 | *CG34189-like* | NW_013583061.1.3 | 7.629 | 1.1E-11 |
| 12 | c37552_g1 | *not predicted* | | 7.480 | 3.0E-11 |
| 13 | c56753_g2 | *Heat shock protein 23* | NW_013581228.1.43 | 7.323 | 2.9E-04 |
| 14 | c14215_g1 | *not predicted* | | 7.323 | 2.9E-04 |
| 15 | c13478_g1 | *antigen 5-related 2* | NW_013581440.1.14 | 7.172 | 5.3E-04 |
| 16 | c38273_g1 | *CG14958-like* | NW_013581493.1.7 | 7.057 | 7.0E-12 |
| 17 | c32508_g1 | *not predicted* | | 6.903 | 1.2E-09 |
| 18 | c47470_g1 | *not predicted* | | 6.849 | 1.6E-09 |
| 19 | c24782_g1 | *CG34426-like* | NW_013581493.1.9 | 6.776 | 1.9E-13 |
| 20 | c50402_g1 | *not predicted* | | 6.698 | 4.1E-09 |
| 21 | c36907_g1 | *CG16727-like* | NW_013581259.1.32 | 6.597 | 1.3E-10 |
| 22 | c45607_g1 | *CG9259-like* | NW_013581453.1.8 | 6.544 | 1.1E-08 |
| 23 | c84195_g1 | *CG2157-like* | NW_013581212.1.127 | 6.544 | 1.1E-08 |
| 24 | c44747_g1 | *CG6337-like* | NW_013582004.1.1 | 6.323 | 8.6E-11 |
| 25 | c51337_g1 | *not predicted* | | 6.308 | 4.4E-08 |
| 26 | c97069_g1 | *uncharacterized protein F12A10.7-like* | NW_013581215.1.52 | 6.191 | 7.9E-08 |
| 27 | c39724_g1 | *uncharacterized protein LOC106615417* | 6178_t | 6.104 | 2.2E-13 |
| 28 | c35561_g1 | *location of vulva defective 1* | NW_013581251.1.58 | 5.839 | 5.8E-07 |
| 29 | c25108_g1 | *uncharacterized protein LOC106615433* | 6172_t | 5.745 | 5.1E-12 |
| 30 | c122821_g1 | *vacuolar H[+] ATPase 100kD subunit 2* | NW_013583611.1.1 | 5.650 | 1.6E-06 |
| 31 | c34116_g1 | *not predicted* | | 5.603 | 2.2E-11 |
| 32 | c123047_g1 | *CG31789-like* | NW_013581220.1.134 | 5.599 | 2.1E-06 |
| 33 | c34524_g1 | *uncharacterized protein NW_013591147.1.1* | NW_013591147.1.1 | 5.584 | 7.0E-08 |
| 34 | c42518_g1 | *not predicted* | | 5.468 | 1.4E-07 |
| 35 | c122834_g1 | *uncharacterized protein LOC106615425* | 6177_t | 5.395 | 5.1E-11 |
| 36 | c15819_g1 | *uncharacterized protein LOC106615424* | NW_013581248.1.9 | 5.348 | 1.6E-11 |
| 37 | c55481_g1 | *not predicted* | | 5.202 | 5.8E-07 |
| 38 | c72383_g1 | *CG8560-like* | NW_013583355.1.2 | 5.109 | 1.6E-07 |
| 39 | c58415_g1 | *CG15043-like* | NW_013581513.1.17 | 5.094 | 2.0E-08 |
| 40 | c34152_g1 | *Cyp6a16* | NW_013583451.1.1 | 5.070 | 2.3E-08 |
| 41 | c15924_g1 | *CG31233-like* | NW_013582303.1.1 | 4.989 | 4.7E-05 |
| 42 | c48988_g3 | *uncharacterized protein LOC106615425* | NW_013581248.1.11 | 4.926 | 1.6E-08 |
| 43 | c54167_g1 | *CG15406-like* | NW_013583385.1.2 | 4.811 | 2.7E-07 |
| 44 | c39853_g1 | *not predicted* | | 4.709 | 1.2E-09 |
| 45 | c33022_g1 | *uncharacterized protein LOC106615431* | NW_013581248.1.10 | 4.682 | 2.4E-07 |
| 46 | c52085_g2 | *not predicted* | | 4.661 | 2.7E-07 |
| 47 | c48380_g1 | *not predicted* | | 4.589 | 2.1E-07 |
| 48 | c49725_g1 | *CG4363-like* | NW_013581215.1.132 | 4.572 | 1.7E-05 |
| 49 | c123043_g1 | *uncharacterized protein LOC106627291* | NW_013581220.1.47 | 4.477 | 3.1E-09 |
| 50 | c48988_g1 | *uncharacterized protein NW_013581248.1.14* | NW_013581248.1.14 | 4.468 | 7.1E-08 |
| 51 | c55436_g1 | *CG31106-like* | NW_013581235.1.12 | 4.392 | 4.0E-08 |
| 52 | c96078_g1 | *not predicted* | | 4.360 | 4.9E-05 |
| 53 | c53162_g1 | *not predicted* | | 4.327 | 5.8E-05 |
| 54 | c13906_g1 | *CG33282-like* | NW_013581214.1.54 | 4.258 | 4.2E-08 |
| 55 | c97206_g1 | *CG33998-like* | NW_013581220.1.132 | 4.247 | 4.3E-08 |
| 56 | c84109_g1 | *CG10650-like* | NW_013581251.1.59 | 4.201 | 2.2E-05 |
| 57 | c49026_g1 | *beta-site APP-cleaving enzyme* | NW_013581223.1.72 | 4.140 | 3.5E-07 |
| 58 | c33506_g1 | *acanthoscurrin-1-like* | ** NO NAME ASSIGNED ** | 4.110 | 1.6E-04 |
| 59 | c45977_g1 | *CG42235-like* | NW_013581576.1.7 | 4.091 | 1.3E-05 |
| 60 | c42936_g2 | *CG10031-like* | NW_013581471.1.3 | 4.086 | 7.0E-07 |
| 61 | c123354_g1 | *CG10911-like* | NW_013581220.1.59 | 3.983 | 7.6E-08 |
| 62 | c36293_g1 | *Cyp6a16* | NW_013583451.1.1 | 3.946 | 3.4E-04 |
| 63 | c48988_g2 | *uncharacterized protein LOC106615425* | NW_013581248.1.11 | 3.945 | 9.3E-07 |

| | | | | | |
|---|---|---|---|---|---|
| 64 | c112992_g1 | *CG3168-like* | NW_013583488.1.2 | 3.911 | 9.9E-07 |
| 65 | c55736_g1 | *CG30375-like* | 19027_t | 3.740 | 1.4E-06 |
| 66 | c109541_g1 | *CG15096-like* | NW_013581220.1.91 | 3.724 | 8.7E-05 |
| 67 | c50185_g1 | *SLC22A* | NW_013581465.1.4 | 3.698 | 5.6E-07 |
| 68 | c31533_g1 | *CG4363-like* | NW_013581215.1.131 | 3.694 | 5.0E-05 |
| 69 | c21199_g1 | *not predicted* | | 3.694 | 3.1E-06 |
| 70 | c43247_g1 | *not predicted* | | 3.606 | 2.3E-05 |
| 71 | c32887_g1 | *nucleolar protein 3-like* | NW_013581215.1.64 | 3.559 | 9.8E-05 |
| 72 | c46266_g1 | *black* | NW_013581211.1.19 | 3.528 | 2.2E-04 |
| 73 | c110791_g1 | *Cyp313a4* | NW_013581299.1.20 | 3.502 | 2.3E-06 |
| 74 | c49500_g1 | *tetraspanin 47F* | NW_013582559.1.4 | 3.497 | 2.6E-04 |
| 75 | c39986_g1 | *CG5246-like* | NW_013581294.1.10 | 3.482 | 3.3E-05 |
| 76 | c48959_g1 | *not predicted* | | 3.468 | 1.9E-06 |
| 77 | c55551_g1 | *not predicted* | | 3.369 | 4.6E-04 |
| 78 | c44647_g1 | *immune induced molecule 33* | NW_013581471.1.2 | 3.318 | 6.2E-05 |
| 79 | c111644_g1 | *CG13308-like* | NW_013581493.1.12 | 3.240 | 1.9E-04 |
| 80 | c44955_g1 | *CG5399-like* | NW_013581371.1.4 | 3.217 | 3.9E-05 |
| 81 | c27147_g1 | *Ecdysone-dependent gene 91* | NW_013581239.1.59 | 3.189 | 2.1E-05 |
| 82 | c49730_g1 | *urate oxidase* | NW_013581246.1.40 | 3.171 | 2.6E-05 |
| 83 | c54159_g1 | *CG33514-like* | NW_013581217.1.22 | 3.164 | 1.3E-05 |
| 84 | c41732_g1 | *serine-aspartate repeat-containing protein I-like* | NW_013581209.1.24 | 3.153 | 1.6E-04 |
| 85 | c48496_g1 | *CG9380-like* | NW_013583883.1.2 | 3.144 | 1.1E-05 |
| 86 | c43362_g1 | *not predicted* | | 3.130 | 8.4E-05 |
| 87 | c56485_g2 | *CG8303-like* | NW_013581233.1.1 | 3.127 | 4.7E-04 |
| 88 | c55628_g1 | *Neural Lazarillo* | NW_013581373.1.1 | 3.100 | 5.3E-04 |
| 89 | c26491_g1 | *uncharacterized protein LOC106624419* | NW_013582993.1.1 | 3.095 | 1.4E-05 |
| 90 | c42352_g1 | *CG8323-like* | NW_013581250.1.4 | 3.062 | 2.4E-05 |
| 91 | c59379_g1 | *alpha esterase-4* | NW_013581369.1.6 | 3.043 | 1.7E-04 |
| 92 | c52144_g1 | *not predicted* | | 3.043 | 1.7E-04 |
| 93 | c57220_g4 | *Ugt36Ba* | NW_013581212.1.175 | 3.024 | 4.6E-05 |
| 94 | c51861_g2 | *CG8834-like* | NW_013581215.1.3 | 2.965 | 7.6E-05 |
| 95 | c41816_g1 | *uncharacterized protein LOC106623971* | NW_013582629.1.1 | 2.956 | 7.0E-05 |
| 96 | c84264_g1 | *Sodium-dependent multivitamin transporter* | NW_013581377.1.15 | 2.924 | 2.1E-04 |
| 97 | c34087_g1 | *CG5096-like* | NW_013581248.1.32 | 2.885 | 4.1E-04 |
| 98 | c13969_g1 | not predicted | | 2.852 | 2.5E-04 |
| 99 | c39639_g1 | not predicted | | 2.791 | 1.0E-04 |
| 100 | c26571_g1 | not predicted | | 2.757 | 1.8E-04 |
| 101 | c29083_g1 | | | 2.688 | 1.21E-04 |
| 102 | c52914_g1 | | | 2.646 | 2.85E-04 |
| 103 | c58240_g1 | | | 2.589 | 3.92E-04 |
| 104 | c122942_g1 | | | 2.575 | 2.28E-04 |
| 105 | c19100_g1 | | | 2.474 | 4.05E-04 |
| 106 | c122679_g1 | | | 2.415 | 5.10E-04 |
| 107 | c50538_g1 | | | -2.387 | 5.45E-04 |
| 108 | c55344_g2 | | | -2.417 | 5.30E-04 |
| 109 | c57765_g3 | | | -2.436 | 4.28E-04 |
| 110 | c47407_g1 | | | -2.445 | 4.20E-04 |
| 111 | c56265_g1 | | | -2.457 | 4.00E-04 |
| 112 | c43732_g1 | | | -2.457 | 4.63E-04 |
| 113 | c51276_g1 | | | -2.479 | 3.89E-04 |
| 114 | c46996_g1 | | | -2.501 | 3.90E-04 |
| 115 | c55344_g1 | | | -2.520 | 2.94E-04 |
| 116 | c56371_g1 | | | -2.526 | 5.39E-04 |
| 117 | c44092_g1 | | | -2.553 | 2.93E-04 |
| 118 | c57759_g4 | | | -2.554 | 2.48E-04 |
| 119 | c55206_g1 | | | -2.559 | 2.99E-04 |
| 120 | c57355_g1 | | | -2.577 | 2.49E-04 |
| 121 | c31286_g1 | | | -2.593 | 2.42E-04 |
| 122 | c56249_g1 | | | -2.594 | 2.18E-04 |
| 123 | c56992_g5 | | | -2.615 | 5.39E-04 |
| 124 | c55353_g1 | | | -2.623 | 1.90E-04 |
| 125 | c57733_g1 | | | -2.624 | 1.65E-04 |
| 126 | c47427_g1 | | | -2.634 | 4.93E-04 |
| 127 | c57763_g1 | | | -2.637 | 2.53E-04 |

| | | | |
|---|---|---|---|
| 128 | c57312_g1 | -2.639 | 3.25E-04 |
| 129 | c53079_g2 | -2.644 | 3.99E-04 |
| 130 | c58068_g1 | -2.650 | 1.48E-04 |
| 131 | c110096_g1 | -2.654 | 4.65E-04 |
| 132 | c49218_g1 | -2.655 | 3.21E-04 |
| 133 | c45643_g1 | -2.656 | 2.10E-04 |
| 134 | c55193_g1 | -2.661 | 3.39E-04 |
| 135 | c43987_g1 | -2.667 | 4.57E-04 |
| 136 | c53267_g1 | -2.670 | 2.87E-04 |
| 137 | c52375_g1 | -2.673 | 1.34E-04 |
| 138 | c48857_g2 | -2.673 | 1.61E-04 |
| 139 | c56986_g1 | -2.686 | 2.76E-04 |
| 140 | c53096_g1 | -2.696 | 2.31E-04 |
| 141 | c84182_g1 | -2.699 | 4.56E-04 |
| 142 | c57475_g1 | -2.699 | 1.26E-04 |
| 143 | c49934_g1 | -2.716 | 2.47E-04 |
| 144 | c52618_g1 | -2.721 | 1.97E-04 |
| 145 | c56982_g1 | -2.730 | 1.23E-04 |
| 146 | c58058_g1 | -2.730 | 1.04E-04 |
| 147 | c53383_g1 | -2.733 | 1.59E-04 |
| 148 | c30480_g1 | -2.740 | 3.36E-04 |
| 149 | c56473_g1 | -2.745 | 9.11E-05 |
| 150 | c50698_g3 | -2.746 | 1.75E-04 |
| 151 | c57476_g1 | -2.758 | 4.93E-04 |
| 152 | c57488_g2 | -2.764 | 9.55E-05 |
| 153 | c41118_g1 | -2.764 | 1.69E-04 |
| 154 | c46296_g1 | -2.770 | 2.91E-04 |
| 155 | c55370_g2 | -2.771 | 1.03E-04 |
| 156 | c58044_g1 | -2.778 | 6.96E-05 |
| 157 | c54470_g1 | -2.786 | 7.27E-05 |
| 158 | c55938_g1 | -2.792 | 8.37E-05 |
| 159 | c97629_g1 | -2.794 | 1.75E-04 |
| 160 | c52929_g2 | -2.797 | 1.02E-04 |
| 161 | c52011_g1 | -2.809 | 2.19E-04 |
| 162 | c40049_g1 | -2.831 | 1.27E-04 |
| 163 | c47671_g1 | -2.840 | 1.14E-04 |
| 164 | c57095_g3 | -2.842 | 6.20E-05 |
| 165 | c33415_g1 | -2.844 | 1.92E-04 |
| 166 | c49275_g1 | -2.859 | 1.64E-04 |
| 167 | c46960_g1 | -2.860 | 5.69E-05 |
| 168 | c49571_g1 | -2.872 | 5.48E-05 |
| 169 | c40254_g1 | -2.884 | 4.97E-05 |
| 170 | c29046_g1 | -2.885 | 4.80E-05 |
| 171 | c57465_g1 | -2.887 | 5.38E-05 |
| 172 | c52535_g1 | -2.891 | 4.16E-05 |
| 173 | c85185_g1 | -2.900 | 3.80E-05 |
| 174 | c50803_g1 | -2.904 | 3.88E-05 |
| 175 | c55911_g2 | -2.904 | 3.04E-04 |
| 176 | c23092_g1 | -2.943 | 5.39E-05 |
| 177 | c74646_g1 | -2.963 | 2.29E-04 |
| 178 | c46348_g1 | -2.978 | 4.19E-05 |
| 179 | c57289_g3 | -2.996 | 1.42E-04 |
| 180 | c40895_g1 | -2.996 | 1.42E-04 |
| 181 | c54701_g1 | -2.997 | 3.91E-05 |
| 182 | c51200_g1 | -2.997 | 5.71E-05 |
| 183 | c48389_g2 | -3.012 | 2.00E-05 |
| 184 | c56712_g2 | -3.018 | 4.80E-05 |
| 185 | c50003_g1 | -3.018 | 1.55E-04 |
| 186 | c41212_g1 | -3.019 | 3.68E-04 |
| 187 | c27294_g1 | -3.040 | 2.03E-05 |
| 188 | c57161_g1 | -3.041 | 2.26E-05 |
| 189 | c53386_g2 | -3.074 | 3.20E-05 |
| 190 | c62696_g1 | -3.077 | 1.50E-04 |
| 191 | c55202_g1 | -3.081 | 3.36E-05 |
| 192 | c51547_g1 | -3.086 | 2.46E-05 |
| 193 | c57780_g1 | -3.106 | 3.13E-04 |

| | | | |
|---|---|---|---|
| 194 | c55013_g1 | -3.117 | 3.83E-05 |
| 195 | c56822_g1 | -3.118 | 3.98E-04 |
| 196 | c42551_g1 | -3.129 | 1.10E-05 |
| 197 | c57275_g2 | -3.141 | 3.37E-05 |
| 198 | c48497_g1 | -3.155 | 1.53E-05 |
| 199 | c10274_g1 | -3.155 | 7.01E-05 |
| 200 | c52211_g3 | -3.157 | 1.57E-05 |
| 201 | c54336_g3 | -3.160 | 9.91E-05 |
| 202 | c54965_g1 | -3.164 | 2.97E-05 |
| 203 | c64784_g1 | -3.187 | 4.16E-04 |
| 204 | c52365_g1 | -3.187 | 5.94E-05 |
| 205 | c122852_g1 | -3.187 | 2.97E-05 |
| 206 | c34646_g1 | -3.195 | 3.75E-05 |
| 207 | c39315_g1 | -3.200 | 2.90E-05 |
| 208 | c10933_g1 | -3.203 | 1.13E-04 |
| 209 | c39172_g1 | -3.216 | 8.00E-06 |
| 210 | c57699_g1 | -3.221 | 1.38E-04 |
| 211 | c97168_g1 | -3.229 | 1.90E-05 |
| 212 | c41861_g1 | -3.233 | 9.67E-05 |
| 213 | c40163_g1 | -3.238 | 3.28E-04 |
| 214 | c52020_g1 | -3.243 | 7.65E-05 |
| 215 | c53271_g1 | -3.257 | 4.90E-06 |
| 216 | c45672_g1 | -3.257 | 7.13E-05 |
| 217 | c28791_g1 | -3.263 | 2.91E-04 |
| 218 | c54949_g3 | -3.264 | 4.46E-05 |
| 219 | c54630_g1 | -3.289 | 3.49E-05 |
| 220 | c37731_g1 | -3.292 | 5.71E-06 |
| 221 | c57149_g1 | -3.297 | 8.14E-06 |
| 222 | c42373_g1 | -3.321 | 7.76E-05 |
| 223 | c52287_g1 | -3.323 | 6.17E-05 |
| 224 | c47613_g1 | -3.324 | 8.90E-06 |
| 225 | c52991_g1 | -3.327 | 4.00E-06 |
| 226 | c56631_g1 | -3.345 | 3.99E-06 |
| 227 | c44673_g1 | -3.349 | 1.14E-05 |
| 228 | c48961_g1 | -3.349 | 3.77E-05 |
| 229 | c44690_g1 | -3.351 | 5.34E-05 |
| 230 | c40912_g1 | -3.361 | 3.54E-05 |
| 231 | c52088_g1 | -3.369 | 5.34E-04 |
| 232 | c51738_g2 | -3.380 | 2.81E-06 |
| 233 | c57460_g1 | -3.385 | 3.47E-06 |
| 234 | c45148_g1 | -3.389 | 7.81E-05 |
| 235 | c58211_g1 | -3.396 | 1.12E-05 |
| 236 | c19115_g1 | -3.396 | 1.04E-04 |
| 237 | c50248_g1 | -3.405 | 1.48E-04 |
| 238 | c49755_g1 | -3.407 | 2.65E-06 |
| 239 | c47531_g1 | -3.412 | 2.33E-05 |
| 240 | c46178_g1 | -3.417 | 2.28E-04 |
| 241 | c40189_g1 | -3.421 | 1.73E-05 |
| 242 | c53900_g1 | -3.434 | 8.62E-05 |
| 243 | c44081_g1 | -3.436 | 3.95E-04 |
| 244 | c57523_g2 | -3.440 | 1.08E-05 |
| 245 | c57335_g3 | -3.443 | 9.15E-06 |
| 246 | c57768_g1 | -3.448 | 1.92E-06 |
| 247 | c33133_g1 | -3.468 | 2.63E-06 |
| 248 | c56821_g1 | -3.469 | 1.77E-06 |
| 249 | c56302_g1 | -3.493 | 6.94E-06 |
| 250 | c52329_g1 | -3.561 | 4.53E-05 |
| 251 | c48371_g1 | -3.564 | 8.97E-06 |
| 252 | c48474_g1 | -3.573 | 1.68E-05 |
| 253 | c37881_g1 | -3.578 | 3.05E-06 |
| 254 | c27193_g1 | -3.578 | 4.15E-05 |
| 255 | c46734_g1 | -3.587 | 4.81E-04 |
| 256 | c48397_g1 | -3.591 | 1.93E-04 |
| 257 | c44557_g1 | -3.594 | 9.80E-07 |
| 258 | c57187_g5 | -3.603 | 1.26E-06 |
| 259 | c40503_g1 | -3.608 | 9.48E-06 |

| | | | |
|---|---|---|---|
| 260 | c57459_g1 | -3.609 | 1.38E-05 |
| 261 | c57429_g1 | -3.626 | 7.35E-06 |
| 262 | c48199_g1 | -3.643 | 1.20E-06 |
| 263 | c48656_g1 | -3.644 | 6.66E-06 |
| 264 | c55380_g1 | -3.648 | 1.47E-04 |
| 265 | c57565_g3 | -3.648 | 1.47E-04 |
| 266 | c36326_g1 | -3.654 | 4.83E-06 |
| 267 | c46862_g1 | -3.654 | 4.83E-06 |
| 268 | c42390_g1 | -3.661 | 8.06E-07 |
| 269 | c42834_g1 | -3.661 | 4.63E-06 |
| 270 | c51291_g2 | -3.699 | 2.91E-04 |
| 271 | c45839_g1 | -3.699 | 2.91E-04 |
| 272 | c49521_g1 | -3.708 | 3.24E-05 |
| 273 | c54522_g2 | -3.721 | 3.31E-06 |
| 274 | c34264_g1 | -3.732 | 6.94E-07 |
| 275 | c56554_g1 | -3.734 | 1.10E-06 |
| 276 | c33419_g1 | -3.735 | 2.48E-04 |
| 277 | c55084_g2 | -3.749 | 3.29E-07 |
| 278 | c57026_g1 | -3.759 | 7.85E-06 |
| 279 | c44596_g1 | -3.775 | 4.02E-07 |
| 280 | c54063_g1 | -3.783 | 7.67E-05 |
| 281 | c63028_g1 | -3.787 | 1.27E-06 |
| 282 | c55915_g2 | -3.792 | 2.21E-06 |
| 283 | c26153_g1 | -3.803 | 1.81E-04 |
| 284 | c43509_g2 | -3.822 | 7.12E-07 |
| 285 | c53338_g1 | -3.833 | 5.98E-05 |
| 286 | c42248_g2 | -3.837 | 1.91E-07 |
| 287 | c57759_g1 | -3.862 | 1.19E-06 |
| 288 | c56175_g1 | -3.866 | 5.88E-06 |
| 289 | c38916_g1 | -3.869 | 1.33E-04 |
| 290 | c56951_g2 | -3.871 | 3.30E-07 |
| 291 | c57056_g1 | -3.875 | 2.26E-07 |
| 292 | c53374_g2 | -3.907 | 2.70E-06 |
| 293 | c54496_g1 | -3.911 | 4.21E-07 |
| 294 | c51614_g1 | -3.931 | 9.86E-05 |
| 295 | c53737_g1 | -3.964 | 6.35E-07 |
| 296 | c44883_g1 | -3.979 | 3.36E-04 |
| 297 | c22879_g1 | -3.979 | 3.36E-04 |
| 298 | c46130_g1 | -3.997 | 2.61E-05 |
| 299 | c84227_g1 | -4.003 | 1.04E-06 |
| 300 | c55216_g1 | -4.014 | 2.23E-07 |
| 301 | c29685_g1 | -4.019 | 2.33E-05 |
| 302 | c57856_g2 | -4.020 | 6.41E-05 |
| 303 | c40300_g1 | -4.022 | 2.78E-04 |
| 304 | c43713_g1 | -4.022 | 2.78E-04 |
| 305 | c56612_g1 | -4.022 | 2.78E-04 |
| 306 | c52848_g1 | -4.033 | 8.73E-07 |
| 307 | c51509_g2 | -4.041 | 2.08E-05 |
| 308 | c52465_g2 | -4.043 | 3.20E-06 |
| 309 | c55977_g3 | -4.053 | 1.18E-06 |
| 310 | c32079_g1 | -4.054 | 1.42E-07 |
| 311 | c28809_g1 | -4.056 | 2.99E-06 |
| 312 | c34934_g1 | -4.063 | 2.31E-04 |
| 313 | c58025_g1 | -4.076 | 1.35E-06 |
| 314 | c44543_g1 | -4.083 | 6.51E-07 |
| 315 | c51015_g1 | -4.088 | 7.66E-06 |
| 316 | c54053_g2 | -4.092 | 4.45E-07 |
| 317 | c51721_g1 | -4.104 | 1.92E-04 |
| 318 | c51754_g1 | -4.104 | 4.23E-05 |
| 319 | c56210_g2 | -4.104 | 4.23E-05 |
| 320 | c45019_g1 | -4.104 | 4.23E-05 |
| 321 | c56234_g1 | -4.107 | 1.18E-07 |
| 322 | c57830_g1 | -4.124 | 4.32E-07 |
| 323 | c57952_g1 | -4.128 | 5.19E-08 |
| 324 | c28857_g1 | -4.143 | 2.79E-07 |
| 325 | c51224_g1 | -4.143 | 9.46E-08 |

| | | | |
|---|---|---|---|
| 326 | c52025_g1 | -4.153 | 4.93E-06 |
| 327 | c56528_g1 | -4.155 | 1.27E-07 |
| 328 | c52729_g1 | -4.182 | 1.34E-04 |
| 329 | c53747_g1 | -4.183 | 2.84E-05 |
| 330 | c56928_g1 | -4.183 | 2.84E-05 |
| 331 | c71597_g1 | -4.185 | 4.15E-06 |
| 332 | c46777_g1 | -4.209 | 2.50E-05 |
| 333 | c52966_g1 | -4.274 | 1.41E-06 |
| 334 | c53631_g1 | -4.279 | 4.18E-07 |
| 335 | c20346_g1 | -4.283 | 1.71E-05 |
| 336 | c42212_g1 | -4.296 | 3.80E-07 |
| 337 | c54800_g1 | -4.308 | 4.88E-07 |
| 338 | c53638_g1 | -4.314 | 4.90E-06 |
| 339 | c42132_g1 | -4.316 | 1.14E-07 |
| 340 | c51579_g1 | -4.328 | 3.19E-08 |
| 341 | c109590_g1 | -4.359 | 5.78E-05 |
| 342 | c44734_g1 | -4.370 | 7.17E-08 |
| 343 | c56412_g1 | -4.375 | 3.27E-08 |
| 344 | c56056_g6 | -4.378 | 7.58E-09 |
| 345 | c33376_g1 | -4.398 | 9.39E-06 |
| 346 | c36801_g1 | -4.406 | 1.51E-07 |
| 347 | c44337_g1 | -4.425 | 4.20E-05 |
| 348 | c57307_g1 | -4.429 | 3.24E-08 |
| 349 | c54748_g1 | -4.431 | 1.02E-07 |
| 350 | c57630_g3 | -4.463 | 6.66E-06 |
| 351 | c39905_g1 | -4.484 | 5.95E-06 |
| 352 | c55854_g1 | -4.530 | 7.18E-08 |
| 353 | c58085_g1 | -4.544 | 3.63E-09 |
| 354 | c54804_g1 | -4.553 | 3.85E-04 |
| 355 | c57630_g5 | -4.553 | 3.85E-04 |
| 356 | c25218_g1 | -4.558 | 1.13E-07 |
| 357 | c52207_g1 | -4.563 | 2.69E-07 |
| 358 | c55977_g2 | -4.576 | 1.98E-05 |
| 359 | c47541_g1 | -4.589 | 9.87E-07 |
| 360 | c79516_g1 | -4.633 | 1.48E-05 |
| 361 | c55430_g1 | -4.633 | 1.48E-05 |
| 362 | c57146_g2 | -4.636 | 1.06E-07 |
| 363 | c54930_g1 | -4.642 | 1.70E-07 |
| 364 | c23816_g1 | -4.654 | 6.35E-08 |
| 365 | c22300_g1 | -4.660 | 2.44E-04 |
| 366 | c53649_g1 | -4.660 | 2.44E-04 |
| 367 | c39523_g1 | -4.660 | 1.29E-05 |
| 368 | c49786_g1 | -4.710 | 1.95E-04 |
| 369 | c51664_g1 | -4.710 | 1.95E-04 |
| 370 | c57146_g1 | -4.723 | 4.17E-08 |
| 371 | c46693_g1 | -4.735 | 1.26E-08 |
| 372 | c51676_g1 | -4.748 | 1.73E-07 |
| 373 | c56696_g1 | -4.748 | 1.73E-07 |
| 374 | c43721_g2 | -4.759 | 1.57E-04 |
| 375 | c41006_g1 | -4.759 | 1.57E-04 |
| 376 | c52071_g1 | -4.764 | 6.98E-10 |
| 377 | c57080_g3 | -4.790 | 6.56E-06 |
| 378 | c46897_g1 | -4.806 | 1.27E-04 |
| 379 | c36988_g1 | -4.806 | 1.27E-04 |
| 380 | c55953_g1 | -4.829 | 5.58E-08 |
| 381 | c52350_g1 | -4.896 | 8.44E-05 |
| 382 | c56933_g1 | -4.905 | 1.63E-07 |
| 383 | c50418_g1 | -4.939 | 6.91E-05 |
| 384 | c16849_g1 | -5.060 | 3.89E-05 |
| 385 | c56745_g4 | -5.064 | 5.85E-10 |
| 386 | c41322_g1 | -5.099 | 3.23E-05 |
| 387 | c36354_g1 | -5.099 | 3.23E-05 |
| 388 | c44172_g1 | -5.099 | 5.84E-10 |
| 389 | c49931_g1 | -5.103 | 5.06E-08 |
| 390 | c53943_g1 | -5.208 | 1.89E-05 |
| 391 | c51110_g1 | -5.310 | 1.14E-05 |

| | | | |
|---|---|---|---|
| 392 | c27665_g1 | -5.342 | 9.66E-06 |
| 393 | c22339_g1 | -5.342 | 9.66E-06 |
| 394 | c57729_g1 | -5.374 | 4.94E-10 |
| 395 | c36680_g1 | -5.404 | 7.02E-06 |
| 396 | c52158_g5 | -5.404 | 7.02E-06 |
| 397 | c58077_g1 | -5.414 | 1.69E-10 |
| 398 | c54336_g2 | -5.462 | 2.09E-08 |
| 399 | c53791_g1 | -5.464 | 5.15E-06 |
| 400 | c57138_g1 | -5.483 | 1.84E-08 |
| 401 | c51609_g1 | -5.543 | 1.38E-09 |
| 402 | c122664_g1 | -5.614 | 8.82E-10 |
| 403 | c97540_g1 | -5.732 | 1.24E-06 |
| 404 | c35555_g1 | -5.751 | 6.46E-13 |
| 405 | c49566_g1 | -5.752 | 7.32E-12 |
| 406 | c96830_g1 | -6.182 | 3.55E-14 |
| 407 | c53744_g1 | -6.603 | 7.70E-09 |
| 408 | c55702_g3 | -6.837 | 3.23E-11 |
| 409 | c56529_g2 | -7.095 | 3.80E-10 |
| 410 | c56969_g1 | -7.188 | 5.30E-04 |
| 411 | c50940_g1 | -7.188 | 5.30E-04 |
| 412 | c39646_g1 | -7.266 | 3.91E-04 |
| 413 | c54754_g1 | -7.266 | 3.91E-04 |
| 414 | c37426_g1 | -7.266 | 3.91E-04 |
| 415 | c50234_g1 | -7.266 | 3.91E-04 |
| 416 | c49497_g1 | -7.266 | 3.91E-04 |
| 417 | c127908_g1 | -7.339 | 2.91E-04 |
| 418 | c57032_g1 | -7.409 | 2.18E-04 |
| 419 | c45809_g1 | -7.409 | 2.18E-04 |
| 420 | c87715_g1 | -7.409 | 2.18E-04 |
| 421 | c25757_g1 | -7.476 | 1.65E-04 |
| 422 | c36221_g1 | -7.476 | 1.65E-04 |
| 423 | c56239_g1 | -7.476 | 1.65E-04 |
| 424 | c43357_g1 | -7.476 | 1.65E-04 |
| 425 | c23167_g1 | -7.540 | 1.26E-04 |
| 426 | c30197_g1 | -7.540 | 1.26E-04 |
| 427 | c54896_g1 | -7.601 | 9.64E-05 |
| 428 | c58091_g1 | -7.770 | 4.53E-05 |
| 429 | c14610_g1 | -7.770 | 4.53E-05 |
| 430 | c61895_g1 | -7.822 | 3.57E-05 |
| 431 | c129997_g1 | -7.822 | 3.57E-05 |
| 432 | c11533_g1 | -7.822 | 3.57E-05 |
| 433 | c44875_g1 | -7.873 | 2.82E-05 |
| 434 | c47892_g7 | -7.873 | 2.82E-05 |
| 435 | c26015_g1 | -7.921 | 2.25E-05 |
| 436 | c46922_g1 | -7.921 | 2.25E-05 |
| 437 | c15769_g1 | -7.921 | 2.25E-05 |
| 438 | c48096_g1 | -8.143 | 7.72E-06 |
| 439 | c47807_g1 | -8.183 | 6.32E-06 |
| 440 | c32888_g1 | -8.223 | 5.19E-06 |
| 441 | c55216_g2 | -8.261 | 4.28E-06 |
| 442 | c43606_g1 | -8.405 | 2.05E-06 |
| 443 | c53237_g1 | -8.472 | 1.45E-06 |
| 444 | c37751_g1 | -8.793 | 2.57E-07 |
| 445 | c39173_g1 | -8.870 | 1.68E-07 |
| 446 | c56973_g2 | -8.918 | 1.28E-07 |
| 447 | c61964_g1 | -8.966 | 9.84E-08 |
| 448 | c111150_g1 | -9.502 | 3.94E-09 |
| 449 | c53286_g1 | -9.683 | 1.34E-09 |
| 450 | c52187_g1 | -9.724 | 1.04E-09 |
