## Supplementary material for "Decoding the reproductive system of the olive fruit fly, *Bactrocera oleae*"

| | | *B. oleae* transcriptome<br>Tissue: male accessory glands, ejaculatory bulb | | | |
|---|---|---|---|---|---|
| N | transcript_id | Annotation name | gene_id | logFC | PValue |
| 1 | c31616_g1 | *attacin-A* | NW_013581217.1.68 | -13.051 | 1.03E-16 |
| 2 | c52655_g1 | *CG2254-like* | NW_013581268.1.20 | -12.304 | 1.58E-14 |
| 3 | c51710_g1 | *CG31729-like* | NW_013581210.1.33 | -12.110 | 5.89E-14 |
| 4 | c47341_g2 | *CG31798-like* | NW_013581459.1.12 | -11.860 | 3.11E-13 |
| 5 | c47596_g1 | *CG10096-like* | NW_013581245.1.66 | -11.831 | 3.79E-13 |
| 6 | c52892_g1 | *CG10435-like* | NW_013581672.1.3 | -11.801 | 4.63E-13 |
| 7 | c57023_g2 | *timeless* | NW_013581222.1.24 | -11.708 | 8.58E-13 |
| 8 | c23397_g1 | *CG4666-like* | NW_013581218.1.116 | -11.707 | 8.58E-13 |
| 9 | c47442_g1 | *sorting nexin 3* | NW_013581987.1.5 | -11.700 | 8.94E-13 |
| 10 | c53574_g1 | *CG7840-like* | NW_013588004.1.1 | -11.667 | 1.11E-12 |
| 11 | c40374_g1 | *catalase* | NW_013581552.1.5 | -11.645 | 1.30E-12 |
| 12 | c47533_g1 | *Nuclear protein localization 4* | NW_013585174.1.1 | -11.578 | 2.03E-12 |
| 13 | c55275_g2 | *CG43693-like* | NW_013581324.1.28 | -11.254 | 1.68E-11 |
| 14 | c41928_g1 | *mus81* | NW_013581254.1.18 | -11.230 | 1.94E-11 |
| 15 | c45555_g1 | *CG3690-like* | NW_013584662.1.1 | -11.191 | 2.43E-11 |
| 16 | c57217_g1 | *twenty four* | NW_013581628.1.11 | -11.185 | 2.62E-11 |
| 17 | c84745_g1 | *Nuclear polyadenosine RNA-binding 2-RA* | NW_013581231.1.68 | -11.180 | 2.72E-11 |
| 18 | c57024_g2 | *Na+/H+ hydrogen exchanger 2* | NW_013581246.1.67 | -11.153 | 3.24E-11 |
| 19 | c53204_g1 | *CG3394-like* | NW_013581407.1.1 | -11.101 | 4.53E-11 |
| 20 | c53812_g1 | *not predicted* | *not predicted* | -11.098 | 4.63E-11 |
| 21 | c57131_g1 | *CG34408-like* | NW_013582394.1.4 | -11.024 | 7.44E-11 |
| 22 | c39257_g2 | *CG3420-like* | NW_013581226.1.64 | -11.006 | 8.27E-11 |
| 23 | c10660_g1 | *Tetraspanin 42Ee* | NW_013581220.1.29 | -10.976 | 1.00E-10 |
| 24 | c47311_g1 | *Gustatory receptor 32a* | NW_013581242.1.18 | -10.965 | 1.07E-10 |
| 25 | c72530_g1 | | *not predicted* | -10.960 | 1.12E-10 |
| 26 | c53055_g1 | *Isoleucyl-tRNA synthetase* | NW_013581248.1.21 | -10.948 | 1.22E-10 |
| 27 | c55859_g1 | *galaktokinase* | NW_013583755.1.2 | -10.922 | 1.43E-10 |
| 28 | c39648_g1 | *CG7322-like* | NW_013581236.1.21 | -10.914 | 1.50E-10 |
| 29 | c57875_g1 | | *not predicted* | -10.907 | 1.57E-10 |
| 30 | c53746_g1 | *Imaginal disc growth factor 3-RB* | NW_013581222.1.28 | -10.849 | 2.27E-10 |
| 31 | c58086_g1 | *domino* | 13374_t | -10.834 | 2.50E-10 |
| 32 | c54574_g1 | *juvenile hormone epoxide hydrolase 2* | NW_013581209.1.139 | -10.788 | 3.34E-10 |
| 33 | c43349_g1 | c43349_g1 | 18614_t | -10.772 | 3.69E-10 |
| 34 | c55965_g3 | *Dopamine/Ecdysteroid receptor-RA* | NW_013581302.1.11 | -10.762 | 3.97E-10 |
| 35 | c54053_g5 | *Signal-transducer and activator of transcription protein at 92E* | NW_013581294.1.27 | -10.746 | 4.39E-10 |
| 36 | c52465_g1 | *Guanine nucleotide exchange factor in mesoderm* | NW_013581692.1.2 | -10.683 | 6.46E-10 |
| 37 | c55810_g1 | *CG3376-like* | NW_013581229.1.55 | -10.667 | 7.18E-10 |
| 38 | c33324_g1 | *Small ribonucleoprotein particle protein SmB* | NW_013581247.1.17 | -10.650 | 7.99E-10 |
| 39 | c54844_g2 | *Glucose transporter 1* | NW_013581411.1.14 | -10.645 | 8.21E-10 |
| 40 | c26099_g1 | *CG30008-LIKE* | NW_013581852.1.2 | -10.644 | 8.43E-10 |
| 41 | c58149_g1 | *expanded* | NW_013581242.1.89 | -10.630 | 9.14E-10 |
| 42 | c51872_g1 | *CG43143-like* | NW_013582061.1.2 | -10.620 | 9.66E-10 |
| 43 | c30784_g2 | *Methylthioadenosine phosphorylase* | NW_013581238.1.9 | -10.593 | 1.14E-09 |
| 44 | c10274_g1 | *echinoid* | NW_013581222.1.2 | -10.574 | 1.31E-09 |
| 45 | c56997_g1 | *Gustatory receptor 21a* | NW_013584380.1.1 | -10.563 | 1.39E-09 |
| 46 | c32538_g1 | *CG12321-like* | NW_013581448.1.7 | -10.544 | 1.55E-09 |
| 47 | c54799_g1 | *Cysteine string protein* | NW_013581269.1.4 | -10.532 | 1.69E-09 |
| 48 | c57176_g2 | *wide awake* | NW_013581262.1.13 | -10.527 | 1.74E-09 |
| 49 | c43839_g1 | *Gamma-interferon-inducible reductase 1* | NW_013581310.1.15 | -10.525 | 1.74E-09 |
| 50 | c52007_g1 | *Pyruvate carboxylase* | NW_013581314.1.23 | -10.520 | 1.85E-09 |
| 51 | c57583_g1 | *Abl tyrosine kinase* | NW_013581213.1.160 | -10.516 | 1.85E-09 |
| 52 | c54018_g2 | *CG17746-like* | NW_013582369.1.3 | -10.500 | 2.08E-09 |
| 53 | c54728_g1 | *yellow-g* | NW_013581221.1.69 | -10.490 | 2.14E-09 |
| 54 | c49066_g1 | *CG4757-like* | 17950_t | -10.481 | 2.34E-09 |
| 55 | c45487_g1 | *CG5590-like* | NW_013581593.1.5 | -10.480 | 2.34E-09 |
| 56 | c47712_g1 | *astray* | NW_013581245.1.45 | -10.475 | 2.41E-09 |
| 57 | c52931_g1 | *CG9747-like* | NW_013581266.1.30 | -10.454 | 2.80E-09 |
| 58 | c49237_g1 | *CG4266-like* | NW_013581233.1.39 | -10.379 | 4.33E-09 |
| 59 | c34297_g1 | *CG7185-like* | NW_013581249.1.17 | -10.372 | 4.61E-09 |
| 60 | c57747_g1 | *CG1024-like* | NW_013581280.1.32 | -10.353 | 5.07E-09 |
| 61 | c56234_g1 | *split ends* | NW_013581242.1.94 | -10.352 | 5.24E-09 |
| 62 | c53020_g1 | *CG8243-like* | NW_013581233.1.54 | -10.341 | 5.41E-09 |
| 63 | c56726_g1 | *PTEN-induced putative kinase 1* | NW_013581231.1.28 | -10.275 | 8.33E-09 |
| 64 | c57660_g1 | | *not predicted* | -10.269 | 8.62E-09 |
| 65 | c50703_g1 | *CG40160-like* | NW_013582263.1.2 | -10.253 | 9.55E-09 |

| | | | | | |
|---|---|---|---|---|---|
| 66 | c50574_g1 | *CG12121-like* | NW_013582327.1.2 | -10.229 | 1.10E-08 |
| 67 | c54908_g1 | *CG8858-like* | NW_013581229.1.3 | -10.222 | 1.13E-08 |
| 68 | c55183_g1 | *CG16935-like* | NW_013582071.1.6 | -10.211 | 1.22E-08 |
| 69 | c55990_g1 | *ance-4-ra* | NW_013583092.1.3 | -10.197 | 1.35E-08 |
| 70 | c56768_g6 | *CG1090-like* | NW_013582064.1.1 | -10.193 | 1.35E-08 |
| 71 | c46700_g1 | *Hexosaminidase 1* | NW_013581275.1.12 | -10.187 | 1.40E-08 |
| 72 | c34700_g1 | *Smad anchor for receptor activation* | NW_013581209.1.20 | -10.171 | 1.56E-08 |
| 73 | c56128_g2 | *moody* | NW_013582266.1.2 | -10.162 | 1.62E-08 |
| 74 | c54701_g1 | *strawberry notch* | NW_013581414.1.4 | -10.148 | 1.81E-08 |
| 75 | c47569_g1 | *CG11414-like* | NW_013581215.1.108 | -10.135 | 1.94E-08 |
| 76 | c55175_g1 | *Vacuolar protein sorting 24* | NW_013581374.1.3 | -10.133 | 1.94E-08 |
| 77 | c52369_g1 | *CG8888-like* | NW_013581233.1.61 | -10.114 | 2.17E-08 |
| 78 | c48551_g1 | *Nucleoporin 214kD* | NW_013581215.1.31 | -10.098 | 2.43E-08 |
| 79 | c71420_g1 | *Phosphoenolpyruvate carboxykinase* | NW_013581568.1.4 | -10.082 | 7.02E-09 |
| 80 | c53306_g2 | *ldlCp-related protein* | NW_013582073.1.8 | -10.079 | 7.02E-09 |
| 81 | c51656_g1 | *CG5390-like* | NW_013581211.1.51 | -10.069 | 7.02E-09 |
| 82 | c37891_g1 | *Heat shock protein cognate 20* | NW_013581285.1.42 | -10.067 | 7.58E-09 |
| 83 | c55308_g1 | *odd skipped* | NW_013581667.1.1 | -10.065 | 7.58E-09 |
| 84 | c54308_g1 | *CG1208-like* | NW_013581589.1.12 | -10.054 | 8.20E-09 |
| 85 | c46027_g1 | *CG8417-like* | NW_013583228.1.2 | -10.047 | 8.52E-09 |
| 86 | c53094_g7 | *Acid phosphatase 1* | NW_013581380.1.16 | -10.046 | 8.52E-09 |
| 87 | c56461_g1 | *elbow B* | NW_013583577.1.1 | -10.013 | 1.04E-08 |
| 88 | c47034_g1 | *CG5254-like* | NW_013581478.1.4 | -10.009 | 1.08E-08 |
| 89 | c56555_g3 | *GluClα* | NW_013581276.1.14 | -10.008 | 1.08E-08 |
| 90 | c56397_g3 | *Ryanodine receptor* | NW_013581420.1.2 | -10.007 | 1.08E-08 |
| 91 | c57800_g1 | *heixuedian* | NW_013581242.1.70 | -10.002 | 1.13E-08 |
| 92 | c10628_g1 | *CG4332-like* | NW_013583288.1.2 | -9.996 | 1.17E-08 |
| 93 | c36907_g1 | *CG16727-like* | NW_013581259.1.32 | -9.994 | 1.08E-08 |
| 94 | c49121_g2 | *Microcephalin* | NW_013581448.1.9 | -9.994 | 1.17E-08 |
| 95 | c55037_g1 | | *not predicted* | -9.987 | 1.22E-08 |
| 96 | c43052_g1 | *CG7011-like* | NW_013581241.1.24 | -9.972 | 1.33E-08 |
| 97 | c51676_g1 | *SP2637-RC like* | NW_013581226.1.57 | -9.972 | 1.33E-08 |
| 98 | c46198_g1 | *CG4603-like* | NW_013581228.1.2 | -9.971 | 1.33E-08 |
| 99 | c23655_g1 | *CG17221-like* | NW_013581662.1.6 | -9.968 | 1.38E-08 |
| 100 | c55935_g1 | *CG4896-like* | NW_013581251.1.13 | -9.959 | 1.44E-08 |
| 101 | c47358_g2 | | | -9.947 | 1.56E-08 |
| 102 | c57740_g4 | | | -9.936 | 1.70E-08 |
| 103 | c84178_g1 | | | -9.911 | 1.85E-08 |
| 104 | c24730_g1 | | | -9.895 | 2.20E-08 |
| 105 | c49709_g1 | | | -9.893 | 2.20E-08 |
| 106 | c54174_g1 | | | -9.889 | 2.30E-08 |
| 107 | c47367_g1 | | | -9.888 | 2.20E-08 |
| 108 | c36529_g1 | | | -9.884 | 2.30E-08 |
| 109 | c109723_g1 | | | -9.883 | 2.30E-08 |
| 110 | c39363_g1 | | | -9.879 | 2.40E-08 |
| 111 | c44810_g1 | | | -9.871 | 2.50E-08 |
| 112 | c48513_g1 | | | -9.868 | 2.40E-08 |
| 113 | c53513_g1 | | | -9.864 | 2.62E-08 |
| 114 | c53176_g1 | | | -9.845 | 2.86E-08 |
| 115 | c57054_g1 | | | -9.842 | 2.99E-08 |
| 116 | c42361_g1 | | | -9.838 | 2.99E-08 |
| 117 | c57559_g1 | | | -9.822 | 3.27E-08 |
| 118 | c42395_g1 | | | -9.819 | 3.42E-08 |
| 119 | c56756_g1 | | | -9.818 | 3.42E-08 |
| 120 | c36028_g1 | | | -9.817 | 3.27E-08 |
| 121 | c43148_g1 | | | -9.812 | 3.42E-08 |
| 122 | c47769_g1 | | | -9.812 | 3.58E-08 |
| 123 | c56109_g1 | | | -9.807 | 3.58E-08 |
| 124 | c57289_g3 | | | -9.800 | 3.74E-08 |
| 125 | c55614_g1 | | | -9.798 | 3.74E-08 |
| 126 | c23039_g1 | | | -9.794 | 3.92E-08 |
| 127 | c58429_g1 | | | -9.785 | 4.10E-08 |
| 128 | c55049_g1 | | | -9.784 | 4.10E-08 |
| 129 | c54905_g1 | | | -9.781 | 4.30E-08 |
| 130 | c56979_g1 | | | -9.770 | 4.50E-08 |
| 131 | c46230_g1 | | | -9.766 | 4.71E-08 |
| 132 | c57287_g3 | | | -9.756 | 4.94E-08 |
| 133 | c57544_g1 | | | -9.751 | 4.94E-08 |
| 134 | c54779_g3 | | | -9.751 | 5.18E-08 |

| | | | |
|---|---|---|---|
| 135 | c50340_g1 | -9.746 | 5.18E-08 |
| 136 | c54327_g1 | -9.745 | 5.18E-08 |
| 137 | c49506_g1 | -9.743 | 5.43E-08 |
| 138 | c55988_g1 | -9.737 | 5.43E-08 |
| 139 | c43974_g1 | -9.735 | 5.43E-08 |
| 140 | c122920_g1 | -9.733 | 5.43E-08 |
| 141 | c53769_g1 | -9.722 | 5.97E-08 |
| 142 | c49458_g1 | -9.715 | 6.26E-08 |
| 143 | c56560_g1 | -9.710 | 6.57E-08 |
| 144 | c46120_g1 | -9.709 | 6.57E-08 |
| 145 | c51692_g1 | -9.709 | 6.57E-08 |
| 146 | c56087_g1 | -9.703 | 6.57E-08 |
| 147 | c56786_g4 | -9.700 | 6.89E-08 |
| 148 | c36776_g1 | -9.699 | 6.89E-08 |
| 149 | c52416_g1 | -9.696 | 6.89E-08 |
| 150 | c49617_g1 | -9.691 | 7.23E-08 |
| 151 | c53862_g1 | -9.688 | 7.23E-08 |
| 152 | c47841_g1 | -9.679 | 7.60E-08 |
| 153 | c49752_g1 | -9.674 | 7.98E-08 |
| 154 | c72383_g1 | -9.669 | 7.23E-08 |
| 155 | c54746_g5 | -9.668 | 8.38E-08 |
| 156 | c38329_g1 | -9.662 | 8.38E-08 |
| 157 | c36507_g1 | -9.659 | 8.81E-08 |
| 158 | c53032_g1 | -9.653 | 8.81E-08 |
| 159 | c47521_g1 | -9.647 | 8.81E-08 |
| 160 | c58154_g3 | -9.640 | 9.73E-08 |
| 161 | c53847_g1 | -9.626 | 1.08E-07 |
| 162 | c54326_g1 | -9.598 | 1.25E-07 |
| 163 | c57202_g1 | -9.595 | 1.25E-07 |
| 164 | c54309_g1 | -9.592 | 1.25E-07 |
| 165 | c55372_g1 | -9.591 | 1.25E-07 |
| 166 | c56198_g1 | -9.590 | 1.32E-07 |
| 167 | c57339_g1 | -9.583 | 1.39E-07 |
| 168 | c55488_g1 | -9.581 | 1.39E-07 |
| 169 | c29799_g1 | -9.581 | 1.39E-07 |
| 170 | c37516_g1 | -9.573 | 1.39E-07 |
| 171 | c54688_g2 | -9.568 | 1.46E-07 |
| 172 | c50533_g1 | -9.555 | 1.63E-07 |
| 173 | c56117_g3 | -9.547 | 1.63E-07 |
| 174 | c27746_g1 | -9.546 | 1.63E-07 |
| 175 | c71442_g1 | -9.542 | 1.71E-07 |
| 176 | c32929_g1 | -9.542 | 1.71E-07 |
| 177 | c57257_g1 | -9.529 | 1.81E-07 |
| 178 | c52857_g1 | -9.526 | 1.91E-07 |
| 179 | c71604_g1 | -9.525 | 1.91E-07 |
| 180 | c55825_g1 | -9.523 | 1.91E-07 |
| 181 | c25284_g1 | -9.519 | 1.91E-07 |
| 182 | c57571_g4 | -9.517 | 2.01E-07 |
| 183 | c51418_g1 | -9.511 | 2.12E-07 |
| 184 | c49047_g1 | -9.510 | 2.12E-07 |
| 185 | c24597_g1 | -9.509 | 2.01E-07 |
| 186 | c57828_g1 | -9.508 | 2.12E-07 |
| 187 | c55077_g1 | -9.507 | 2.12E-07 |
| 188 | c49465_g1 | -9.500 | 2.12E-07 |
| 189 | c10586_g1 | -9.492 | 2.24E-07 |
| 190 | c52312_g1 | -9.488 | 2.37E-07 |
| 191 | c57338_g3 | -9.487 | 2.37E-07 |
| 192 | c54951_g2 | -9.482 | 2.50E-07 |
| 193 | c54954_g1 | -9.478 | 2.50E-07 |
| 194 | c56282_g1 | -9.475 | 2.50E-07 |
| 195 | c52101_g1 | -9.470 | 2.65E-07 |
| 196 | c34968_g1 | -9.468 | 2.65E-07 |
| 197 | c56584_g5 | -9.451 | 2.80E-07 |
| 198 | c55513_g3 | -9.445 | 2.96E-07 |
| 199 | c54036_g1 | -9.444 | 2.96E-07 |
| 200 | c32455_g1 | -9.441 | 3.13E-07 |
| 201 | c46987_g1 | -9.439 | 3.13E-07 |
| 202 | c45548_g1 | -9.435 | 3.13E-07 |
| 203 | c39884_g1 | -9.419 | 3.31E-07 |

| | | | |
|---|---|---|---|
| 204 | c54070_g1 | -9.418 | 3.50E-07 |
| 205 | c51292_g1 | -9.410 | 3.50E-07 |
| 206 | c57127_g1 | -9.408 | 3.71E-07 |
| 207 | c49531_g1 | -9.405 | 3.71E-07 |
| 208 | c39796_g1 | -9.404 | 3.93E-07 |
| 209 | c44934_g1 | -9.398 | 3.93E-07 |
| 210 | c52661_g1 | -9.398 | 3.93E-07 |
| 211 | c53337_g1 | -9.396 | 3.93E-07 |
| 212 | c55043_g1 | -9.394 | 3.93E-07 |
| 213 | c54597_g1 | -9.390 | 4.16E-07 |
| 214 | c45909_g1 | -9.387 | 4.16E-07 |
| 215 | c10568_g1 | -9.386 | 4.16E-07 |
| 216 | c52914_g1 | -9.384 | 3.71E-07 |
| 217 | c58030_g1 | -9.378 | 4.41E-07 |
| 218 | c56800_g1 | -9.372 | 4.41E-07 |
| 219 | c34576_g1 | -9.372 | 4.41E-07 |
| 220 | c50975_g1 | -9.366 | 4.67E-07 |
| 221 | c53043_g1 | -9.352 | 5.26E-07 |
| 222 | c55223_g1 | -9.350 | 5.26E-07 |
| 223 | c51209_g1 | -9.348 | 5.26E-07 |
| 224 | c58938_g1 | -9.345 | 5.26E-07 |
| 225 | c48323_g1 | -9.344 | 5.26E-07 |
| 226 | c45033_g1 | -9.339 | 5.58E-07 |
| 227 | c32761_g1 | -9.337 | 5.58E-07 |
| 228 | c41198_g1 | -9.335 | 5.58E-07 |
| 229 | c55117_g1 | -9.333 | 5.58E-07 |
| 230 | c53527_g1 | -9.333 | 5.58E-07 |
| 231 | c50085_g1 | -9.332 | 5.58E-07 |
| 232 | c53840_g1 | -9.328 | 5.92E-07 |
| 233 | c44418_g1 | -9.324 | 5.92E-07 |
| 234 | c57220_g4 | -9.323 | 5.58E-07 |
| 235 | c51512_g1 | -9.321 | 5.92E-07 |
| 236 | c51138_g1 | -9.317 | 6.29E-07 |
| 237 | c48978_g1 | -9.309 | 6.68E-07 |
| 238 | c43752_g1 | -9.306 | 6.68E-07 |
| 239 | c53370_g1 | -9.304 | 6.68E-07 |
| 240 | c50866_g1 | -9.304 | 6.68E-07 |
| 241 | c48591_g1 | -9.296 | 7.10E-07 |
| 242 | c46050_g1 | -9.296 | 6.68E-07 |
| 243 | c54493_g2 | -9.294 | 7.10E-07 |
| 244 | c54940_g1 | -9.287 | 7.10E-07 |
| 245 | c27232_g1 | -9.285 | 7.55E-07 |
| 246 | c47829_g2 | -9.283 | 7.55E-07 |
| 247 | c55348_g1 | -9.281 | 7.55E-07 |
| 248 | c46703_g1 | -9.276 | 8.03E-07 |
| 249 | c55949_g1 | -9.275 | 7.55E-07 |
| 250 | c57701_g1 | -9.272 | 8.03E-07 |
| 251 | c44049_g1 | -9.264 | 8.55E-07 |
| 252 | c57848_g1 | -9.262 | 8.55E-07 |
| 253 | c57745_g1 | -9.261 | 8.55E-07 |
| 254 | c40654_g1 | -9.260 | 8.55E-07 |
| 255 | c51136_g1 | -9.256 | 8.55E-07 |
| 256 | c42808_g1 | -9.256 | 8.55E-07 |
| 257 | c49462_g1 | -9.251 | 9.10E-07 |
| 258 | c54688_g1 | -9.248 | 9.10E-07 |
| 259 | c71410_g1 | -9.248 | 8.55E-07 |
| 260 | c58230_g1 | -9.241 | 9.69E-07 |
| 261 | c42127_g1 | -9.237 | 9.69E-07 |
| 262 | c57784_g1 | -9.236 | 9.69E-07 |
| 263 | c41973_g1 | -9.233 | 9.69E-07 |
| 264 | c52946_g1 | -9.233 | 9.69E-07 |
| 265 | c13906_g1 | -9.227 | 9.10E-07 |
| 266 | c54049_g1 | -9.227 | 1.03E-06 |
| 267 | c56909_g3 | -9.223 | 1.03E-06 |
| 268 | c56302_g1 | -9.221 | 1.03E-06 |
| 269 | c26755_g1 | -9.221 | 1.03E-06 |
| 270 | c47182_g1 | -9.216 | 1.03E-06 |
| 271 | c50571_g1 | -9.207 | 1.10E-06 |
| 272 | c57411_g1 | -9.206 | 1.17E-06 |

| | | | |
|---|---|---|---|
| 273 | c55732_g1 | -9.205 | 1.17E-06 |
| 274 | c48948_g1 | -9.203 | 1.17E-06 |
| 275 | c55472_g1 | -9.201 | 1.17E-06 |
| 276 | c47914_g1 | -9.200 | 1.17E-06 |
| 277 | c38251_g1 | -9.197 | 1.17E-06 |
| 278 | c56986_g2 | -9.196 | 1.25E-06 |
| 279 | c53439_g1 | -9.194 | 1.25E-06 |
| 280 | c25322_g1 | -9.187 | 1.25E-06 |
| 281 | c54951_g1 | -9.187 | 1.25E-06 |
| 282 | c109602_g1 | -9.185 | 1.25E-06 |
| 283 | c47291_g1 | -9.183 | 1.33E-06 |
| 284 | c56676_g1 | -9.181 | 1.33E-06 |
| 285 | c34861_g1 | -9.181 | 1.33E-06 |
| 286 | c55357_g1 | -9.175 | 1.33E-06 |
| 287 | c53808_g2 | -9.174 | 1.42E-06 |
| 288 | c52669_g1 | -9.170 | 1.33E-06 |
| 289 | c122989_g1 | -9.169 | 1.42E-06 |
| 290 | c55220_g3 | -9.169 | 1.42E-06 |
| 291 | c44767_g1 | -9.166 | 1.42E-06 |
| 292 | c48858_g1 | -9.165 | 1.42E-06 |
| 293 | c57233_g1 | -9.162 | 1.52E-06 |
| 294 | c57683_g1 | -9.153 | 1.52E-06 |
| 295 | c56138_g2 | -9.147 | 1.62E-06 |
| 296 | c54430_g1 | -9.145 | 1.62E-06 |
| 297 | c36286_g1 | -9.137 | 1.62E-06 |
| 298 | c39441_g1 | -9.135 | 1.74E-06 |
| 299 | c48399_g1 | -9.127 | 1.74E-06 |
| 300 | c51847_g1 | -9.127 | 1.74E-06 |
| 301 | c53384_g1 | -9.122 | 1.74E-06 |
| 302 | c50336_g1 | -9.121 | 1.74E-06 |
| 303 | c54144_g1 | -9.121 | 1.74E-06 |
| 304 | c52447_g2 | -9.118 | 1.62E-06 |
| 305 | c32480_g1 | -9.118 | 1.86E-06 |
| 306 | c49755_g1 | -9.115 | 1.99E-06 |
| 307 | c54339_g3 | -9.114 | 1.99E-06 |
| 308 | c55069_g1 | -9.105 | 1.99E-06 |
| 309 | c55735_g1 | -9.105 | 1.99E-06 |
| 310 | c47662_g1 | -9.103 | 2.12E-06 |
| 311 | c33943_g1 | -9.100 | 2.12E-06 |
| 312 | c44704_g1 | -9.093 | 2.12E-06 |
| 313 | c51421_g1 | -9.086 | 2.28E-06 |
| 314 | c49443_g1 | -9.085 | 2.28E-06 |
| 315 | c52988_g1 | -9.080 | 2.44E-06 |
| 316 | c55362_g1 | -9.071 | 2.44E-06 |
| 317 | c57651_g2 | -9.070 | 2.44E-06 |
| 318 | c57826_g1 | -9.066 | 2.44E-06 |
| 319 | c47425_g1 | -9.062 | 2.61E-06 |
| 320 | c47867_g1 | -9.056 | 2.61E-06 |
| 321 | c55125_g1 | -9.055 | 2.61E-06 |
| 322 | c54324_g2 | -9.052 | 2.61E-06 |
| 323 | c57070_g1 | -9.049 | 2.80E-06 |
| 324 | c57734_g1 | -9.049 | 2.80E-06 |
| 325 | c43968_g1 | -9.046 | 2.80E-06 |
| 326 | c45007_g1 | -9.045 | 2.61E-06 |
| 327 | c35166_g1 | -9.041 | 2.80E-06 |
| 328 | c51973_g1 | -9.037 | 3.00E-06 |
| 329 | c48797_g1 | -9.032 | 3.00E-06 |
| 330 | c49237_g2 | -9.032 | 3.00E-06 |
| 331 | c56548_g1 | -9.026 | 3.00E-06 |
| 332 | c84119_g1 | -9.016 | 3.22E-06 |
| 333 | c57095_g1 | -9.015 | 3.22E-06 |
| 334 | c36475_g1 | -9.013 | 3.46E-06 |
| 335 | c49523_g1 | -9.012 | 3.46E-06 |
| 336 | c53918_g1 | -9.010 | 3.46E-06 |
| 337 | c47196_g1 | -9.003 | 3.46E-06 |
| 338 | c51021_g1 | -9.000 | 3.46E-06 |
| 339 | c20684_g1 | -8.993 | 3.72E-06 |
| 340 | c53617_g1 | -8.991 | 3.72E-06 |
| 341 | c55129_g1 | -8.990 | 3.72E-06 |

| | | | |
|---|---|---|---|
| 342 | c43940_g1 | -8.990 | 3.72E-06 |
| 343 | c10519_g1 | -8.986 | 3.72E-06 |
| 344 | c25201_g1 | -8.984 | 4.00E-06 |
| 345 | c53541_g1 | -8.983 | 4.00E-06 |
| 346 | c57315_g1 | -8.979 | 3.72E-06 |
| 347 | c53532_g1 | -8.979 | 3.72E-06 |
| 348 | c42772_g1 | -8.978 | 4.00E-06 |
| 349 | c57796_g2 | -8.973 | 4.00E-06 |
| 350 | c40914_g1 | -8.971 | 4.30E-06 |
| 351 | c42417_g1 | -8.970 | 4.30E-06 |
| 352 | c54008_g1 | -8.968 | 4.30E-06 |
| 353 | c47378_g1 | -8.965 | 4.30E-06 |
| 354 | c46715_g1 | -8.959 | 4.30E-06 |
| 355 | c53163_g1 | -8.958 | 4.30E-06 |
| 356 | c56913_g6 | -8.957 | 4.30E-06 |
| 357 | c109657_g1 | -8.956 | 4.63E-06 |
| 358 | c26619_g1 | -8.955 | 4.63E-06 |
| 359 | c55907_g1 | -8.940 | 4.98E-06 |
| 360 | c52088_g1 | -8.939 | 4.98E-06 |
| 361 | c56114_g2 | -8.939 | 4.98E-06 |
| 362 | c50508_g1 | -8.937 | 4.98E-06 |
| 363 | c42861_g1 | -8.935 | 4.98E-06 |
| 364 | c97179_g1 | -8.934 | 4.98E-06 |
| 365 | c56931_g1 | -8.933 | 4.98E-06 |
| 366 | c53026_g1 | -8.932 | 4.63E-06 |
| 367 | c49037_g1 | -8.932 | 4.63E-06 |
| 368 | c28991_g1 | -8.930 | 5.37E-06 |
| 369 | c55293_g1 | -8.930 | 4.98E-06 |
| 370 | c52836_g2 | -8.928 | 4.98E-06 |
| 371 | c36438_g1 | -8.924 | 5.37E-06 |
| 372 | c48395_g1 | -8.923 | 5.37E-06 |
| 373 | c72899_g1 | -8.919 | 5.37E-06 |
| 374 | c55809_g5 | -8.912 | 5.78E-06 |
| 375 | c47590_g1 | -8.911 | 5.78E-06 |
| 376 | c57122_g1 | -8.911 | 5.78E-06 |
| 377 | c50547_g1 | -8.910 | 5.78E-06 |
| 378 | c46900_g1 | -8.895 | 5.78E-06 |
| 379 | c52633_g2 | -8.887 | 6.73E-06 |
| 380 | c55142_g5 | -8.885 | 6.24E-06 |
| 381 | c54468_g2 | -8.883 | 6.73E-06 |
| 382 | c52127_g1 | -8.879 | 6.73E-06 |
| 383 | c36293_g1 | -8.874 | 6.24E-06 |
| 384 | c57032_g2 | -8.873 | 7.27E-06 |
| 385 | c10553_g1 | -8.872 | 6.73E-06 |
| 386 | c57501_g1 | -8.870 | 7.27E-06 |
| 387 | c55764_g1 | -8.869 | 7.27E-06 |
| 388 | c57386_g1 | -8.869 | 7.27E-06 |
| 389 | c52330_g1 | -8.864 | 7.85E-06 |
| 390 | c57484_g1 | -8.861 | 7.27E-06 |
| 391 | c57010_g1 | -8.857 | 7.85E-06 |
| 392 | c34677_g1 | -8.856 | 7.27E-06 |
| 393 | c49638_g1 | -8.852 | 7.85E-06 |
| 394 | c57126_g1 | -8.851 | 7.85E-06 |
| 395 | c56727_g1 | -8.850 | 7.85E-06 |
| 396 | c49394_g1 | -8.841 | 7.85E-06 |
| 397 | c54606_g1 | -8.840 | 7.85E-06 |
| 398 | c49389_g1 | -8.834 | 7.85E-06 |
| 399 | c12401_g1 | -8.833 | 7.85E-06 |
| 400 | c36775_g1 | -8.832 | 8.48E-06 |
| 401 | c55023_g1 | -8.819 | 9.18E-06 |
| 402 | c34513_g1 | -8.818 | 9.18E-06 |
| 403 | c24774_g1 | -8.815 | 9.93E-06 |
| 404 | c55121_g1 | -8.809 | 9.93E-06 |
| 405 | c44747_g1 | -8.808 | 8.48E-06 |
| 406 | c29575_g1 | -8.798 | 1.08E-05 |
| 407 | c58198_g3 | -8.792 | 1.08E-05 |
| 408 | c52057_g1 | -8.792 | 1.08E-05 |
| 409 | c25307_g1 | -8.788 | 1.08E-05 |
| 410 | c52778_g1 | -8.786 | 1.08E-05 |

| | | | |
|---|---|---|---|
| 411 | c51163_g1 | -8.786 | 1.08E-05 |
| 412 | c50698_g2 | -8.786 | 1.17E-05 |
| 413 | c57547_g1 | -8.778 | 1.17E-05 |
| 414 | c57306_g8 | -8.777 | 1.17E-05 |
| 415 | c26497_g1 | -8.772 | 1.17E-05 |
| 416 | c55702_g2 | -8.768 | 1.17E-05 |
| 417 | c56310_g1 | -8.767 | 1.26E-05 |
| 418 | c45312_g1 | -8.747 | 1.37E-05 |
| 419 | c55072_g1 | -8.746 | 1.37E-05 |
| 420 | c36246_g1 | -8.744 | 1.26E-05 |
| 421 | c57885_g2 | -8.744 | 1.37E-05 |
| 422 | c33259_g1 | -8.741 | 1.37E-05 |
| 423 | c71328_g1 | -8.740 | 1.37E-05 |
| 424 | c56692_g1 | -8.740 | 1.37E-05 |
| 425 | c55837_g1 | -8.738 | 1.37E-05 |
| 426 | c57362_g2 | -8.733 | 1.49E-05 |
| 427 | c56063_g4 | -8.732 | 1.49E-05 |
| 428 | c84143_g1 | -8.731 | 1.49E-05 |
| 429 | c49492_g1 | -8.731 | 1.49E-05 |
| 430 | c57034_g1 | -8.727 | 1.49E-05 |
| 431 | c54643_g1 | -8.727 | 1.49E-05 |
| 432 | c56291_g1 | -8.726 | 1.49E-05 |
| 433 | c42178_g1 | -8.722 | 1.49E-05 |
| 434 | c56371_g1 | -8.720 | 1.62E-05 |
| 435 | c54813_g2 | -8.719 | 1.49E-05 |
| 436 | c39708_g1 | -8.716 | 1.62E-05 |
| 437 | c35604_g1 | -8.715 | 1.62E-05 |
| 438 | c56498_g2 | -8.709 | 1.62E-05 |
| 439 | c52780_g2 | -8.708 | 1.62E-05 |
| 440 | c40857_g1 | -8.706 | 1.62E-05 |
| 441 | c57531_g2 | -8.703 | 1.62E-05 |
| 442 | c39856_g1 | -8.703 | 1.76E-05 |
| 443 | c45720_g1 | -8.702 | 1.76E-05 |
| 444 | c55962_g1 | -8.700 | 1.62E-05 |
| 445 | c10085_g1 | -8.698 | 1.62E-05 |
| 446 | c27264_g1 | -8.696 | 1.76E-05 |
| 447 | c45387_g1 | -8.694 | 1.62E-05 |
| 448 | c33638_g1 | -8.694 | 1.76E-05 |
| 449 | c57969_g2 | -8.690 | 1.76E-05 |
| 450 | c54032_g1 | -8.689 | 1.76E-05 |
| 451 | c49141_g2 | -8.683 | 1.76E-05 |
| 452 | c56659_g1 | -8.682 | 1.76E-05 |
| 453 | c54013_g4 | -8.680 | 1.91E-05 |
| 454 | c45595_g1 | -8.680 | 1.91E-05 |
| 455 | c55407_g3 | -8.675 | 1.91E-05 |
| 456 | c54485_g1 | -8.671 | 2.08E-05 |
| 457 | c46714_g1 | -8.665 | 2.08E-05 |
| 458 | c47404_g1 | -8.664 | 2.08E-05 |
| 459 | c56291_g2 | -8.664 | 2.08E-05 |
| 460 | c109573_g1 | -8.663 | 2.08E-05 |
| 461 | c42201_g1 | -8.660 | 2.08E-05 |
| 462 | c31444_g1 | -8.657 | 2.08E-05 |
| 463 | c57615_g2 | -8.656 | 2.08E-05 |
| 464 | c50815_g1 | -8.654 | 2.08E-05 |
| 465 | c55150_g1 | -8.645 | 2.27E-05 |
| 466 | c39254_g1 | -8.639 | 2.27E-05 |
| 467 | c47446_g1 | -8.633 | 2.47E-05 |
| 468 | c34065_g1 | -8.631 | 2.47E-05 |
| 469 | c52804_g1 | -8.629 | 2.47E-05 |
| 470 | c54525_g1 | -8.627 | 2.47E-05 |
| 471 | c40720_g1 | -8.627 | 2.47E-05 |
| 472 | c57589_g2 | -8.624 | 2.47E-05 |
| 473 | c54247_g1 | -8.617 | 2.70E-05 |
| 474 | c53551_g1 | -8.616 | 2.47E-05 |
| 475 | c84264_g1 | -8.616 | 2.08E-05 |
| 476 | c57879_g4 | -8.613 | 2.70E-05 |
| 477 | c55294_g1 | -8.610 | 2.70E-05 |
| 478 | c56077_g1 | -8.604 | 2.70E-05 |
| 479 | c57939_g1 | -8.600 | 2.70E-05 |

| | | | |
|---|---|---|---|
| 480 | c50604_g1 | -8.600 | 2.95E-05 |
| 481 | c53292_g1 | -8.594 | 2.95E-05 |
| 482 | c55509_g3 | -8.594 | 2.95E-05 |
| 483 | c57935_g3 | -8.589 | 2.95E-05 |
| 484 | c56342_g1 | -8.588 | 2.95E-05 |
| 485 | c57938_g6 | -8.585 | 2.95E-05 |
| 486 | c55796_g1 | -8.584 | 2.95E-05 |
| 487 | c55035_g1 | -8.574 | 3.22E-05 |
| 488 | c56510_g1 | -8.573 | 3.22E-05 |
| 489 | c42321_g1 | -8.571 | 3.22E-05 |
| 490 | c56623_g1 | -8.557 | 2.95E-05 |
| 491 | c48854_g1 | -8.553 | 3.52E-05 |
| 492 | c55941_g1 | -8.551 | 3.85E-05 |
| 493 | c46894_g1 | -8.547 | 3.85E-05 |
| 494 | c49117_g1 | -8.544 | 3.85E-05 |
| 495 | c56051_g2 | -8.542 | 3.85E-05 |
| 496 | c53508_g1 | -8.542 | 3.52E-05 |
| 497 | c49092_g1 | -8.541 | 3.85E-05 |
| 498 | c25207_g1 | -8.538 | 3.85E-05 |
| 499 | c39355_g1 | -8.534 | 3.85E-05 |
| 500 | c50357_g1 | -8.533 | 3.85E-05 |
| 501 | c55498_g2 | -8.533 | 3.85E-05 |
| 502 | c42360_g1 | -8.531 | 3.85E-05 |
| 503 | c54176_g1 | -8.522 | 4.22E-05 |
| 504 | c30784_g1 | -8.521 | 4.22E-05 |
| 505 | c56449_g1 | -8.521 | 4.22E-05 |
| 506 | c46442_g1 | -8.520 | 4.22E-05 |
| 507 | c56722_g3 | -8.518 | 4.22E-05 |
| 508 | c58278_g1 | -8.512 | 4.22E-05 |
| 509 | c56896_g1 | -8.511 | 4.63E-05 |
| 510 | c23843_g1 | -8.507 | 4.22E-05 |
| 511 | c53257_g2 | -8.506 | 4.63E-05 |
| 512 | c48488_g1 | -8.505 | 4.63E-05 |
| 513 | c41752_g1 | -8.504 | 4.63E-05 |
| 514 | c57572_g2 | -8.504 | 4.63E-05 |
| 515 | c55514_g1 | -8.504 | 4.63E-05 |
| 516 | c26811_g1 | -8.495 | 5.08E-05 |
| 517 | c53396_g1 | -8.495 | 5.08E-05 |
| 518 | c40771_g2 | -8.490 | 4.63E-05 |
| 519 | c55130_g2 | -8.488 | 5.08E-05 |
| 520 | c40141_g1 | -8.488 | 5.08E-05 |
| 521 | c10550_g1 | -8.487 | 5.08E-05 |
| 522 | c23548_g1 | -8.482 | 5.08E-05 |
| 523 | c53671_g2 | -8.481 | 5.08E-05 |
| 524 | c42782_g1 | -8.480 | 5.08E-05 |
| 525 | c55958_g5 | -8.472 | 5.57E-05 |
| 526 | c52188_g1 | -8.472 | 5.57E-05 |
| 527 | c56370_g1 | -8.470 | 5.08E-05 |
| 528 | c49951_g1 | -8.465 | 5.57E-05 |
| 529 | c34100_g1 | -8.463 | 5.57E-05 |
| 530 | c26857_g1 | -8.461 | 5.57E-05 |
| 531 | c46269_g1 | -8.458 | 6.12E-05 |
| 532 | c56598_g1 | -8.456 | 6.12E-05 |
| 533 | c15611_g1 | -8.451 | 6.12E-05 |
| 534 | c50850_g1 | -8.450 | 6.12E-05 |
| 535 | c41767_g1 | -8.449 | 6.12E-05 |
| 536 | c51241_g1 | -8.448 | 6.12E-05 |
| 537 | c51662_g1 | -8.447 | 6.12E-05 |
| 538 | c50095_g1 | -8.446 | 6.12E-05 |
| 539 | c57645_g1 | -8.446 | 6.12E-05 |
| 540 | c44325_g1 | -8.445 | 6.12E-05 |
| 541 | c51899_g1 | -8.443 | 6.12E-05 |
| 542 | c46981_g1 | -8.441 | 6.74E-05 |
| 543 | c36220_g1 | -8.440 | 6.12E-05 |
| 544 | c57414_g1 | -8.439 | 6.12E-05 |
| 545 | c53938_g1 | -8.437 | 6.74E-05 |
| 546 | c52220_g1 | -8.436 | 6.12E-05 |
| 547 | c57094_g2 | -8.435 | 6.74E-05 |
| 548 | c56831_g2 | -8.435 | 6.74E-05 |

| | | | |
|---|---|---|---|
| 549 | c57759_g4 | -8.434 | 6.74E-05 |
| 550 | c57185_g6 | -8.430 | 6.74E-05 |
| 551 | c57488_g9 | -8.426 | 6.12E-05 |
| 552 | c55033_g1 | -8.423 | 6.74E-05 |
| 553 | c57758_g1 | -8.422 | 6.74E-05 |
| 554 | c51814_g1 | -8.419 | 7.42E-05 |
| 555 | c47730_g1 | -8.418 | 6.74E-05 |
| 556 | c56207_g1 | -8.417 | 6.74E-05 |
| 557 | c57045_g1 | -8.414 | 6.74E-05 |
| 558 | c9502_g1 | -8.412 | 7.42E-05 |
| 559 | c44082_g1 | -8.409 | 7.42E-05 |
| 560 | c52656_g2 | -8.403 | 3.35E-15 |
| 561 | c38257_g1 | -8.399 | 7.42E-05 |
| 562 | c55715_g1 | -8.399 | 7.42E-05 |
| 563 | c49432_g1 | -8.397 | 7.42E-05 |
| 564 | c58241_g1 | -8.397 | 7.42E-05 |
| 565 | c50445_g1 | -8.388 | 8.17E-05 |
| 566 | c110061_g1 | -8.388 | 8.17E-05 |
| 567 | c50994_g1 | -8.388 | 8.17E-05 |
| 568 | c42413_g1 | -8.388 | 8.17E-05 |
| 569 | c48581_g1 | -8.387 | 7.42E-05 |
| 570 | c55680_g3 | -8.386 | 8.17E-05 |
| 571 | c26764_g1 | -8.384 | 8.17E-05 |
| 572 | c51056_g1 | -8.380 | 8.17E-05 |
| 573 | c13970_g1 | -8.380 | 8.17E-05 |
| 574 | c42618_g1 | -8.376 | 8.17E-05 |
| 575 | c52509_g1 | -8.371 | 9.02E-05 |
| 576 | c56042_g1 | -8.364 | 9.02E-05 |
| 577 | c45690_g1 | -8.362 | 9.02E-05 |
| 578 | c53569_g1 | -8.362 | 9.02E-05 |
| 579 | c42623_g1 | -8.360 | 9.02E-05 |
| 580 | c52058_g2 | -8.358 | 9.02E-05 |
| 581 | c57459_g1 | -8.353 | 9.95E-05 |
| 582 | c52378_g1 | -8.349 | 9.95E-05 |
| 583 | c53291_g1 | -8.342 | 9.95E-05 |
| 584 | c56991_g1 | -8.338 | 9.95E-05 |
| 585 | c43295_g1 | -8.333 | 9.95E-05 |
| 586 | c45656_g1 | -8.329 | 9.95E-05 |
| 587 | c25138_g1 | -8.329 | 9.95E-05 |
| 588 | c51253_g1 | -8.327 | 1.10E-04 |
| 589 | c27180_g1 | -8.325 | 1.10E-04 |
| 590 | c19595_g1 | -8.325 | 1.10E-04 |
| 591 | c54015_g1 | -8.324 | 1.10E-04 |
| 592 | c54038_g1 | -8.322 | 1.10E-04 |
| 593 | c37542_g1 | -8.322 | 1.10E-04 |
| 594 | c57561_g1 | -8.320 | 1.10E-04 |
| 595 | c30098_g1 | -8.319 | 9.95E-05 |
| 596 | c54042_g1 | -8.318 | 1.10E-04 |
| 597 | c52004_g1 | -8.313 | 1.10E-04 |
| 598 | c56942_g1 | -8.311 | 1.22E-04 |
| 599 | c56341_g1 | -8.306 | 1.22E-04 |
| 600 | c44849_g1 | -8.304 | 1.22E-04 |
| 601 | c33624_g1 | -8.301 | 1.22E-04 |
| 602 | c57737_g1 | -8.297 | 1.22E-04 |
| 603 | c52651_g2 | -8.297 | 1.22E-04 |
| 604 | c26639_g1 | -8.296 | 1.10E-04 |
| 605 | c53109_g1 | -8.295 | 1.22E-04 |
| 606 | c53970_g1 | -8.289 | 1.35E-04 |
| 607 | c42588_g1 | -8.287 | 1.22E-04 |
| 608 | c16657_g1 | -8.281 | 1.35E-04 |
| 609 | c123090_g1 | -8.280 | 1.35E-04 |
| 610 | c42248_g1 | -8.279 | 1.35E-04 |
| 611 | c52215_g2 | -8.275 | 1.49E-04 |
| 612 | c40773_g1 | -8.274 | 1.35E-04 |
| 613 | c21944_g1 | -8.269 | 1.49E-04 |
| 614 | c53555_g1 | -8.267 | 1.49E-04 |
| 615 | c26480_g1 | -8.267 | 1.49E-04 |
| 616 | c58121_g1 | -8.265 | 1.49E-04 |
| 617 | c55655_g6 | -8.261 | 1.49E-04 |

| | | | |
|---|---|---|---|
| 618 | c15369_g1 | -8.261 | 1.49E-04 |
| 619 | c56605_g1 | -8.259 | 1.49E-04 |
| 620 | c46275_g1 | -8.259 | 1.35E-04 |
| 621 | c52773_g1 | -8.258 | 1.49E-04 |
| 622 | c42815_g1 | -8.253 | 1.35E-04 |
| 623 | c48363_g1 | -8.246 | 1.66E-04 |
| 624 | c32215_g1 | -8.245 | 1.49E-04 |
| 625 | c52367_g1 | -8.244 | 1.66E-04 |
| 626 | c34663_g1 | -8.239 | 1.49E-04 |
| 627 | c54027_g1 | -8.235 | 1.66E-04 |
| 628 | c57638_g1 | -8.231 | 1.66E-04 |
| 629 | c47163_g1 | -8.225 | 1.84E-04 |
| 630 | c55465_g1 | -8.224 | 1.84E-04 |
| 631 | c46214_g1 | -8.220 | 1.84E-04 |
| 632 | c50344_g1 | -8.217 | 1.66E-04 |
| 633 | c55632_g1 | -8.214 | 1.84E-04 |
| 634 | c48431_g1 | -8.213 | 1.84E-04 |
| 635 | c48899_g1 | -8.212 | 1.66E-04 |
| 636 | c54235_g1 | -8.207 | 1.84E-04 |
| 637 | c41888_g1 | -8.204 | 2.05E-04 |
| 638 | c46095_g1 | -8.204 | 1.84E-04 |
| 639 | c39797_g1 | -8.203 | 1.84E-04 |
| 640 | c53249_g1 | -8.201 | 2.05E-04 |
| 641 | c40382_g1 | -8.200 | 2.05E-04 |
| 642 | c52555_g2 | -8.199 | 2.05E-04 |
| 643 | c46885_g1 | -8.199 | 2.05E-04 |
| 644 | c51359_g1 | -8.197 | 1.84E-04 |
| 645 | c46449_g1 | -8.192 | 2.05E-04 |
| 646 | c13996_g1 | -8.189 | 2.05E-04 |
| 647 | c56569_g1 | -8.188 | 2.05E-04 |
| 648 | c37720_g1 | -8.185 | 2.05E-04 |
| 649 | c31603_g1 | -8.185 | 2.05E-04 |
| 650 | c29763_g1 | -8.183 | 2.05E-04 |
| 651 | c37461_g1 | -8.179 | 2.05E-04 |
| 652 | c51800_g1 | -8.179 | 2.05E-04 |
| 653 | c57378_g1 | -8.175 | 2.28E-04 |
| 654 | c56930_g1 | -8.174 | 2.28E-04 |
| 655 | c52839_g1 | -8.172 | 2.28E-04 |
| 656 | c52878_g1 | -8.169 | 2.28E-04 |
| 657 | c47836_g1 | -8.167 | 2.28E-04 |
| 658 | c49256_g1 | -8.165 | 2.28E-04 |
| 659 | c53726_g1 | -8.159 | 2.54E-04 |
| 660 | c122935_g1 | -8.156 | 2.28E-04 |
| 661 | c27926_g1 | -8.155 | 2.54E-04 |
| 662 | c36366_g1 | -8.154 | 2.54E-04 |
| 663 | c45396_g1 | -8.151 | 2.28E-04 |
| 664 | c42729_g1 | -8.150 | 2.28E-04 |
| 665 | c23016_g1 | -8.148 | 2.54E-04 |
| 666 | c50715_g2 | -8.146 | 2.54E-04 |
| 667 | c47389_g1 | -8.143 | 2.54E-04 |
| 668 | c11914_g1 | -8.141 | 2.54E-04 |
| 669 | c57553_g1 | -8.141 | 2.54E-04 |
| 670 | c41557_g1 | -8.139 | 2.54E-04 |
| 671 | c54661_g2 | -8.137 | 2.54E-04 |
| 672 | c34286_g1 | -8.132 | 2.54E-04 |
| 673 | c52306_g2 | -8.131 | 2.54E-04 |
| 674 | c55797_g1 | -8.128 | 2.83E-04 |
| 675 | c54812_g1 | -8.128 | 2.54E-04 |
| 676 | c53545_g3 | -8.125 | 2.83E-04 |
| 677 | c40254_g1 | -8.123 | 2.83E-04 |
| 678 | c47246_g1 | -8.123 | 2.83E-04 |
| 679 | c23080_g1 | -8.123 | 2.83E-04 |
| 680 | c52030_g1 | -8.123 | 2.83E-04 |
| 681 | c55731_g1 | -8.121 | 2.83E-04 |
| 682 | c57837_g1 | -8.121 | 2.83E-04 |
| 683 | c52122_g1 | -8.119 | 2.83E-04 |
| 684 | c39196_g1 | -8.117 | 2.83E-04 |
| 685 | c54239_g1 | -8.116 | 2.83E-04 |
| 686 | c55330_g2 | -8.112 | 2.83E-04 |

| | | | |
|---|---|---|---|
| 687 | c27140_g1 | -8.112 | 2.83E-04 |
| 688 | c42248_g2 | -8.112 | 3.17E-04 |
| 689 | c51964_g1 | -8.110 | 2.83E-04 |
| 690 | c50878_g1 | -8.104 | 3.17E-04 |
| 691 | c53784_g3 | -8.099 | 3.17E-04 |
| 692 | c56512_g1 | -8.096 | 3.17E-04 |
| 693 | c57345_g1 | -8.090 | 3.17E-04 |
| 694 | c50483_g1 | -8.090 | 3.17E-04 |
| 695 | c56760_g1 | -8.089 | 3.17E-04 |
| 696 | c54120_g1 | -8.087 | 3.17E-04 |
| 697 | c57039_g2 | -8.085 | 3.17E-04 |
| 698 | c44282_g1 | -8.081 | 3.54E-04 |
| 699 | c48720_g1 | -8.081 | 3.54E-04 |
| 700 | c49223_g1 | -8.080 | 3.17E-04 |
| 701 | c50539_g1 | -8.068 | 3.54E-04 |
| 702 | c53003_g1 | -8.068 | 3.54E-04 |
| 703 | c26695_g1 | -8.065 | 3.54E-04 |
| 704 | c56232_g1 | -8.063 | 3.54E-04 |
| 705 | c50163_g1 | -8.062 | 3.54E-04 |
| 706 | c18743_g1 | -8.059 | 3.54E-04 |
| 707 | c10879_g1 | -8.055 | 3.54E-04 |
| 708 | c26365_g1 | -8.054 | 3.54E-04 |
| 709 | c52602_g1 | -8.051 | 3.54E-04 |
| 710 | c42923_g1 | -8.047 | 3.54E-04 |
| 711 | c58083_g1 | -8.046 | 3.97E-04 |
| 712 | c24147_g1 | -8.042 | 3.97E-04 |
| 713 | c53153_g1 | -8.036 | 4.45E-04 |
| 714 | c49965_g4 | -8.035 | 3.97E-04 |
| 715 | c24274_g1 | -8.035 | 3.97E-04 |
| 716 | c43416_g1 | -8.031 | 3.97E-04 |
| 717 | c57042_g1 | -8.029 | 4.45E-04 |
| 718 | c55927_g3 | -8.025 | 4.45E-04 |
| 719 | c45061_g1 | -8.025 | 3.97E-04 |
| 720 | c24411_g1 | -8.023 | 4.45E-04 |
| 721 | c56382_g1 | -8.017 | 4.45E-04 |
| 722 | c56772_g1 | -8.015 | 4.45E-04 |
| 723 | c56399_g1 | -8.011 | 4.45E-04 |
| 724 | c50056_g1 | -8.007 | 4.45E-04 |
| 725 | c52406_g1 | -8.004 | 4.45E-04 |
| 726 | c45516_g1 | -7.999 | 4.99E-04 |
| 727 | c57624_g6 | -7.998 | 4.45E-04 |
| 728 | c23672_g1 | -7.989 | 4.99E-04 |
| 729 | c49256_g2 | -7.984 | 4.99E-04 |
| 730 | c48885_g1 | -7.975 | 5.61E-04 |
| 731 | c57800_g2 | -7.975 | 5.61E-04 |
| 732 | c57759_g2 | -7.971 | 4.99E-04 |
| 733 | c56216_g3 | -7.971 | 5.61E-04 |
| 734 | c50599_g1 | -7.969 | 5.61E-04 |
| 735 | c58213_g1 | -7.968 | 5.61E-04 |
| 736 | c57296_g1 | -7.966 | 5.61E-04 |
| 737 | c56287_g2 | -7.965 | 5.61E-04 |
| 738 | c51786_g1 | -7.964 | 5.61E-04 |
| 739 | c40687_g1 | -7.964 | 5.61E-04 |
| 740 | c46491_g1 | -7.964 | 5.61E-04 |
| 741 | c49956_g1 | -7.956 | 5.61E-04 |
| 742 | c53600_g1 | -7.952 | 6.32E-04 |
| 743 | c43981_g3 | -7.951 | 5.61E-04 |
| 744 | c42555_g1 | -7.948 | 6.32E-04 |
| 745 | c46325_g1 | -7.947 | 6.32E-04 |
| 746 | c56238_g1 | -7.945 | 6.32E-04 |
| 747 | c39309_g1 | -7.944 | 6.32E-04 |
| 748 | c57533_g1 | -7.943 | 6.32E-04 |
| 749 | c52997_g1 | -7.942 | 5.61E-04 |
| 750 | c54747_g1 | -7.942 | 6.32E-04 |
| 751 | c50948_g1 | -7.935 | 6.32E-04 |
| 752 | c54581_g1 | -7.933 | 6.32E-04 |
| 753 | c41233_g1 | -7.925 | 7.12E-04 |
| 754 | c56142_g2 | -7.924 | 6.32E-04 |
| 755 | c49916_g1 | -7.921 | 6.32E-04 |

| | | | |
|---|---|---|---|
| 756 | c45675_g1 | -7.919 | 6.32E-04 |
| 757 | c43849_g1 | -7.915 | 6.32E-04 |
| 758 | c43115_g1 | -7.913 | 7.12E-04 |
| 759 | c84559_g1 | -7.911 | 5.61E-04 |
| 760 | c54975_g2 | -7.911 | 7.12E-04 |
| 761 | c58069_g1 | -7.904 | 7.12E-04 |
| 762 | c31515_g1 | -7.904 | 7.12E-04 |
| 763 | c57406_g1 | -7.903 | 7.12E-04 |
| 764 | c57216_g1 | -7.901 | 7.12E-04 |
| 765 | c52765_g1 | -7.900 | 7.12E-04 |
| 766 | c40022_g1 | -7.889 | 7.12E-04 |
| 767 | c56686_g3 | -7.889 | 8.03E-04 |
| 768 | c44266_g1 | -7.887 | 7.12E-04 |
| 769 | c53878_g2 | -7.885 | 8.03E-04 |
| 770 | c43838_g1 | -7.880 | 8.03E-04 |
| 771 | c43290_g1 | -7.880 | 8.03E-04 |
| 772 | c55416_g1 | -7.870 | 8.03E-04 |
| 773 | c53879_g1 | -7.865 | 9.08E-04 |
| 774 | c55060_g1 | -7.856 | 8.03E-04 |
| 775 | c52935_g1 | -7.853 | 9.08E-04 |
| 776 | c53608_g1 | -7.850 | 8.03E-04 |
| 777 | c52608_g1 | -7.849 | 9.08E-04 |
| 778 | c56826_g1 | -7.849 | 9.08E-04 |
| 779 | c49577_g2 | -7.849 | 9.08E-04 |
| 780 | c54510_g3 | -7.847 | 9.08E-04 |
| 781 | c45160_g1 | -7.843 | 9.08E-04 |
| 782 | c54496_g1 | -7.842 | 9.08E-04 |
| 783 | c51736_g1 | -7.842 | 9.08E-04 |
| 784 | c42232_g1 | -7.837 | 1.03E-03 |
| 785 | c57623_g6 | -7.837 | 1.03E-03 |
| 786 | c55132_g1 | -7.835 | 9.08E-04 |
| 787 | c53280_g1 | -7.835 | 9.08E-04 |
| 788 | c47582_g1 | -7.834 | 9.08E-04 |
| 789 | c55872_g1 | -7.833 | 1.03E-03 |
| 790 | c40824_g1 | -7.829 | 9.08E-04 |
| 791 | c47525_g1 | -7.826 | 1.03E-03 |
| 792 | c56609_g1 | -7.826 | 1.03E-03 |
| 793 | c59379_g1 | -7.825 | 8.03E-04 |
| 794 | c46706_g1 | -7.824 | 9.08E-04 |
| 795 | c55486_g1 | -7.824 | 9.08E-04 |
| 796 | c54323_g2 | -7.824 | 1.03E-03 |
| 797 | c45247_g2 | -7.822 | 1.03E-03 |
| 798 | c52573_g1 | -7.818 | 1.03E-03 |
| 799 | c28092_g1 | -7.818 | 1.03E-03 |
| 800 | c57284_g2 | -7.817 | 1.03E-03 |
| 801 | c43872_g1 | -7.814 | 1.03E-03 |
| 802 | c45092_g1 | -7.813 | 1.03E-03 |
| 803 | c47595_g1 | -7.812 | 1.17E-03 |
| 804 | c48097_g1 | -7.809 | 1.03E-03 |
| 805 | c54179_g1 | -7.809 | 1.03E-03 |
| 806 | c50715_g3 | -7.807 | 1.17E-03 |
| 807 | c53533_g1 | -7.807 | 1.17E-03 |
| 808 | c44053_g1 | -7.800 | 1.17E-03 |
| 809 | c56375_g1 | -7.797 | 2.61E-04 |
| 810 | c48299_g1 | -7.796 | 1.17E-03 |
| 811 | c110033_g1 | -7.791 | 2.98E-04 |
| 812 | c47193_g1 | -7.791 | 2.98E-04 |
| 813 | c53765_g8 | -7.785 | 2.98E-04 |
| 814 | c58179_g1 | -7.779 | 2.98E-04 |
| 815 | c53794_g1 | -7.779 | 2.98E-04 |
| 816 | c46238_g1 | -7.776 | 3.41E-04 |
| 817 | c55009_g1 | -7.775 | 2.98E-04 |
| 818 | c47666_g1 | -7.773 | 2.61E-04 |
| 819 | c56977_g3 | -7.769 | 3.41E-04 |
| 820 | c42518_g1 | -7.769 | 3.41E-04 |
| 821 | c54167_g2 | -7.767 | 2.98E-04 |
| 822 | c53843_g1 | -7.766 | 3.41E-04 |
| 823 | c51967_g1 | -7.765 | 2.98E-04 |
| 824 | c43025_g1 | -7.758 | 3.41E-04 |

| | | | |
|---|---|---|---|
| 825 | c53676_g1 | -7.758 | 3.41E-04 |
| 826 | c56148_g1 | -7.758 | 2.98E-04 |
| 827 | c39949_g1 | -7.757 | 3.41E-04 |
| 828 | c28798_g1 | -7.755 | 3.41E-04 |
| 829 | c31520_g1 | -7.755 | 3.41E-04 |
| 830 | c56023_g1 | -7.750 | 3.41E-04 |
| 831 | c19561_g1 | -7.749 | 3.90E-04 |
| 832 | c56392_g1 | -7.744 | 3.90E-04 |
| 833 | c39298_g1 | -7.743 | 3.90E-04 |
| 834 | c33331_g1 | -7.737 | 3.90E-04 |
| 835 | c57764_g2 | -7.735 | 3.90E-04 |
| 836 | c57653_g1 | -7.732 | 3.90E-04 |
| 837 | c44092_g1 | -7.731 | 3.90E-04 |
| 838 | c52434_g1 | -7.728 | 3.90E-04 |
| 839 | c50586_g1 | -7.722 | 3.90E-04 |
| 840 | c48887_g1 | -7.721 | 3.90E-04 |
| 841 | c58082_g1 | -7.721 | 3.90E-04 |
| 842 | c56194_g1 | -7.719 | 3.90E-04 |
| 843 | c56848_g3 | -7.718 | 3.90E-04 |
| 844 | c48943_g1 | -7.718 | 3.90E-04 |
| 845 | c44533_g1 | -7.718 | 3.90E-04 |
| 846 | c96869_g1 | -7.717 | 3.90E-04 |
| 847 | c58054_g10 | -7.710 | 3.90E-04 |
| 848 | c52660_g1 | -7.709 | 4.48E-04 |
| 849 | c50657_g1 | -7.707 | 3.90E-04 |
| 850 | c39617_g1 | -7.707 | 3.90E-04 |
| 851 | c39362_g1 | -7.703 | 4.48E-04 |
| 852 | c55286_g1 | -7.703 | 4.48E-04 |
| 853 | c48515_g2 | -7.701 | 3.90E-04 |
| 854 | c50941_g1 | -7.694 | 4.48E-04 |
| 855 | c56747_g1 | -7.694 | 4.48E-04 |
| 856 | c57218_g2 | -7.687 | 4.48E-04 |
| 857 | c52831_g1 | -7.684 | 4.48E-04 |
| 858 | c33392_g1 | -7.679 | 4.48E-04 |
| 859 | c49610_g1 | -7.670 | 5.15E-04 |
| 860 | c55759_g1 | -7.669 | 4.48E-04 |
| 861 | c53595_g1 | -7.669 | 5.15E-04 |
| 862 | c54516_g1 | -7.665 | 5.15E-04 |
| 863 | c56558_g1 | -7.662 | 5.15E-04 |
| 864 | c110791_g1 | -7.661 | 5.15E-04 |
| 865 | c38225_g1 | -7.659 | 5.15E-04 |
| 866 | c44573_g1 | -7.659 | 5.15E-04 |
| 867 | c56807_g1 | -7.649 | 3.90E-04 |
| 868 | c55743_g1 | -7.646 | 5.93E-04 |
| 869 | c34353_g1 | -7.644 | 4.48E-04 |
| 870 | c34152_g1 | -7.644 | 4.48E-04 |
| 871 | c55195_g1 | -7.641 | 5.93E-04 |
| 872 | c53723_g2 | -7.636 | 5.15E-04 |
| 873 | c57270_g1 | -7.634 | 5.93E-04 |
| 874 | c38009_g2 | -7.634 | 5.93E-04 |
| 875 | c49158_g1 | -7.626 | 5.93E-04 |
| 876 | c57887_g1 | -7.626 | 5.93E-04 |
| 877 | c50911_g1 | -7.625 | 5.93E-04 |
| 878 | c46163_g1 | -7.623 | 5.93E-04 |
| 879 | c52996_g1 | -7.623 | 5.93E-04 |
| 880 | c54036_g2 | -7.623 | 5.93E-04 |
| 881 | c53054_g1 | -7.618 | 5.93E-04 |
| 882 | c45279_g1 | -7.615 | 5.93E-04 |
| 883 | c18631_g1 | -7.613 | 5.93E-04 |
| 884 | c56186_g1 | -7.613 | 5.93E-04 |
| 885 | c51015_g1 | -7.612 | 6.84E-04 |
| 886 | c122528_g1 | -7.612 | 6.84E-04 |
| 887 | c43163_g1 | -7.608 | 6.84E-04 |
| 888 | c50988_g1 | -7.608 | 5.93E-04 |
| 889 | c48444_g1 | -7.607 | 6.84E-04 |
| 890 | c33914_g1 | -7.605 | 6.84E-04 |
| 891 | c32837_g1 | -7.604 | 6.84E-04 |
| 892 | c40432_g1 | -7.601 | 5.93E-04 |
| 893 | c50412_g1 | -7.593 | 5.93E-04 |

| | | | |
|---|---|---|---|
| 894 | c8866_g1 | -7.592 | 6.84E-04 |
| 895 | c57215_g1 | -7.589 | 6.84E-04 |
| 896 | c52618_g1 | -7.586 | 6.84E-04 |
| 897 | c35611_g1 | -7.580 | 6.84E-04 |
| 898 | c51600_g1 | -7.575 | 7.91E-04 |
| 899 | c52818_g5 | -7.572 | 6.84E-04 |
| 900 | c54971_g8 | -7.570 | 6.84E-04 |
| 901 | c45761_g1 | -7.568 | 6.84E-04 |
| 902 | c55896_g1 | -7.567 | 6.84E-04 |
| 903 | c56155_g1 | -7.566 | 7.91E-04 |
| 904 | c47543_g1 | -7.563 | 7.91E-04 |
| 905 | c43300_g1 | -7.560 | 7.91E-04 |
| 906 | c46227_g1 | -7.555 | 5.93E-04 |
| 907 | c25629_g2 | -7.554 | 7.91E-04 |
| 908 | c55446_g1 | -7.553 | 7.91E-04 |
| 909 | c39368_g1 | -7.553 | 7.91E-04 |
| 910 | c46456_g1 | -7.552 | 7.91E-04 |
| 911 | c26901_g1 | -7.552 | 6.84E-04 |
| 912 | c26006_g1 | -7.551 | 9.17E-04 |
| 913 | c31091_g1 | -7.548 | 7.91E-04 |
| 914 | c46859_g1 | -7.544 | 7.91E-04 |
| 915 | c24254_g1 | -7.531 | 9.17E-04 |
| 916 | c58114_g1 | -7.531 | 9.17E-04 |
| 917 | c57491_g1 | -7.523 | 9.17E-04 |
| 918 | c53931_g1 | -7.523 | 9.17E-04 |
| 919 | c56592_g1 | -7.516 | 9.17E-04 |
| 920 | c55930_g1 | -7.515 | 9.17E-04 |
| 921 | c44867_g1 | -7.514 | 9.17E-04 |
| 922 | c49331_g1 | -7.512 | 1.06E-03 |
| 923 | c49374_g1 | -7.512 | 1.06E-03 |
| 924 | c25218_g2 | -7.512 | 1.06E-03 |
| 925 | c54724_g1 | -7.503 | 1.06E-03 |
| 926 | c58154_g4 | -7.502 | 1.06E-03 |
| 927 | c48884_g1 | -7.497 | 9.17E-04 |
| 928 | c54264_g1 | -7.495 | 1.06E-03 |
| 929 | c39222_g1 | -7.495 | 1.06E-03 |
| 930 | c56337_g1 | -7.486 | 1.06E-03 |
| 931 | c56200_g1 | -7.486 | 1.06E-03 |
| 932 | c57956_g1 | -7.484 | 1.06E-03 |
| 933 | c98263_g1 | -7.483 | 1.06E-03 |
| 934 | c109750_g1 | -7.482 | 1.06E-03 |
| 935 | c52230_g1 | -7.481 | 1.06E-03 |
| 936 | c56661_g2 | -7.480 | 1.24E-03 |
| 937 | c57632_g1 | -7.474 | 1.06E-03 |
| 938 | c53453_g1 | -7.474 | 1.06E-03 |
| 939 | c56213_g1 | -7.471 | 1.24E-03 |
| 940 | c47758_g1 | -7.466 | 1.06E-03 |
| 941 | c56209_g1 | -7.465 | 1.06E-03 |
| 942 | c55002_g1 | -7.463 | 1.24E-03 |
| 943 | c57895_g1 | -7.453 | 1.24E-03 |
| 944 | c34922_g1 | -7.453 | 1.24E-03 |
| 945 | c52605_g1 | -7.452 | 1.24E-03 |
| 946 | c47514_g1 | -7.452 | 1.24E-03 |
| 947 | c57505_g2 | -7.450 | 1.24E-03 |
| 948 | c49501_g1 | -7.445 | 1.06E-03 |
| 949 | c52168_g1 | -7.443 | 1.24E-03 |
| 950 | c51666_g19 | -7.439 | 1.44E-03 |
| 951 | c50711_g1 | -7.433 | 1.24E-03 |
| 952 | c54515_g1 | -7.431 | 1.24E-03 |
| 953 | c57011_g1 | -7.431 | 1.44E-03 |
| 954 | c58225_g1 | -7.430 | 1.44E-03 |
| 955 | c46117_g1 | -7.425 | 9.17E-04 |
| 956 | c51711_g1 | -7.420 | 1.44E-03 |
| 957 | c52212_g1 | -7.419 | 1.44E-03 |
| 958 | c34530_g1 | -7.419 | 1.44E-03 |
| 959 | c55990_g3 | -7.411 | 1.24E-03 |
| 960 | c48798_g1 | -7.410 | 1.44E-03 |
| 961 | c53421_g1 | -7.410 | 1.44E-03 |
| 962 | c34582_g1 | -7.409 | 1.44E-03 |

| | | | |
|---|---|---|---|
| 963 | c57204_g1 | -7.400 | 1.24E-03 |
| 964 | c44262_g1 | -7.400 | 1.44E-03 |
| 965 | c40209_g1 | -7.399 | 1.44E-03 |
| 966 | c49112_g1 | -7.399 | 1.44E-03 |
| 967 | c10610_g1 | -7.399 | 1.44E-03 |
| 968 | c37475_g2 | -7.388 | 1.44E-03 |
| 969 | c55840_g1 | -7.387 | 1.69E-03 |
| 970 | c84063_g1 | -7.377 | 1.44E-03 |
| 971 | c57159_g1 | -7.377 | 1.69E-03 |
| 972 | c57851_g1 | -7.377 | 1.69E-03 |
| 973 | c55605_g1 | -7.365 | 1.69E-03 |
| 974 | c15646_g1 | -7.365 | 1.69E-03 |
| 975 | c58520_g1 | -7.365 | 1.69E-03 |
| 976 | c32808_g1 | -7.354 | 1.98E-03 |
| 977 | c56024_g1 | -7.354 | 1.69E-03 |
| 978 | c57473_g1 | -7.354 | 1.69E-03 |
| 979 | c50211_g1 | -7.354 | 1.69E-03 |
| 980 | c56300_g1 | -7.354 | 1.69E-03 |
| 981 | c56742_g1 | -7.343 | 1.98E-03 |
| 982 | c41200_g1 | -7.343 | 1.98E-03 |
| 983 | c44608_g1 | -7.343 | 1.69E-03 |
| 984 | c46039_g1 | -7.343 | 1.69E-03 |
| 985 | c50148_g1 | -7.343 | 1.69E-03 |
| 986 | c46404_g1 | -7.342 | 1.69E-03 |
| 987 | c57087_g1 | -7.332 | 1.98E-03 |
| 988 | c50276_g1 | -7.331 | 1.98E-03 |
| 989 | c52207_g1 | -7.331 | 1.98E-03 |
| 990 | c56913_g4 | -7.321 | 1.98E-03 |
| 991 | c54212_g1 | -7.308 | 1.98E-03 |
| 992 | c54846_g3 | -7.308 | 1.98E-03 |
| 993 | c38895_g1 | -7.296 | 2.32E-03 |
| 994 | c57322_g2 | -7.295 | 1.98E-03 |
| 995 | c50967_g1 | -7.294 | 1.98E-03 |
| 996 | c54346_g1 | -7.286 | 2.32E-03 |
| 997 | c51703_g1 | -7.286 | 2.32E-03 |
| 998 | c49929_g1 | -7.285 | 2.32E-03 |
| 999 | c57398_g1 | -7.284 | 2.32E-03 |
| 1000 | c50923_g1 | -7.283 | 2.32E-03 |
| 1001 | c15585_g1 | -7.282 | 1.98E-03 |
| 1002 | c53623_g2 | -7.274 | 2.32E-03 |
| 1003 | c57672_g2 | -7.273 | 2.32E-03 |
| 1004 | c19096_g1 | -7.271 | 2.32E-03 |
| 1005 | c36300_g1 | -7.271 | 2.32E-03 |
| 1006 | c51978_g1 | -7.263 | 2.74E-03 |
| 1007 | c50645_g1 | -7.258 | 2.32E-03 |
| 1008 | c44491_g1 | -7.258 | 2.32E-03 |
| 1009 | c52756_g1 | -7.257 | 2.32E-03 |
| 1010 | c49571_g2 | -7.250 | 2.74E-03 |
| 1011 | c24327_g2 | -7.248 | 2.74E-03 |
| 1012 | c109557_g1 | -7.246 | 2.32E-03 |
| 1013 | c56226_g1 | -7.246 | 2.32E-03 |
| 1014 | c54445_g1 | -7.245 | 2.32E-03 |
| 1015 | c33096_g1 | -7.245 | 2.32E-03 |
| 1016 | c54218_g2 | -7.244 | 2.32E-03 |
| 1017 | c53521_g1 | -7.237 | 2.74E-03 |
| 1018 | c53713_g1 | -7.235 | 2.74E-03 |
| 1019 | c50942_g1 | -7.233 | 2.74E-03 |
| 1020 | c56199_g2 | -7.232 | 2.32E-03 |
| 1021 | c54268_g1 | -7.227 | 3.23E-03 |
| 1022 | c57185_g4 | -7.211 | 3.23E-03 |
| 1023 | c56031_g1 | -7.210 | 2.74E-03 |
| 1024 | c30327_g1 | -7.210 | 2.74E-03 |
| 1025 | c46695_g1 | -7.210 | 2.74E-03 |
| 1026 | c43466_g1 | -7.201 | 3.23E-03 |
| 1027 | c48592_g1 | -7.201 | 1.98E-03 |
| 1028 | c45991_g2 | -7.196 | 3.23E-03 |
| 1029 | c45857_g1 | -7.196 | 3.23E-03 |
| 1030 | c57858_g2 | -7.188 | 3.23E-03 |
| 1031 | c57878_g2 | -7.188 | 3.23E-03 |

| | | | |
|---|---|---|---|
| 1032 | c43981_g1 | -7.187 | 3.23E-03 |
| 1033 | c57285_g1 | -7.187 | 3.23E-03 |
| 1034 | c57100_g2 | -7.186 | 1.13E-11 |
| 1035 | c57948_g1 | -7.185 | 3.23E-03 |
| 1036 | c50726_g1 | -7.184 | 3.23E-03 |
| 1037 | c53018_g1 | -7.184 | 3.23E-03 |
| 1038 | c50739_g1 | -7.178 | 2.74E-03 |
| 1039 | c58206_g1 | -7.177 | 3.83E-03 |
| 1040 | c58174_g2 | -7.177 | 3.83E-03 |
| 1041 | c38314_g1 | -7.175 | 3.23E-03 |
| 1042 | c56540_g1 | -7.174 | 3.23E-03 |
| 1043 | c35853_g1 | -7.171 | 3.23E-03 |
| 1044 | c50126_g1 | -7.170 | 3.23E-03 |
| 1045 | c56845_g1 | -7.163 | 3.83E-03 |
| 1046 | c57306_g5 | -7.161 | 3.83E-03 |
| 1047 | c48175_g1 | -7.160 | 3.83E-03 |
| 1048 | c55133_g1 | -7.156 | 3.23E-03 |
| 1049 | c58018_g3 | -7.155 | 3.23E-03 |
| 1050 | c56479_g1 | -7.155 | 3.23E-03 |
| 1051 | c46469_g1 | -7.146 | 1.47E-11 |
| 1052 | c44180_g1 | -7.132 | 3.83E-03 |
| 1053 | c43895_g1 | -7.131 | 3.83E-03 |
| 1054 | c49057_g1 | -7.124 | 4.54E-03 |
| 1055 | c57271_g1 | -7.124 | 4.54E-03 |
| 1056 | c40801_g1 | -7.121 | 3.83E-03 |
| 1057 | c52586_g1 | -7.119 | 3.83E-03 |
| 1058 | c54893_g1 | -7.118 | 3.83E-03 |
| 1059 | c39599_g2 | -7.116 | 3.83E-03 |
| 1060 | c56181_g1 | -7.112 | 4.54E-03 |
| 1061 | c15308_g1 | -7.111 | 3.83E-03 |
| 1062 | c47150_g1 | -7.108 | 4.54E-03 |
| 1063 | c48009_g2 | -7.108 | 4.54E-03 |
| 1064 | c46160_g1 | -7.104 | 4.54E-03 |
| 1065 | c47626_g1 | -7.104 | 4.54E-03 |
| 1066 | c14157_g1 | -7.097 | 4.54E-03 |
| 1067 | c50027_g1 | -7.094 | 4.54E-03 |
| 1068 | c32783_g1 | -7.085 | 3.83E-03 |
| 1069 | c39644_g1 | -7.083 | 4.54E-03 |
| 1070 | c50399_g1 | -7.076 | 5.41E-03 |
| 1071 | c55270_g1 | -7.075 | 4.54E-03 |
| 1072 | c53404_g3 | -7.075 | 4.54E-03 |
| 1073 | c45750_g1 | -7.067 | 3.83E-03 |
| 1074 | c57321_g1 | -7.065 | 5.41E-03 |
| 1075 | c48904_g1 | -7.061 | 5.41E-03 |
| 1076 | c54365_g1 | -7.057 | 5.41E-03 |
| 1077 | c56233_g1 | -7.049 | 5.41E-03 |
| 1078 | c56883_g2 | -7.044 | 5.41E-03 |
| 1079 | c52792_g1 | -7.044 | 4.54E-03 |
| 1080 | c48425_g1 | -7.040 | 6.45E-03 |
| 1081 | c38003_g1 | -7.040 | 5.41E-03 |
| 1082 | c58057_g4 | -7.038 | 5.41E-03 |
| 1083 | c51169_g4 | -7.038 | 5.41E-03 |
| 1084 | c50185_g1 | -7.030 | 3.83E-03 |
| 1085 | c54664_g1 | -7.027 | 5.41E-03 |
| 1086 | c55919_g7 | -7.025 | 5.41E-03 |
| 1087 | c53614_g1 | -7.025 | 5.41E-03 |
| 1088 | c50214_g1 | -7.022 | 5.41E-03 |
| 1089 | c55170_g4 | -7.020 | 5.41E-03 |
| 1090 | c53939_g1 | -7.004 | 5.41E-03 |
| 1091 | c53092_g3 | -7.001 | 6.45E-03 |
| 1092 | c43078_g1 | -6.993 | 6.45E-03 |
| 1093 | c49643_g1 | -6.987 | 6.45E-03 |
| 1094 | c51249_g1 | -6.985 | 6.45E-03 |
| 1095 | c49517_g1 | -6.985 | 6.45E-03 |
| 1096 | c57735_g1 | -6.985 | 6.45E-03 |
| 1097 | c57008_g1 | -6.982 | 6.45E-03 |
| 1098 | c53092_g1 | -6.982 | 6.45E-03 |
| 1099 | c109529_g1 | -6.979 | 6.45E-03 |
| 1100 | c45370_g1 | -6.976 | 5.41E-03 |

| | | | |
|---|---|---|---|
| 1101 | c55490_g1 | -6.968 | 6.45E-03 |
| 1102 | c55911_g4 | -6.963 | 6.45E-03 |
| 1103 | c57500_g1 | -6.958 | 5.41E-03 |
| 1104 | c53999_g1 | -6.957 | 6.45E-03 |
| 1105 | c124540_g1 | -6.956 | 7.73E-03 |
| 1106 | c31075_g1 | -6.952 | 7.73E-03 |
| 1107 | c54105_g2 | -6.949 | 7.73E-03 |
| 1108 | c57426_g1 | -6.947 | 6.45E-03 |
| 1109 | c49215_g1 | -6.939 | 6.45E-03 |
| 1110 | c54769_g1 | -6.938 | 7.73E-03 |
| 1111 | c57214_g1 | -6.938 | 7.73E-03 |
| 1112 | c43033_g1 | -6.936 | 6.45E-03 |
| 1113 | c30979_g1 | -6.936 | 6.45E-03 |
| 1114 | c54152_g1 | -6.935 | 7.73E-03 |
| 1115 | c29535_g1 | -6.934 | 9.28E-03 |
| 1116 | c31394_g2 | -6.932 | 7.73E-03 |
| 1117 | c57331_g1 | -6.921 | 7.73E-03 |
| 1118 | c36298_g1 | -6.918 | 7.73E-03 |
| 1119 | c57078_g1 | -6.918 | 7.73E-03 |
| 1120 | c57240_g1 | -6.914 | 7.73E-03 |
| 1121 | c51532_g1 | -6.914 | 7.73E-03 |
| 1122 | c56474_g1 | -6.911 | 7.73E-03 |
| 1123 | c57727_g1 | -6.911 | 9.28E-03 |
| 1124 | c53115_g1 | -6.907 | 9.28E-03 |
| 1125 | c53172_g1 | -6.904 | 9.28E-03 |
| 1126 | c41580_g1 | -6.900 | 7.73E-03 |
| 1127 | c44937_g1 | -6.900 | 7.73E-03 |
| 1128 | c45174_g1 | -6.893 | 7.73E-03 |
| 1129 | c45302_g1 | -6.886 | 9.28E-03 |
| 1130 | c55982_g1 | -6.883 | 9.28E-03 |
| 1131 | c56089_g1 | -6.876 | 9.28E-03 |
| 1132 | c39345_g1 | -6.874 | 7.73E-03 |
| 1133 | c44954_g1 | -6.872 | 9.28E-03 |
| 1134 | c52373_g1 | -6.869 | 1.12E-02 |
| 1135 | c49409_g2 | -6.864 | 9.28E-03 |
| 1136 | c41121_g1 | -6.862 | 7.73E-03 |
| 1137 | c51475_g1 | -6.862 | 9.28E-03 |
| 1138 | c53052_g4 | -6.858 | 9.28E-03 |
| 1139 | c43089_g1 | -6.856 | 1.12E-02 |
| 1140 | c53721_g1 | -6.854 | 9.28E-03 |
| 1141 | c30118_g1 | -6.852 | 1.12E-02 |
| 1142 | c57046_g7 | -6.851 | 9.28E-03 |
| 1143 | c57771_g1 | -6.849 | 9.28E-03 |
| 1144 | c35816_g1 | -6.845 | 9.28E-03 |
| 1145 | c56750_g1 | -6.843 | 1.12E-02 |
| 1146 | c46899_g1 | -6.843 | 1.12E-02 |
| 1147 | c49449_g1 | -6.837 | 1.12E-02 |
| 1148 | c50364_g1 | -6.836 | 9.28E-03 |
| 1149 | c52817_g1 | -6.835 | 1.12E-02 |
| 1150 | c55589_g1 | -6.835 | 1.12E-02 |
| 1151 | c54262_g1 | -6.833 | 1.12E-02 |
| 1152 | c42008_g2 | -6.832 | 1.12E-02 |
| 1153 | c46478_g1 | -6.830 | 9.28E-03 |
| 1154 | c55746_g1 | -6.825 | 1.12E-02 |
| 1155 | c109999_g1 | -6.820 | 1.12E-02 |
| 1156 | c57718_g1 | -6.816 | 9.28E-03 |
| 1157 | c34557_g1 | -6.815 | 1.12E-02 |
| 1158 | c51753_g1 | -6.810 | 1.12E-02 |
| 1159 | c45735_g1 | -6.806 | 1.12E-02 |
| 1160 | c53730_g1 | -6.806 | 1.12E-02 |
| 1161 | c50121_g1 | -6.805 | 1.12E-02 |
| 1162 | c56434_g1 | -6.801 | 1.12E-02 |
| 1163 | c47490_g1 | -6.800 | 1.35E-02 |
| 1164 | c56140_g1 | -6.796 | 9.28E-03 |
| 1165 | c58172_g1 | -6.795 | 1.35E-02 |
| 1166 | c52389_g1 | -6.791 | 1.12E-02 |
| 1167 | c57110_g1 | -6.771 | 1.35E-02 |
| 1168 | c55582_g2 | -6.771 | 1.35E-02 |
| 1169 | c51699_g1 | -6.770 | 1.12E-02 |

| | | | |
|---|---|---|---|
| 1170 | c57237_g4 | -6.761 | 1.35E-02 |
| 1171 | c57750_g1 | -6.761 | 1.35E-02 |
| 1172 | c109742_g1 | -6.757 | 1.35E-02 |
| 1173 | c55186_g1 | -6.756 | 1.84E-10 |
| 1174 | c48979_g1 | -6.756 | 1.35E-02 |
| 1175 | c55170_g11 | -6.756 | 1.35E-02 |
| 1176 | c52276_g1 | -6.755 | 1.12E-02 |
| 1177 | c54011_g1 | -6.749 | 1.12E-02 |
| 1178 | c36281_g1 | -6.745 | 1.35E-02 |
| 1179 | c14192_g1 | -6.742 | 1.64E-02 |
| 1180 | c50732_g1 | -6.739 | 1.35E-02 |
| 1181 | c49246_g1 | -6.737 | 1.64E-02 |
| 1182 | c53357_g1 | -6.730 | 1.35E-02 |
| 1183 | c58074_g1 | -6.726 | 1.64E-02 |
| 1184 | c29104_g1 | -6.722 | 1.64E-02 |
| 1185 | c21277_g1 | -6.719 | 1.64E-02 |
| 1186 | c54115_g1 | -6.716 | 1.64E-02 |
| 1187 | c56675_g1 | -6.716 | 1.64E-02 |
| 1188 | c52467_g1 | -6.711 | 1.64E-02 |
| 1189 | c57961_g1 | -6.705 | 1.64E-02 |
| 1190 | c58134_g1 | -6.705 | 1.64E-02 |
| 1191 | c39688_g2 | -6.705 | 1.64E-02 |
| 1192 | c49414_g2 | -6.701 | 1.64E-02 |
| 1193 | c71523_g1 | -6.701 | 1.64E-02 |
| 1194 | c42176_g1 | -6.698 | 2.00E-02 |
| 1195 | c50490_g1 | -6.695 | 1.64E-02 |
| 1196 | c122431_g1 | -6.692 | 1.64E-02 |
| 1197 | c54831_g1 | -6.689 | 1.64E-02 |
| 1198 | c56715_g1 | -6.689 | 1.64E-02 |
| 1199 | c96846_g1 | -6.678 | 2.00E-02 |
| 1200 | c30933_g1 | -6.674 | 1.64E-02 |
| 1201 | c56785_g1 | -6.674 | 1.64E-02 |
| 1202 | c53050_g1 | -6.668 | 1.64E-02 |
| 1203 | c45667_g1 | -6.665 | 2.00E-02 |
| 1204 | c27311_g1 | -6.659 | 1.35E-02 |
| 1205 | c57908_g1 | -6.659 | 2.00E-02 |
| 1206 | c40499_g1 | -6.659 | 2.00E-02 |
| 1207 | c55056_g1 | -6.657 | 1.64E-02 |
| 1208 | c55765_g1 | -6.657 | 1.64E-02 |
| 1209 | c33840_g1 | -6.654 | 1.64E-02 |
| 1210 | c48127_g1 | -6.652 | 2.00E-02 |
| 1211 | c56372_g1 | -6.652 | 2.00E-02 |
| 1212 | c58160_g1 | -6.650 | 2.00E-02 |
| 1213 | c36439_g1 | -6.649 | 1.35E-02 |
| 1214 | c55542_g1 | -6.647 | 1.64E-02 |
| 1215 | c57445_g1 | -6.643 | 2.00E-02 |
| 1216 | c123047_g1 | -6.641 | 2.00E-02 |
| 1217 | c24374_g1 | -6.638 | 2.00E-02 |
| 1218 | c49375_g1 | -6.638 | 2.00E-02 |
| 1219 | c53439_g2 | -6.635 | 2.00E-02 |
| 1220 | c42834_g2 | -6.633 | 1.64E-02 |
| 1221 | c56783_g1 | -6.631 | 1.64E-02 |
| 1222 | c44620_g1 | -6.628 | 4.17E-10 |
| 1223 | c23833_g1 | -6.628 | 2.00E-02 |
| 1224 | c51967_g2 | -6.624 | 1.64E-02 |
| 1225 | c27671_g1 | -6.621 | 2.00E-02 |
| 1226 | c109598_g1 | -6.615 | 2.00E-02 |
| 1227 | c45259_g1 | -6.614 | 2.00E-02 |
| 1228 | c49084_g1 | -6.606 | 2.00E-02 |
| 1229 | c55628_g2 | -6.605 | 4.80E-10 |
| 1230 | c51814_g2 | -6.599 | 2.00E-02 |
| 1231 | c54953_g1 | -6.599 | 2.00E-02 |
| 1232 | c55149_g2 | -6.599 | 2.00E-02 |
| 1233 | c11450_g1 | -6.557 | 2.00E-02 |
| 1234 | c96899_g1 | -6.540 | 1.64E-02 |
| 1235 | c57944_g1 | -6.522 | 8.21E-10 |
| 1236 | c56469_g1 | -6.463 | 1.19E-09 |
| 1237 | c23602_g1 | -6.427 | 1.52E-09 |
| 1238 | c57311_g1 | -6.133 | 9.42E-09 |

| | | | |
|---|---|---|---|
| 1239 | c53192_g1 | -6.056 | 1.51E-08 |
| 1240 | c55389_g3 | -6.008 | 2.01E-08 |
| 1241 | c43382_g1 | -5.988 | 3.60E-12 |
| 1242 | c122426_g1 | -5.954 | 8.10E-10 |
| 1243 | c15658_g1 | -5.917 | 3.56E-08 |
| 1244 | c55101_g2 | -5.887 | 4.30E-08 |
| 1245 | c45950_g1 | -5.854 | 5.20E-08 |
| 1246 | c55094_g2 | -5.847 | 5.37E-08 |
| 1247 | c53824_g1 | -5.835 | 5.92E-08 |
| 1248 | c57671_g1 | -5.831 | 5.92E-08 |
| 1249 | c49173_g1 | -5.747 | 9.76E-08 |
| 1250 | c49608_g1 | -5.589 | 2.57E-07 |
| 1251 | c48875_g2 | -5.587 | 2.57E-07 |
| 1252 | c56350_g1 | -5.553 | 3.11E-07 |
| 1253 | c48317_g1 | -5.538 | 1.11E-08 |
| 1254 | c47895_g1 | -5.528 | 2.85E-10 |
| 1255 | c49323_g1 | -5.501 | 1.40E-08 |
| 1256 | c38325_g1 | -5.439 | 6.10E-07 |
| 1257 | c52176_g1 | -5.361 | 9.27E-07 |
| 1258 | c54151_g2 | -5.317 | 4.62E-09 |
| 1259 | c56989_g1 | -5.259 | 1.72E-06 |
| 1260 | c57012_g2 | -5.223 | 2.06E-06 |
| 1261 | c52387_g1 | -5.210 | 2.16E-06 |
| 1262 | c57993_g1 | -5.182 | 2.61E-06 |
| 1263 | c49256_g3 | -5.085 | 4.45E-06 |
| 1264 | c56473_g1 | -5.085 | 4.45E-06 |
| 1265 | c30202_g1 | -5.052 | 2.41E-08 |
| 1266 | c52880_g2 | -5.028 | 6.03E-06 |
| 1267 | c71326_g1 | -5.013 | 2.71E-07 |
| 1268 | c51521_g1 | -5.001 | 7.05E-06 |
| 1269 | c52261_g1 | -5.000 | 7.05E-06 |
| 1270 | c43343_g2 | -4.989 | 3.14E-07 |
| 1271 | c57410_g2 | -4.982 | 7.83E-06 |
| 1272 | c50627_g1 | -4.977 | 5.73E-09 |
| 1273 | c54222_g1 | -4.956 | 9.18E-06 |
| 1274 | c56935_g1 | -4.929 | 1.08E-05 |
| 1275 | c53372_g1 | -4.889 | 1.35E-05 |
| 1276 | c56129_g1 | -4.864 | 1.51E-05 |
| 1277 | c35155_g1 | -4.804 | 9.27E-07 |
| 1278 | c52069_g1 | -4.789 | 2.25E-05 |
| 1279 | c51859_g2 | -4.785 | 2.25E-05 |
| 1280 | c45382_g1 | -4.767 | 1.37E-07 |
| 1281 | c53478_g1 | -4.735 | 3.03E-05 |
| 1282 | c58237_g1 | -4.709 | 1.95E-07 |
| 1283 | c51542_g1 | -4.675 | 5.78E-09 |
| 1284 | c48420_g1 | -4.639 | 4.96E-05 |
| 1285 | c52737_g1 | -4.636 | 4.66E-05 |
| 1286 | c10566_g1 | -4.586 | 1.61E-06 |
| 1287 | c53816_g3 | -4.584 | 6.41E-05 |
| 1288 | c56891_g1 | -4.582 | 4.16E-07 |
| 1289 | c53620_g1 | -4.550 | 7.80E-05 |
| 1290 | c57338_g4 | -4.511 | 8.91E-05 |
| 1291 | c43425_g1 | -4.505 | 6.55E-07 |
| 1292 | c49669_g1 | -4.489 | 2.76E-06 |
| 1293 | c57017_g1 | -4.476 | 1.09E-04 |
| 1294 | c51424_g1 | -4.441 | 1.25E-04 |
| 1295 | c54437_g1 | -4.435 | 3.65E-06 |
| 1296 | c57114_g1 | -4.417 | 8.89E-08 |
| 1297 | c51265_g2 | -4.406 | 1.54E-04 |
| 1298 | c54902_g1 | -4.369 | 1.91E-04 |
| 1299 | c41895_g1 | -4.332 | 6.54E-06 |
| 1300 | c54886_g1 | -4.326 | 3.21E-07 |
| 1301 | c8189_g1 | -4.318 | 1.96E-06 |
| 1302 | c50650_g1 | -4.315 | 7.45E-06 |
| 1303 | c38514_g1 | -4.287 | 2.56E-04 |
| 1304 | c55956_g1 | -4.263 | 2.67E-06 |
| 1305 | c55795_g3 | -4.261 | 2.97E-04 |
| 1306 | c57565_g3 | -4.229 | 3.23E-06 |
| 1307 | c46562_g1 | -4.214 | 3.56E-06 |

| | | | |
|---|---|---|---|
| 1308 | c47887_g1 | -4.162 | 5.10E-04 |
| 1309 | c50395_g1 | -4.131 | 5.52E-04 |
| 1310 | c47898_g1 | -4.122 | 1.71E-06 |
| 1311 | c10120_g1 | -4.104 | 6.48E-04 |
| 1312 | c57583_g2 | -3.995 | 6.45E-07 |
| 1313 | c51752_g1 | -3.983 | 3.80E-06 |
| 1314 | c56222_g1 | -3.942 | 4.79E-06 |
| 1315 | c55022_g3 | -3.940 | 1.38E-03 |
| 1316 | c50481_g1 | -3.934 | 2.63E-08 |
| 1317 | c55391_g1 | -3.934 | 2.08E-06 |
| 1318 | c56319_g1 | -3.889 | 2.08E-06 |
| 1319 | c47668_g1 | -3.889 | 2.15E-05 |
| 1320 | c42523_g1 | -3.874 | 1.21E-07 |
| 1321 | c44589_g1 | -3.869 | 3.05E-07 |
| 1322 | c48878_g2 | -3.858 | 1.79E-03 |
| 1323 | c84052_g1 | -3.822 | 1.17E-06 |
| 1324 | c57516_g2 | -3.818 | 1.30E-07 |
| 1325 | c44183_g2 | -3.816 | 3.18E-05 |
| 1326 | c25398_g1 | -3.815 | 9.76E-04 |
| 1327 | c57918_g1 | -3.803 | 1.07E-03 |
| 1328 | c23731_g1 | -3.778 | 1.20E-05 |
| 1329 | c57728_g2 | -3.778 | 1.17E-05 |
| 1330 | c46499_g1 | -3.763 | 4.22E-05 |
| 1331 | c10794_g1 | -3.750 | 1.18E-03 |
| 1332 | c10169_g1 | -3.692 | 6.16E-05 |
| 1333 | c57972_g2 | -3.685 | 1.93E-03 |
| 1334 | c50100_g1 | -3.680 | 2.05E-04 |
| 1335 | c50506_g1 | -3.679 | 2.05E-04 |
| 1336 | c58285_g1 | -3.679 | 8.92E-06 |
| 1337 | c11557_g1 | -3.640 | 7.98E-05 |
| 1338 | c57208_g1 | -3.622 | 2.61E-04 |
| 1339 | c58162_g2 | -3.602 | 2.95E-04 |
| 1340 | c56996_g1 | -3.599 | 2.61E-03 |
| 1341 | c50665_g2 | -3.596 | 3.14E-04 |
| 1342 | c50153_g1 | -3.579 | 3.35E-04 |
| 1343 | c47466_g1 | -3.556 | 2.89E-03 |
| 1344 | c49648_g2 | -3.536 | 5.78E-06 |
| 1345 | c54237_g1 | -3.514 | 3.57E-03 |
| 1346 | c49895_g1 | -3.503 | 1.90E-05 |
| 1347 | c51159_g1 | -3.498 | 1.65E-04 |
| 1348 | c55076_g1 | -3.489 | 3.97E-03 |
| 1349 | c52950_g2 | -3.486 | 5.25E-04 |
| 1350 | c53447_g1 | -3.483 | 2.65E-05 |
| 1351 | c45110_g1 | -3.461 | 4.42E-03 |
| 1352 | c97073_g1 | -3.451 | 2.09E-04 |
| 1353 | c50986_g2 | -3.449 | 6.40E-04 |
| 1354 | c32839_g1 | -3.430 | 6.04E-06 |
| 1355 | c45955_g1 | -3.427 | 6.85E-04 |
| 1356 | c37503_g1 | -3.421 | 8.20E-06 |
| 1357 | c46416_g1 | -3.403 | 7.06E-06 |
| 1358 | c55397_g1 | -3.401 | 7.84E-04 |
| 1359 | c43077_g1 | -3.395 | 2.02E-05 |
| 1360 | c32709_g1 | -3.373 | 1.06E-04 |
| 1361 | c48509_g1 | -3.363 | 3.69E-06 |
| 1362 | c52157_g1 | -3.360 | 1.10E-04 |
| 1363 | c46396_g1 | -3.347 | 1.79E-05 |
| 1364 | c49547_g1 | -3.340 | 1.03E-03 |
| 1365 | c58098_g1 | -3.327 | 7.69E-03 |
| 1366 | c25900_g1 | -3.318 | 7.69E-03 |
| 1367 | c56130_g1 | -3.314 | 9.41E-06 |
| 1368 | c51933_g1 | -3.309 | 5.36E-05 |
| 1369 | c58203_g1 | -3.290 | 4.64E-04 |
| 1370 | c52265_g1 | -3.277 | 4.89E-04 |
| 1371 | c51719_g1 | -3.273 | 1.37E-03 |
| 1372 | c57589_g1 | -3.251 | 2.99E-05 |
| 1373 | c54266_g1 | -3.230 | 2.13E-04 |
| 1374 | c57699_g1 | -3.224 | 2.22E-04 |
| 1375 | c50415_g1 | -3.201 | 5.83E-05 |
| 1376 | c41380_g1 | -3.200 | 1.15E-04 |

| | | | |
|---|---|---|---|
| 1377 | c26573_g1 | -3.195 | 1.84E-03 |
| 1378 | c45821_g1 | -3.176 | 1.37E-02 |
| 1379 | c10667_g1 | -3.165 | 5.00E-04 |
| 1380 | c49848_g2 | -3.165 | 5.00E-04 |
| 1381 | c34184_g1 | -3.163 | 7.23E-05 |
| 1382 | c55628_g1 | -3.152 | 4.69E-06 |
| 1383 | c46841_g1 | -3.130 | 1.93E-05 |
| 1384 | c52305_g1 | -3.109 | 6.63E-04 |
| 1385 | c42941_g2 | -3.100 | 2.28E-05 |
| 1386 | c56779_g2 | -3.100 | 2.11E-05 |
| 1387 | c45746_g1 | -3.097 | 6.63E-04 |
| 1388 | c58233_g1 | -3.086 | 3.45E-05 |
| 1389 | c57083_g1 | -3.084 | 2.89E-03 |
| 1390 | c57605_g1 | -3.080 | 3.12E-03 |
| 1391 | c56722_g2 | -3.058 | 4.81E-05 |
| 1392 | c44046_g1 | -3.052 | 3.03E-05 |
| 1393 | c57480_g1 | -3.049 | 2.67E-05 |
| 1394 | c52188_g2 | -3.019 | 5.91E-04 |
| 1395 | c52993_g1 | -3.010 | 1.06E-03 |
| 1396 | c48575_g2 | -3.000 | 1.69E-04 |
| 1397 | c54611_g1 | -2.996 | 2.69E-04 |
| 1398 | c55692_g1 | -2.995 | 5.73E-05 |
| 1399 | c58347_g1 | -2.985 | 2.87E-04 |
| 1400 | c49611_g1 | -2.972 | 1.20E-03 |
| 1401 | c57241_g1 | -2.970 | 6.36E-05 |
| 1402 | c45419_g1 | -2.962 | 2.08E-04 |
| 1403 | c35068_g1 | -2.941 | 8.65E-04 |
| 1404 | c49418_g2 | -2.929 | 5.15E-05 |
| 1405 | c48817_g1 | -2.927 | 3.78E-04 |
| 1406 | c55912_g1 | -2.914 | 1.00E-03 |
| 1407 | c58058_g1 | -2.907 | 5.41E-05 |
| 1408 | c49278_g1 | -2.902 | 1.41E-04 |
| 1409 | c57149_g1 | -2.887 | 6.01E-05 |
| 1410 | c54978_g1 | -2.886 | 1.16E-03 |
| 1411 | c54679_g1 | -2.862 | 7.74E-05 |
| 1412 | c26676_g1 | -2.856 | 1.17E-04 |
| 1413 | c47346_g1 | -2.847 | 3.94E-04 |
| 1414 | c84087_g1 | -2.840 | 6.21E-05 |
| 1415 | c30304_g1 | -2.838 | 7.17E-04 |
| 1416 | c45428_g1 | -2.833 | 2.76E-04 |
| 1417 | c42822_g1 | -2.831 | 2.03E-04 |
| 1418 | c58017_g1 | -2.828 | 1.64E-04 |
| 1419 | c54457_g1 | -2.823 | 8.61E-05 |
| 1420 | c54671_g9 | -2.821 | 4.62E-05 |
| 1421 | c54698_g1 | -2.799 | 3.19E-04 |
| 1422 | c57460_g2 | -2.793 | 2.51E-04 |
| 1423 | c30138_g1 | -2.792 | 8.67E-05 |
| 1424 | c57382_g1 | -2.791 | 2.71E-03 |
| 1425 | c47100_g1 | -2.774 | 2.68E-05 |
| 1426 | c36605_g1 | -2.769 | 5.93E-04 |
| 1427 | c54240_g1 | -2.768 | 2.79E-04 |
| 1428 | c54395_g1 | -2.755 | 6.40E-04 |
| 1429 | c55536_g1 | -2.754 | 2.03E-03 |
| 1430 | c47897_g1 | -2.724 | 2.38E-03 |
| 1431 | c84251_g1 | -2.720 | 2.38E-03 |
| 1432 | c55170_g12 | -2.710 | 2.51E-03 |
| 1433 | c36021_g1 | -2.705 | 3.85E-04 |
| 1434 | c57361_g1 | -2.695 | 1.16E-04 |
| 1435 | c44548_g1 | -2.688 | 8.71E-04 |
| 1436 | c56292_g1 | -2.687 | 4.32E-03 |
| 1437 | c54538_g1 | -2.676 | 1.06E-04 |
| 1438 | c35673_g1 | -2.664 | 4.32E-03 |
| 1439 | c34305_g1 | -2.663 | 4.63E-03 |
| 1440 | c44191_g1 | -2.642 | 3.48E-03 |
| 1441 | c57858_g3 | -2.637 | 7.38E-04 |
| 1442 | c57829_g1 | -2.615 | 1.79E-02 |
| 1443 | c55277_g1 | -2.605 | 6.37E-04 |
| 1444 | c51968_g1 | -2.604 | 1.96E-02 |
| 1445 | c57575_g2 | -2.578 | 2.14E-04 |

| | | | |
|---|---|---|---|
| 1446 | c51177_g1 | -2.574 | 7.03E-03 |
| 1447 | c52177_g1 | -2.569 | 9.82E-04 |
| 1448 | c54398_g1 | -2.564 | 7.79E-04 |
| 1449 | c50088_g1 | -2.556 | 7.03E-03 |
| 1450 | c57014_g2 | -2.549 | 5.64E-04 |
| 1451 | c46072_g1 | -2.543 | 3.96E-04 |
| 1452 | c57285_g3 | -2.519 | 8.72E-03 |
| 1453 | c29083_g1 | -2.496 | 2.44E-04 |
| 1454 | c42741_g1 | -2.486 | 1.45E-03 |
| 1455 | c53426_g6 | -2.463 | 7.26E-03 |
| 1456 | c50648_g1 | -2.461 | 3.92E-03 |
| 1457 | c46405_g1 | -2.460 | 7.26E-03 |
| 1458 | c57624_g4 | -2.454 | 2.32E-03 |
| 1459 | c55667_g1 | -2.453 | 8.96E-04 |
| 1460 | c55565_g1 | -2.453 | 7.73E-04 |
| 1461 | c52349_g1 | -2.452 | 2.54E-03 |
| 1462 | c53117_g1 | -2.446 | 1.77E-03 |
| 1463 | c84044_g1 | -2.443 | 4.26E-04 |
| 1464 | c53225_g3 | -2.437 | 7.70E-03 |
| 1465 | c41358_g1 | -2.427 | 3.22E-04 |
| 1466 | c71344_g1 | -2.425 | 4.55E-03 |
| 1467 | c52108_g1 | -2.424 | 4.79E-03 |
| 1468 | c50692_g1 | -2.420 | 1.03E-03 |
| 1469 | c54041_g1 | -2.408 | 5.04E-03 |
| 1470 | c56235_g1 | -2.403 | 3.16E-03 |
| 1471 | c54316_g1 | -2.398 | 4.57E-04 |
| 1472 | c57856_g1 | -2.398 | 5.43E-04 |
| 1473 | c10659_g1 | -2.396 | 5.94E-04 |
| 1474 | c44257_g1 | -2.394 | 5.18E-04 |
| 1475 | c53066_g1 | -2.392 | 1.01E-03 |
| 1476 | c46085_g1 | -2.377 | 5.07E-04 |
| 1477 | c56571_g1 | -2.375 | 7.05E-03 |
| 1478 | c43981_g2 | -2.353 | 1.74E-03 |
| 1479 | c50640_g1 | -2.347 | 2.13E-03 |
| 1480 | c47222_g2 | -2.345 | 2.78E-03 |
| 1481 | c43198_g1 | -2.345 | 9.51E-04 |
| 1482 | c52161_g1 | -2.342 | 3.97E-03 |
| 1483 | c54488_g1 | -2.342 | 9.20E-04 |
| 1484 | c42617_g1 | -2.338 | 7.99E-03 |
| 1485 | c96840_g1 | -2.327 | 9.09E-04 |
| 1486 | c55000_g2 | -2.326 | 5.06E-04 |
| 1487 | c34714_g1 | -2.324 | 6.35E-04 |
| 1488 | c53583_g2 | -2.316 | 4.29E-03 |
| 1489 | c55864_g2 | -2.313 | 1.83E-02 |
| 1490 | c57050_g2 | -2.308 | 9.02E-04 |
| 1491 | c54423_g1 | -2.305 | 9.06E-03 |
| 1492 | c42021_g1 | -2.287 | 2.76E-03 |
| 1493 | c54578_g1 | -2.279 | 2.40E-03 |
| 1494 | c41087_g1 | -2.278 | 1.98E-02 |
| 1495 | c48374_g1 | -2.276 | 1.17E-03 |
| 1496 | c56893_g2 | -2.266 | 1.03E-02 |
| 1497 | c49206_g1 | -2.247 | 2.41E-03 |
| 1498 | c57573_g2 | -2.246 | 6.24E-03 |
| 1499 | c37106_g1 | -2.242 | 2.09E-03 |
| 1500 | c50637_g1 | -2.242 | 6.24E-03 |
| 1501 | c55965_g5 | -2.237 | 1.17E-02 |
| 1502 | c50854_g1 | -2.233 | 8.74E-04 |
| 1503 | c25365_g1 | -2.212 | 8.57E-04 |
| 1504 | c55516_g1 | -2.199 | 1.45E-03 |
| 1505 | c71560_g1 | -2.196 | 2.25E-03 |
| 1506 | c55008_g1 | -2.186 | 2.50E-03 |
| 1507 | c45565_g1 | -2.172 | 8.28E-03 |
| 1508 | c53305_g1 | -2.167 | 1.81E-03 |
| 1509 | c32546_g1 | -2.166 | 4.10E-03 |
| 1510 | c52075_g1 | -2.163 | 2.02E-03 |
| 1511 | c49935_g1 | -2.162 | 4.10E-03 |
| 1512 | c43718_g1 | -2.158 | 1.66E-03 |
| 1513 | c39500_g1 | -2.150 | 5.05E-03 |
| 1514 | c56982_g1 | -2.137 | 7.14E-03 |

| | | | |
|---|---|---|---|
| 1515 | c71416_g1 | -2.134 | 1.23E-03 |
| 1516 | c38825_g1 | -2.129 | 1.74E-02 |
| 1517 | c56248_g1 | -2.124 | 1.05E-02 |
| 1518 | c122580_g1 | -2.122 | 3.01E-03 |
| 1519 | c34387_g1 | -2.119 | 2.76E-03 |
| 1520 | c56786_g5 | -2.114 | 1.09E-03 |
| 1521 | c35450_g1 | -2.113 | 2.18E-03 |
| 1522 | c53141_g1 | -2.113 | 4.32E-03 |
| 1523 | c50790_g1 | -2.086 | 8.49E-03 |
| 1524 | c54722_g1 | -2.085 | 1.16E-02 |
| 1525 | c109627_g1 | -2.084 | 4.30E-03 |
| 1526 | c34110_g1 | -2.084 | 1.98E-03 |
| 1527 | c57088_g1 | -2.080 | 3.02E-03 |
| 1528 | c52445_g1 | -2.079 | 1.82E-02 |
| 1529 | c23862_g1 | -2.078 | 1.27E-03 |
| 1530 | c41421_g1 | -2.077 | 1.82E-02 |
| 1531 | c53403_g1 | -2.068 | 7.42E-03 |
| 1532 | c54682_g5 | -2.062 | 9.45E-03 |
| 1533 | c55862_g1 | -2.060 | 5.38E-03 |
| 1534 | c37914_g1 | -2.054 | 1.01E-02 |
| 1535 | c55689_g1 | -2.037 | 1.62E-03 |
| 1536 | c53801_g1 | -2.036 | 2.09E-03 |
| 1537 | c56445_g1 | -2.029 | 1.11E-02 |
| 1538 | c49720_g1 | -2.024 | 5.58E-03 |
| 1539 | c56873_g1 | -2.017 | 6.26E-03 |
| 1540 | c46581_g1 | -2.014 | 1.77E-03 |
| 1541 | c56045_g1 | -2.013 | 2.50E-03 |
| 1542 | c56324_g1 | -2.006 | 1.65E-02 |
| 1543 | c57314_g1 | -1.982 | 2.92E-03 |
| 1544 | c57525_g2 | -1.981 | 5.64E-03 |
| 1545 | c51784_g1 | -1.968 | 2.36E-03 |
| 1546 | c51362_g1 | -1.956 | 5.11E-03 |
| 1547 | c50386_g1 | -1.942 | 5.40E-03 |
| 1548 | c56392_g2 | -1.939 | 1.52E-02 |
| 1549 | c23688_g1 | -1.935 | 6.17E-03 |
| 1550 | c52695_g1 | -1.934 | 1.28E-02 |
| 1551 | c57811_g1 | -1.933 | 8.11E-03 |
| 1552 | c52987_g1 | -1.926 | 1.65E-02 |
| 1553 | c57822_g1 | -1.912 | 5.73E-03 |
| 1554 | c51589_g2 | -1.906 | 4.81E-03 |
| 1555 | c49765_g1 | -1.894 | 3.24E-03 |
| 1556 | c57968_g1 | -1.886 | 5.27E-03 |
| 1557 | c57346_g1 | -1.876 | 2.01E-02 |
| 1558 | c25467_g2 | -1.872 | 1.54E-02 |
| 1559 | c55540_g3 | -1.862 | 4.90E-03 |
| 1560 | c52782_g1 | -1.847 | 4.09E-03 |
| 1561 | c51358_g4 | -1.844 | 6.32E-03 |
| 1562 | c33443_g1 | -1.839 | 9.90E-03 |
| 1563 | c57306_g4 | -1.837 | 1.51E-02 |
| 1564 | c49452_g1 | -1.834 | 1.34E-02 |
| 1565 | c52884_g1 | -1.833 | 8.35E-03 |
| 1566 | c57566_g4 | -1.822 | 4.80E-03 |
| 1567 | c37474_g1 | -1.810 | 6.65E-03 |
| 1568 | c42274_g1 | -1.808 | 2.00E-02 |
| 1569 | c32403_g1 | -1.795 | 1.23E-02 |
| 1570 | c25331_g1 | -1.787 | 1.29E-02 |
| 1571 | c57178_g1 | -1.780 | 9.11E-03 |
| 1572 | c51585_g1 | -1.772 | 7.28E-03 |
| 1573 | c51917_g1 | -1.771 | 2.00E-02 |
| 1574 | c46274_g1 | -1.769 | 7.09E-03 |
| 1575 | c50915_g1 | -1.766 | 1.31E-02 |
| 1576 | c53542_g1 | -1.756 | 6.30E-03 |
| 1577 | c47762_g1 | -1.754 | 8.52E-03 |
| 1578 | c57100_g1 | -1.753 | 1.15E-02 |
| 1579 | c49411_g1 | -1.753 | 8.25E-03 |
| 1580 | c51782_g1 | -1.740 | 5.79E-03 |
| 1581 | c56356_g1 | -1.734 | 1.83E-02 |
| 1582 | c55400_g2 | -1.714 | 1.21E-02 |
| 1583 | c15618_g1 | -1.701 | 1.98E-02 |

| | | | |
|---|---|---|---|
| 1584 | c47909_g1 | -1.692 | 2.05E-02 |
| 1585 | c56584_g3 | -1.690 | 8.84E-03 |
| 1586 | c51849_g1 | -1.687 | 9.23E-03 |
| 1587 | c57352_g1 | -1.671 | 8.44E-03 |
| 1588 | c51856_g1 | -1.667 | 1.62E-02 |
| 1589 | c35185_g1 | -1.648 | 8.47E-03 |
| 1590 | c53268_g1 | -1.641 | 9.52E-03 |
| 1591 | c50396_g3 | -1.627 | 1.45E-02 |
| 1592 | c33044_g1 | -1.620 | 1.37E-02 |
| 1593 | c96827_g1 | -1.615 | 1.26E-02 |
| 1594 | c47020_g1 | -1.612 | 1.22E-02 |
| 1595 | c55727_g4 | -1.600 | 9.97E-03 |
| 1596 | c56753_g1 | -1.586 | 1.59E-02 |
| 1597 | c51017_g1 | -1.576 | 1.89E-02 |
| 1598 | c56741_g1 | -1.550 | 1.38E-02 |
| 1599 | c50761_g3 | -1.548 | 1.62E-02 |
| 1600 | c46488_g2 | -1.529 | 1.66E-02 |
| 1601 | c54618_g1 | -1.518 | 1.61E-02 |
| 1602 | c30806_g1 | -1.514 | 2.02E-02 |
| 1603 | c44570_g1 | -1.511 | 1.82E-02 |
| 1604 | c54301_g1 | -1.502 | 1.78E-02 |
| 1605 | c71380_g1 | -1.498 | 1.86E-02 |
| 1606 | c50040_g1 | -1.482 | 1.84E-02 |
| 1607 | c57332_g3 | -1.458 | 2.06E-02 |
| 1608 | c55492_g4 | 1.291 | 1.96E-02 |
| 1609 | c47130_g1 | 1.292 | 1.97E-02 |
| 1610 | c41582_g1 | 1.320 | 1.68E-02 |
| 1611 | c54696_g1 | 1.325 | 1.95E-02 |
| 1612 | c39759_g1 | 1.327 | 1.55E-02 |
| 1613 | c49578_g1 | 1.328 | 1.46E-02 |
| 1614 | c58111_g1 | 1.332 | 1.75E-02 |
| 1615 | c53382_g1 | 1.339 | 1.79E-02 |
| 1616 | c52205_g2 | 1.350 | 2.06E-02 |
| 1617 | c57276_g5 | 1.354 | 1.79E-02 |
| 1618 | c56939_g1 | 1.364 | 1.39E-02 |
| 1619 | c16562_g1 | 1.372 | 1.68E-02 |
| 1620 | c51316_g1 | 1.384 | 1.28E-02 |
| 1621 | c57145_g1 | 1.386 | 1.72E-02 |
| 1622 | c54210_g1 | 1.387 | 1.69E-02 |
| 1623 | c56272_g1 | 1.390 | 1.68E-02 |
| 1624 | c56069_g1 | 1.390 | 1.22E-02 |
| 1625 | c48009_g1 | 1.392 | 1.19E-02 |
| 1626 | c39584_g1 | 1.395 | 1.32E-02 |
| 1627 | c56499_g1 | 1.395 | 2.03E-02 |
| 1628 | c55095_g1 | 1.402 | 1.06E-02 |
| 1629 | c49653_g1 | 1.403 | 1.58E-02 |
| 1630 | c23145_g1 | 1.409 | 1.02E-02 |
| 1631 | c57125_g1 | 1.411 | 1.06E-02 |
| 1632 | c33554_g2 | 1.411 | 1.33E-02 |
| 1633 | c26950_g1 | 1.423 | 1.52E-02 |
| 1634 | c19925_g1 | 1.426 | 9.40E-03 |
| 1635 | c57841_g1 | 1.427 | 1.57E-02 |
| 1636 | c55820_g1 | 1.427 | 9.81E-03 |
| 1637 | c42893_g1 | 1.432 | 1.11E-02 |
| 1638 | c54633_g1 | 1.444 | 1.88E-02 |
| 1639 | c53977_g1 | 1.450 | 7.22E-03 |
| 1640 | c53015_g1 | 1.452 | 6.72E-03 |
| 1641 | c52776_g1 | 1.463 | 8.87E-03 |
| 1642 | c49046_g1 | 1.465 | 1.62E-02 |
| 1643 | c54809_g2 | 1.476 | 1.23E-02 |
| 1644 | c48536_g1 | 1.482 | 1.68E-02 |
| 1645 | c29913_g1 | 1.483 | 8.92E-03 |
| 1646 | c55686_g3 | 1.485 | 8.43E-03 |
| 1647 | c50253_g1 | 1.489 | 7.08E-03 |
| 1648 | c36055_g1 | 1.490 | 1.13E-02 |
| 1649 | c45414_g1 | 1.495 | 9.32E-03 |
| 1650 | c40964_g1 | 1.498 | 9.92E-03 |
| 1651 | c42539_g2 | 1.500 | 7.20E-03 |
| 1652 | c50420_g1 | 1.507 | 7.31E-03 |

| | | | |
|---|---|---|---|
| 1653 | c51319_g1 | 1.511 | 5.38E-03 |
| 1654 | c54547_g1 | 1.512 | 5.43E-03 |
| 1655 | c49420_g1 | 1.513 | 7.56E-03 |
| 1656 | c33815_g1 | 1.513 | 6.55E-03 |
| 1657 | c46174_g1 | 1.516 | 7.71E-03 |
| 1658 | c43853_g1 | 1.516 | 1.20E-02 |
| 1659 | c53718_g1 | 1.520 | 1.38E-02 |
| 1660 | c37731_g1 | 1.520 | 1.06E-02 |
| 1661 | c47652_g1 | 1.522 | 1.76E-02 |
| 1662 | c50864_g1 | 1.531 | 5.70E-03 |
| 1663 | c53019_g1 | 1.539 | 6.82E-03 |
| 1664 | c34247_g1 | 1.543 | 4.86E-03 |
| 1665 | c55022_g1 | 1.546 | 8.67E-03 |
| 1666 | c53426_g5 | 1.553 | 6.30E-03 |
| 1667 | c51094_g1 | 1.563 | 4.51E-03 |
| 1668 | c51228_g2 | 1.563 | 6.82E-03 |
| 1669 | c10746_g1 | 1.581 | 5.23E-03 |
| 1670 | c109618_g1 | 1.592 | 4.53E-03 |
| 1671 | c48866_g1 | 1.600 | 3.59E-03 |
| 1672 | c48531_g1 | 1.602 | 3.65E-03 |
| 1673 | c23172_g1 | 1.607 | 7.72E-03 |
| 1674 | c45563_g1 | 1.611 | 4.39E-03 |
| 1675 | c57282_g4 | 1.611 | 6.08E-03 |
| 1676 | c71572_g1 | 1.616 | 4.75E-03 |
| 1677 | c9297_g1 | 1.616 | 3.77E-03 |
| 1678 | c57960_g1 | 1.626 | 2.51E-03 |
| 1679 | c53395_g1 | 1.632 | 5.09E-03 |
| 1680 | c57723_g1 | 1.633 | 2.27E-03 |
| 1681 | c50684_g1 | 1.636 | 4.32E-03 |
| 1682 | c54765_g5 | 1.639 | 9.06E-03 |
| 1683 | c44904_g1 | 1.643 | 5.26E-03 |
| 1684 | c18689_g1 | 1.646 | 3.85E-03 |
| 1685 | c58033_g1 | 1.647 | 6.13E-03 |
| 1686 | c56922_g1 | 1.647 | 3.72E-03 |
| 1687 | c41707_g1 | 1.654 | 3.09E-03 |
| 1688 | c47718_g1 | 1.662 | 2.01E-03 |
| 1689 | c44902_g1 | 1.670 | 2.03E-03 |
| 1690 | c32781_g1 | 1.676 | 2.36E-03 |
| 1691 | c52727_g1 | 1.677 | 2.66E-03 |
| 1692 | c36796_g1 | 1.680 | 4.47E-03 |
| 1693 | c42796_g1 | 1.681 | 5.66E-03 |
| 1694 | c52797_g1 | 1.681 | 2.71E-03 |
| 1695 | c53976_g1 | 1.685 | 4.60E-03 |
| 1696 | c54424_g4 | 1.689 | 3.46E-03 |
| 1697 | c49504_g1 | 1.694 | 2.64E-03 |
| 1698 | c51931_g1 | 1.696 | 4.14E-03 |
| 1699 | c10737_g1 | 1.698 | 6.91E-03 |
| 1700 | c56585_g1 | 1.711 | 1.54E-03 |
| 1701 | c49378_g1 | 1.713 | 1.95E-03 |
| 1702 | c58152_g1 | 1.719 | 2.42E-03 |
| 1703 | c46374_g1 | 1.721 | 4.83E-03 |
| 1704 | c32558_g1 | 1.722 | 3.53E-03 |
| 1705 | c57173_g1 | 1.737 | 6.38E-03 |
| 1706 | c39444_g1 | 1.742 | 2.09E-03 |
| 1707 | c54013_g1 | 1.748 | 2.55E-03 |
| 1708 | c50501_g1 | 1.756 | 1.94E-03 |
| 1709 | c23940_g1 | 1.758 | 3.47E-03 |
| 1710 | c45882_g1 | 1.759 | 1.15E-03 |
| 1711 | c52162_g1 | 1.767 | 1.80E-03 |
| 1712 | c33415_g1 | 1.785 | 1.63E-03 |
| 1713 | c46084_g2 | 1.787 | 1.37E-03 |
| 1714 | c49025_g1 | 1.789 | 2.75E-03 |
| 1715 | c22919_g1 | 1.794 | 2.48E-03 |
| 1716 | c49559_g1 | 1.807 | 1.67E-03 |
| 1717 | c23689_g1 | 1.807 | 6.34E-03 |
| 1718 | c54844_g1 | 1.810 | 1.18E-03 |
| 1719 | c41890_g1 | 1.816 | 1.47E-03 |
| 1720 | c37476_g1 | 1.821 | 1.36E-03 |
| 1721 | c51148_g1 | 1.823 | 9.77E-04 |

| | | | |
|---|---|---|---|
| 1722 | c52863_g1 | 1.827 | 4.53E-03 |
| 1723 | c52291_g1 | 1.831 | 1.15E-03 |
| 1724 | c56823_g2 | 1.834 | 1.45E-03 |
| 1725 | c55213_g1 | 1.848 | 1.66E-03 |
| 1726 | c58151_g4 | 1.854 | 8.91E-03 |
| 1727 | c47506_g1 | 1.858 | 1.23E-03 |
| 1728 | c29222_g1 | 1.859 | 7.84E-04 |
| 1729 | c58072_g6 | 1.866 | 7.32E-04 |
| 1730 | c57184_g1 | 1.867 | 5.99E-04 |
| 1731 | c54228_g1 | 1.869 | 1.17E-03 |
| 1732 | c57966_g1 | 1.869 | 6.16E-04 |
| 1733 | c55006_g2 | 1.871 | 5.42E-04 |
| 1734 | c57392_g1 | 1.878 | 1.09E-03 |
| 1735 | c54576_g1 | 1.880 | 1.05E-03 |
| 1736 | c55789_g1 | 1.896 | 6.44E-04 |
| 1737 | c84246_g1 | 1.907 | 4.89E-04 |
| 1738 | c47095_g1 | 1.911 | 1.74E-03 |
| 1739 | c56595_g2 | 1.916 | 4.88E-04 |
| 1740 | c53476_g1 | 1.927 | 4.88E-04 |
| 1741 | c12732_g1 | 1.928 | 4.27E-04 |
| 1742 | c53468_g1 | 1.931 | 8.21E-04 |
| 1743 | c50877_g1 | 1.933 | 8.01E-04 |
| 1744 | c57655_g3 | 1.937 | 6.94E-04 |
| 1745 | c47210_g1 | 1.945 | 7.42E-04 |
| 1746 | c48555_g1 | 1.947 | 1.87E-02 |
| 1747 | c57957_g1 | 1.963 | 1.15E-03 |
| 1748 | c55208_g1 | 1.965 | 9.33E-04 |
| 1749 | c56636_g1 | 1.970 | 4.28E-04 |
| 1750 | c50534_g1 | 1.984 | 1.02E-03 |
| 1751 | c58119_g1 | 1.996 | 3.00E-04 |
| 1752 | c51999_g1 | 2.000 | 5.89E-04 |
| 1753 | c39331_g2 | 2.001 | 6.75E-04 |
| 1754 | c58057_g7 | 2.002 | 6.12E-04 |
| 1755 | c46473_g1 | 2.003 | 4.22E-04 |
| 1756 | c57572_g4 | 2.004 | 4.51E-04 |
| 1757 | c48549_g1 | 2.004 | 3.99E-04 |
| 1758 | c55784_g1 | 2.006 | 5.47E-04 |
| 1759 | c50327_g3 | 2.011 | 2.72E-04 |
| 1760 | c52010_g1 | 2.012 | 4.82E-03 |
| 1761 | c31842_g1 | 2.012 | 3.87E-04 |
| 1762 | c57664_g1 | 2.022 | 1.66E-03 |
| 1763 | c56307_g2 | 2.026 | 2.04E-04 |
| 1764 | c47933_g1 | 2.032 | 3.56E-04 |
| 1765 | c47923_g1 | 2.036 | 2.92E-04 |
| 1766 | c55911_g2 | 2.039 | 2.31E-04 |
| 1767 | c56456_g1 | 2.049 | 6.03E-04 |
| 1768 | c53392_g1 | 2.052 | 6.35E-04 |
| 1769 | c23671_g1 | 2.052 | 2.22E-04 |
| 1770 | c58103_g2 | 2.054 | 2.86E-04 |
| 1771 | c50407_g1 | 2.067 | 3.93E-04 |
| 1772 | c52247_g1 | 2.080 | 4.01E-04 |
| 1773 | c34056_g1 | 2.086 | 1.49E-04 |
| 1774 | c57592_g3 | 2.103 | 2.24E-04 |
| 1775 | c54149_g1 | 2.114 | 2.71E-04 |
| 1776 | c51628_g1 | 2.126 | 2.05E-03 |
| 1777 | c53085_g1 | 2.128 | 1.89E-04 |
| 1778 | c123192_g1 | 2.139 | 7.83E-04 |
| 1779 | c39927_g1 | 2.151 | 8.62E-04 |
| 1780 | c51920_g1 | 2.166 | 5.13E-04 |
| 1781 | c37676_g1 | 2.169 | 1.93E-03 |
| 1782 | c54212_g3 | 2.176 | 4.05E-04 |
| 1783 | c52859_g1 | 2.180 | 2.01E-04 |
| 1784 | c55244_g1 | 2.195 | 5.75E-05 |
| 1785 | c10803_g1 | 2.202 | 3.77E-04 |
| 1786 | c50000_g1 | 2.215 | 9.07E-05 |
| 1787 | c54371_g1 | 2.215 | 9.07E-05 |
| 1788 | c52205_g1 | 2.219 | 1.17E-04 |
| 1789 | c41099_g1 | 2.220 | 2.64E-04 |
| 1790 | c58202_g2 | 2.221 | 1.72E-03 |

| | | | |
|---|---|---|---|
| 1791 | c42080_g1 | 2.226 | 1.00E-04 |
| 1792 | c50448_g1 | 2.244 | 1.41E-04 |
| 1793 | c55257_g1 | 2.275 | 4.84E-04 |
| 1794 | c55271_g1 | 2.279 | 2.02E-04 |
| 1795 | c55993_g1 | 2.280 | 3.67E-05 |
| 1796 | c54102_g1 | 2.285 | 6.59E-05 |
| 1797 | c50070_g1 | 2.293 | 2.95E-05 |
| 1798 | c52632_g1 | 2.301 | 3.05E-04 |
| 1799 | c49958_g1 | 2.311 | 4.01E-05 |
| 1800 | c43032_g1 | 2.318 | 7.91E-05 |
| 1801 | c47839_g2 | 2.339 | 5.87E-05 |
| 1802 | c45491_g1 | 2.341 | 6.51E-05 |
| 1803 | c35129_g1 | 2.346 | 2.27E-04 |
| 1804 | c54886_g2 | 2.346 | 5.14E-05 |
| 1805 | c45168_g1 | 2.355 | 4.72E-03 |
| 1806 | c49417_g1 | 2.359 | 2.48E-05 |
| 1807 | c10186_g1 | 2.362 | 4.30E-03 |
| 1808 | c48671_g1 | 2.371 | 2.17E-03 |
| 1809 | c56647_g1 | 2.372 | 1.18E-04 |
| 1810 | c57329_g2 | 2.373 | 6.64E-05 |
| 1811 | c43034_g1 | 2.375 | 2.31E-05 |
| 1812 | c34548_g1 | 2.378 | 1.32E-05 |
| 1813 | c28634_g1 | 2.378 | 5.04E-04 |
| 1814 | c39173_g1 | 2.390 | 3.37E-05 |
| 1815 | c56866_g1 | 2.392 | 2.87E-05 |
| 1816 | c54926_g1 | 2.393 | 2.63E-05 |
| 1817 | c56428_g1 | 2.394 | 4.75E-05 |
| 1818 | c50832_g1 | 2.399 | 2.30E-05 |
| 1819 | c49777_g1 | 2.404 | 1.12E-04 |
| 1820 | c46983_g1 | 2.433 | 5.68E-05 |
| 1821 | c34377_g1 | 2.439 | 2.86E-05 |
| 1822 | c57781_g1 | 2.459 | 1.95E-05 |
| 1823 | c57181_g1 | 2.459 | 1.35E-05 |
| 1824 | c52241_g1 | 2.473 | 1.92E-05 |
| 1825 | c45053_g1 | 2.473 | 3.88E-05 |
| 1826 | c58054_g5 | 2.481 | 1.67E-05 |
| 1827 | c15726_g1 | 2.485 | 9.43E-06 |
| 1828 | c38563_g1 | 2.498 | 2.76E-05 |
| 1829 | c57742_g1 | 2.501 | 1.20E-05 |
| 1830 | c47409_g1 | 2.501 | 5.80E-05 |
| 1831 | c54341_g1 | 2.505 | 2.65E-05 |
| 1832 | c47791_g1 | 2.512 | 1.23E-05 |
| 1833 | c12925_g1 | 2.527 | 1.75E-05 |
| 1834 | c58307_g1 | 2.529 | 7.53E-04 |
| 1835 | c15822_g1 | 2.535 | 1.58E-05 |
| 1836 | c43737_g1 | 2.556 | 1.08E-05 |
| 1837 | c43850_g1 | 2.559 | 1.52E-05 |
| 1838 | c48870_g1 | 2.568 | 1.79E-02 |
| 1839 | c49792_g1 | 2.568 | 4.96E-05 |
| 1840 | c48768_g1 | 2.588 | 4.34E-04 |
| 1841 | c55562_g1 | 2.595 | 7.47E-06 |
| 1842 | c51998_g1 | 2.608 | 9.34E-06 |
| 1843 | c51422_g1 | 2.610 | 2.68E-05 |
| 1844 | c57236_g1 | 2.617 | 2.76E-06 |
| 1845 | c39318_g1 | 2.625 | 8.01E-06 |
| 1846 | c47300_g1 | 2.632 | 2.60E-05 |
| 1847 | c57655_g1 | 2.643 | 4.34E-06 |
| 1848 | c23742_g1 | 2.644 | 1.91E-06 |
| 1849 | c23863_g1 | 2.657 | 1.43E-04 |
| 1850 | c41090_g1 | 2.670 | 5.12E-06 |
| 1851 | c57453_g1 | 2.672 | 1.79E-06 |
| 1852 | c51282_g3 | 2.675 | 1.80E-06 |
| 1853 | c16054_g1 | 2.678 | 1.99E-03 |
| 1854 | c47642_g1 | 2.689 | 2.51E-05 |
| 1855 | c71388_g1 | 2.702 | 3.88E-06 |
| 1856 | c35133_g1 | 2.703 | 7.47E-06 |
| 1857 | c56455_g1 | 2.711 | 4.84E-06 |
| 1858 | c50492_g1 | 2.713 | 1.27E-03 |
| 1859 | c57566_g1 | 2.722 | 1.23E-06 |

| | | | |
|---|---|---|---|
| 1860 | c55256_g1 | 2.725 | 1.32E-06 |
| 1861 | c44892_g1 | 2.731 | 1.51E-06 |
| 1862 | c52193_g1 | 2.757 | 6.85E-07 |
| 1863 | c23571_g1 | 2.764 | 1.11E-06 |
| 1864 | c54276_g1 | 2.766 | 1.16E-06 |
| 1865 | c44096_g1 | 2.777 | 5.89E-06 |
| 1866 | c26184_g1 | 2.779 | 2.79E-03 |
| 1867 | c47674_g1 | 2.791 | 6.33E-06 |
| 1868 | c97754_g1 | 2.799 | 2.47E-06 |
| 1869 | c51691_g1 | 2.818 | 1.19E-06 |
| 1870 | c56941_g4 | 2.826 | 8.24E-06 |
| 1871 | c47667_g1 | 2.826 | 1.84E-06 |
| 1872 | c51193_g1 | 2.830 | 3.01E-06 |
| 1873 | c51563_g1 | 2.830 | 7.04E-03 |
| 1874 | c37765_g1 | 2.833 | 1.45E-06 |
| 1875 | c50715_g4 | 2.834 | 6.72E-06 |
| 1876 | c51118_g1 | 2.844 | 2.84E-06 |
| 1877 | c51686_g1 | 2.846 | 2.43E-06 |
| 1878 | c109632_g1 | 2.847 | 2.64E-06 |
| 1879 | c55812_g1 | 2.850 | 3.11E-05 |
| 1880 | c54765_g3 | 2.886 | 1.03E-06 |
| 1881 | c51406_g1 | 2.897 | 1.22E-06 |
| 1882 | c48994_g1 | 2.912 | 5.55E-06 |
| 1883 | c58163_g1 | 2.914 | 7.01E-04 |
| 1884 | c55099_g1 | 2.922 | 6.15E-06 |
| 1885 | c57912_g1 | 2.922 | 3.56E-06 |
| 1886 | c56860_g1 | 2.943 | 6.44E-07 |
| 1887 | c53165_g1 | 2.959 | 3.21E-07 |
| 1888 | c26072_g1 | 2.965 | 2.53E-05 |
| 1889 | c47541_g1 | 2.980 | 5.93E-07 |
| 1890 | c33027_g1 | 2.984 | 2.08E-04 |
| 1891 | c41871_g1 | 2.987 | 2.42E-07 |
| 1892 | c56204_g1 | 2.991 | 2.13E-07 |
| 1893 | c56643_g1 | 2.999 | 1.30E-07 |
| 1894 | c43561_g1 | 3.017 | 1.88E-07 |
| 1895 | c37440_g1 | 3.018 | 2.57E-07 |
| 1896 | c53973_g1 | 3.022 | 3.29E-06 |
| 1897 | c56974_g1 | 3.023 | 7.85E-07 |
| 1898 | c96874_g1 | 3.031 | 2.42E-06 |
| 1899 | c48522_g1 | 3.032 | 1.06E-07 |
| 1900 | c52846_g1 | 3.043 | 7.74E-07 |
| 1901 | c56072_g1 | 3.048 | 2.77E-06 |
| 1902 | c53327_g1 | 3.081 | 7.75E-08 |
| 1903 | c57306_g1 | 3.083 | 2.67E-07 |
| 1904 | c53469_g3 | 3.112 | 5.71E-07 |
| 1905 | c50620_g1 | 3.172 | 2.89E-07 |
| 1906 | c42211_g1 | 3.198 | 5.00E-03 |
| 1907 | c50607_g1 | 3.223 | 6.49E-08 |
| 1908 | c54682_g2 | 3.249 | 2.15E-07 |
| 1909 | c58109_g1 | 3.263 | 7.37E-08 |
| 1910 | c53315_g1 | 3.266 | 2.47E-08 |
| 1911 | c46003_g1 | 3.287 | 2.54E-08 |
| 1912 | c41073_g1 | 3.287 | 2.10E-07 |
| 1913 | c56439_g1 | 3.329 | 2.10E-08 |
| 1914 | c26871_g1 | 3.331 | 3.34E-08 |
| 1915 | c52568_g1 | 3.334 | 1.63E-08 |
| 1916 | c46910_g1 | 3.388 | 6.83E-08 |
| 1917 | c43825_g1 | 3.412 | 4.33E-08 |
| 1918 | c23807_g1 | 3.418 | 7.86E-09 |
| 1919 | c57649_g1 | 3.423 | 5.32E-09 |
| 1920 | c48936_g2 | 3.449 | 2.89E-06 |
| 1921 | c33687_g1 | 3.450 | 6.04E-09 |
| 1922 | c57755_g2 | 3.481 | 1.02E-09 |
| 1923 | c55761_g1 | 3.512 | 1.61E-07 |
| 1924 | c57466_g1 | 3.526 | 6.76E-04 |
| 1925 | c54339_g4 | 3.534 | 4.36E-08 |
| 1926 | c49810_g1 | 3.567 | 3.84E-08 |
| 1927 | c42884_g1 | 3.609 | 1.80E-03 |
| 1928 | c52721_g1 | 3.657 | 2.45E-05 |

| | | | |
|---|---|---|---|
| 1929 | c54830_g1 | 3.682 | 1.51E-07 |
| 1930 | c57988_g6 | 3.790 | 6.86E-08 |
| 1931 | c42832_g2 | 3.844 | 1.98E-08 |
| 1932 | c57197_g1 | 3.868 | 2.15E-09 |
| 1933 | c54424_g5 | 3.869 | 1.35E-09 |
| 1934 | c22881_g1 | 3.870 | 6.56E-03 |
| 1935 | c45384_g1 | 3.870 | 6.56E-03 |
| 1936 | c56289_g1 | 3.890 | 1.57E-07 |
| 1937 | c30879_g1 | 3.895 | 9.88E-10 |
| 1938 | c47907_g1 | 3.921 | 4.42E-11 |
| 1939 | c114314_g1 | 3.921 | 7.87E-11 |
| 1940 | c42285_g1 | 3.931 | 1.02E-07 |
| 1941 | c44214_g1 | 3.941 | 1.00E-07 |
| 1942 | c53664_g1 | 3.942 | 6.56E-03 |
| 1943 | c71324_g1 | 3.974 | 3.78E-09 |
| 1944 | c51557_g1 | 3.988 | 3.44E-08 |
| 1945 | c46527_g1 | 3.995 | 1.33E-10 |
| 1946 | c57908_g2 | 4.015 | 7.74E-11 |
| 1947 | c53733_g1 | 4.070 | 2.16E-07 |
| 1948 | c52934_g1 | 4.086 | 6.43E-13 |
| 1949 | c39874_g1 | 4.101 | 3.55E-12 |
| 1950 | c56415_g1 | 4.130 | 2.15E-08 |
| 1951 | c40063_g1 | 4.145 | 6.87E-12 |
| 1952 | c58037_g1 | 4.174 | 2.14E-10 |
| 1953 | c52696_g1 | 4.208 | 1.69E-11 |
| 1954 | c54919_g1 | 4.225 | 2.40E-11 |
| 1955 | c49167_g1 | 4.292 | 2.94E-09 |
| 1956 | c14276_g1 | 4.343 | 1.86E-08 |
| 1957 | c51761_g1 | 4.380 | 1.78E-07 |
| 1958 | c57185_g8 | 4.421 | 6.17E-03 |
| 1959 | c51941_g1 | 4.427 | 7.46E-11 |
| 1960 | c56286_g1 | 4.543 | 2.75E-04 |
| 1961 | c14760_g1 | 4.615 | 3.90E-05 |
| 1962 | c40774_g2 | 4.625 | 2.66E-09 |
| 1963 | c116579_g1 | 4.645 | 4.08E-07 |
| 1964 | c50147_g1 | 4.753 | 3.15E-13 |
| 1965 | c57098_g1 | 4.758 | 5.51E-11 |
| 1966 | c20831_g1 | 4.771 | 2.74E-11 |
| 1967 | c56730_g2 | 5.100 | 1.38E-15 |
| 1968 | c57577_g7 | 5.178 | 1.95E-14 |
| 1969 | c47944_g2 | 5.241 | 1.03E-14 |
| 1970 | c48876_g1 | 5.242 | 1.50E-11 |
| 1971 | c87135_g1 | 5.354 | 1.48E-09 |
| 1972 | c103700_g1 | 5.369 | 3.08E-20 |
| 1973 | c39138_g1 | 5.442 | 1.48E-15 |
| 1974 | c23373_g1 | 5.561 | 2.60E-11 |
| 1975 | c53753_g1 | 5.652 | 9.73E-16 |
| 1976 | c55948_g2 | 6.016 | 1.44E-02 |
| 1977 | c110324_g1 | 6.129 | 6.03E-19 |
| 1978 | c52530_g1 | 6.278 | 3.97E-08 |
| 1979 | c44994_g1 | 6.676 | 9.27E-18 |
| 1980 | c55058_g1 | 6.744 | 1.26E-03 |
| 1981 | c49874_g1 | 6.811 | 1.49E-17 |
| 1982 | c57242_g4 | 6.911 | 1.79E-17 |
| 1983 | c43804_g1 | 7.005 | 2.91E-04 |
| 1984 | c53455_g1 | 7.013 | 2.49E-11 |
| 1985 | c52925_g1 | 7.417 | 3.90E-05 |
| 1986 | c56919_g1 | 7.828 | 1.72E-15 |
| 1987 | c20195_g1 | 8.582 | 7.17E-09 |
| 1988 | c53122_g2 | 9.455 | 3.07E-13 |
| 1989 | c56947_g1 | 9.658 | 3.00E-14 |
| 1990 | c56744_g1 | 9.934 | 8.16E-16 |
| 1991 | c57839_g1 | 10.054 | 1.98E-16 |
