## Supplementary material for "Decoding the reproductive system of the olive fruit fly, *Bactrocera oleae*"

| | | *B. oleae* transcriptome<br>Tissue: female lower reproductive track | | | |
|---|---|---|---|---|---|
| N | transcript_id | Annotation name | gene_id | logFC | PValue |
| 1 | c52845_g1 | *neprilysin 2* | NW_013581321.1.34 | -15.445 | 1.41E-14 |
| 2 | c55628_g1 | *Neural Lazarillo* | NW_013581373.1.1 | -15.097 | 1.48E-13 |
| 3 | c42759_g2 | *troponin-C* | NW_013581215.1.48 | -14.991 | 3.01E-13 |
| 4 | c51837_g1 | *cryptocephal* | NW_013583142.1.1 | -14.886 | 6.09E-13 |
| 5 | c53966_g8 | *midlin fascelin* | NW_013581231.1.21 | -14.761 | 1.05E-12 |
| 6 | c44387_g1 | *CG34189-like* | NW_013583061.1.3 | -14.744 | 1.18E-12 |
| 7 | c30526_g1 | *Ribosomal protein S16* | NW_013581356.1.32 | -14.608 | 2.90E-12 |
| 8 | c53346_g1 | *yolk protein 2* | NW_013581230.1.18 | -14.608 | 2.90E-12 |
| 9 | c47567_g2 | *rudimentary-like* | NW_013581298.1.19 | -14.424 | 9.85E-12 |
| 10 | c71357_g1 | *CG6426-like* | NW_013581234.1.7 | -14.331 | 1.84E-11 |
| 11 | c53746_g2 | *Imaginal disc growth factor 3* | NW_013581222.1.28 | -14.301 | 2.24E-11 |
| 12 | c53092_g2 | *Ornithine decarboxylase antizyme* | NW_013581383.1.6 | -14.298 | 2.28E-11 |
| 13 | c54618_g1 | *starvin* | NW_013581477.1.2 | -14.271 | 2.72E-11 |
| 14 | c47917_g2 | *Cyp6g1* | NW_013581325.1.20 | -14.195 | 3.31E-11 |
| 15 | c53040_g1 | *Ecdysone-inducible gene L3* | NW_013582894.1.2 | -14.188 | 3.48E-11 |
| 16 | c54829_g2 | *serpin42da* | NW_013581585.1.8 | -14.176 | 3.77E-11 |
| 17 | c57143_g2 | *Vacuolar H+-ATPase 55kD subunit* | NW_013581338.1.3 | -14.107 | 5.92E-11 |
| 18 | c55918_g1 | *CG14210-like* | NW_013581232.1.22 | -14.060 | 8.10E-11 |
| 19 | c52436_g1 | *scylla* | NW_013581315.1.11 | -14.005 | 1.16E-10 |
| 20 | c21198_g1 | *CG9676-like* | 6587_t | -13.942 | 1.74E-10 |
| 21 | c53599_g1 | *Secreted protein, acidic, cysteine-rich* | NW_013581490.1.3 | -13.940 | 1.76E-10 |
| 22 | c51323_g1 | not predicted | | -13.871 | 2.76E-10 |
| 23 | c27262_g1 | *ERp60* | NW_013581448.1.6 | -13.871 | 2.76E-10 |
| 24 | c10647_g1 | *ribosomal protein S17* | NW_013582719.1.1 | -13.869 | 2.80E-10 |
| 25 | c56768_g3 | *CG6770-like* | 3377_t | -13.867 | 2.85E-10 |
| 26 | c45452_g1 | *thin* | NW_013581209.1.155 | -13.780 | 3.67E-10 |
| 27 | c29117_g1 | *defensin* | NW_013581584.1.4 | -13.750 | 4.48E-10 |
| 28 | c44594_g1 | *CG7265-like* | NW_013581239.1.34 | -13.725 | 5.25E-10 |
| 29 | c58085_g1 | *megalin* | NW_013581232.1.2 | -13.692 | 6.54E-10 |
| 30 | c52934_g1 | not predicted | | -13.678 | 7.10E-10 |
| 31 | c57634_g1 | *CG10178-like* | NW_013581223.1.78 | -13.660 | 8.00E-10 |
| 32 | c58253_g1 | *cathD* | NW_013581226.1.24 | -13.590 | 1.26E-09 |
| 33 | c51907_g4 | *ribosomal protein L21* | NW_013581210.1.127 | -13.565 | 1.47E-09 |
| 34 | c32492_g1 | *ribosomal protein L23* | NW_013581313.1.21 | -13.555 | 1.57E-09 |
| 35 | c52931_g1 | *CG9747-like* | NW_013581266.1.30 | -13.521 | 1.96E-09 |
| 36 | c23076_g1 | *NADH dehydrogenase 13 kDa B subunit* | NW_013582233.1.1 | -13.469 | 1.99E-09 |
| 37 | c46559_g1 | *knockdown* | NW_013581243.1.12 | -13.425 | 2.64E-09 |
| 38 | c42417_g1 | *prip* | NW_013581220.1.123 | -13.420 | 2.73E-09 |
| 39 | c44954_g1 | *Cuticular protein 57A* | NW_013583188.1.8 | -13.352 | 4.22E-09 |
| 40 | c44953_g1 | *CG2852-like* | NW_013593435.1.1 | -13.338 | 4.60E-09 |
| 41 | c43875_g1 | *Vacuolar H+-ATPase SFD subunit* | NW_013581318.1.19 | -13.334 | 4.69E-09 |
| 42 | c49340_g1 | *Gelsolin* | NW_013581416.1.11 | -13.307 | 5.58E-09 |
| 43 | c54856_g2 | *Ubiquinol-cytochrome c reductase core* | NW_013581287.1.15 | -13.299 | 5.89E-09 |
| 44 | c51944_g2 | *refractory to sigma P* | NW_013586868.1.2 | -13.296 | 5.99E-09 |
| 45 | c50798_g2 | *ADP ribosylation factor at 79F* | NW_013581336.1.12 | -13.294 | 6.10E-09 |
| 46 | c50481_g1 | *Cytochrome P450-4g1* | NW_013581572.1.3 | -13.239 | 8.61E-09 |
| 47 | c10600_g2 | *Ribosomal protein S14a* | NW_013581258.1.15 | -13.238 | 8.69E-09 |
| 48 | c96905_g1 | *tryptophanyl-tRNA synthetase* | 21294_t | -13.218 | 9.80E-09 |
| 49 | c50073_g1 | *translation elongation factor 1 alpha 2* | NW_013581377.1.9 | -13.195 | 1.14E-08 |
| 50 | c54629_g1 | *Histone H4 replacement* | NW_013581368.1.23 | -13.181 | 1.24E-08 |
| 51 | c55864_g1 | not predicted | | -13.156 | 1.05E-08 |
| 52 | c57594_g1 | *CG42768-like* | NW_013581916.1.3 | -13.155 | 1.06E-08 |
| 53 | c54659_g1 | not predicted | | -13.148 | 1.10E-08 |
| 54 | c58082_g2 | *atlastin* | NW_013581457.1.10 | -13.142 | 1.15E-08 |
| 55 | c48434_g1 | *Rab1* | NW_013581293.1.12 | -13.126 | 1.27E-08 |
| 56 | c84110_g1 | *Heat shock protein cognate 5-RA* | NW_013581313.1.14 | -13.116 | 1.36E-08 |
| 57 | c53327_g2 | *N-m-D-a receptor-associated protein* | NW_013581247.1.40 | -13.059 | 1.94E-08 |
| 58 | c56913_g5 | *Protein kinase regulatory subunit type 1* | NW_013581359.1.26 | -13.048 | 2.09E-08 |
| 59 | c49653_g2 | *Rho1* | NW_013581209.1.264 | -13.046 | 2.11E-08 |

| | | | | | |
|---|---|---|---|---|---|
| 60 | c56580_g1 | *CG4276-like* | NW_013583226.1.1 | -13.019 | 2.49E-08 |
| 61 | c46189_g1 | *purity of essence* | NW_013581503.1.2 | -13.006 | 2.72E-08 |
| 62 | c49765_g1 | *CG9572-like* | NW_013581212.1.180 | -12.984 | 3.12E-08 |
| 63 | c55509_g7 | *Argonaute 2* | NW_013581786.1.1 | -12.971 | 3.37E-08 |
| 64 | c58054_g1 | *CG8177-like* | NW_013581302.1.29 | -12.955 | 3.72E-08 |
| 65 | c56851_g2 | *CG13776-like* | NW_013581210.1.73 | -12.929 | 4.35E-08 |
| 66 | c46581_g1 | *coro* | NW_013581849.1.8 | -12.925 | 4.50E-08 |
| 67 | c53123_g1 | *Mitochondrial assembly regulatory factor* | NW_013581256.1.7 | -12.895 | 5.39E-08 |
| 68 | c55319_g1 | *CG12702-like* | NW_013581246.1.53 | -12.889 | 5.58E-08 |
| 69 | c46085_g1 | *CG12605-like* | NW_013581221.1.123 | -12.887 | 5.65E-08 |
| 70 | c57012_g1 | *calnexin 99a* | NW_013581348.1.10 | -12.879 | 5.91E-08 |
| 71 | c57266_g3 | *CG7115-like* | NW_013581251.1.23 | -12.874 | 6.12E-08 |
| 72 | c27268_g1 | *Adenylosuccinate Synthetase* | NW_013581262.1.41 | -12.864 | 4.68E-08 |
| 73 | c49662_g2 | *Neural conserved at 73EF* | NW_013581423.1.20 | -12.851 | 5.08E-08 |
| 74 | c52754_g1 | *Gelsolin-isoform B* | NW_013581310.1.24 | -12.810 | 6.61E-08 |
| 75 | c42687_g1 | *CG10576-like* | NW_013581296.1.14 | -12.796 | 7.11E-08 |
| 76 | c42956_g1 | *Sterile20-like kinase* | NW_013581314.1.26 | -12.793 | 7.29E-08 |
| 77 | c27318_g1 | *CG6236-like* | NW_013581321.1.8 | -12.768 | 8.45E-08 |
| 78 | c43468_g1 | not predicted | | -12.740 | 1.01E-07 |
| 79 | c50627_g1 | *Imaginal disc growth factor 1* | NW_013581222.1.30 | -12.716 | 1.17E-07 |
| 80 | c48875_g2 | *Glutamine synthetase 2* | NW_013581268.1.16 | -12.708 | 1.24E-07 |
| 81 | c47214_g1 | *Nimrod B2* | NW_013584023.1.3 | -12.704 | 1.27E-07 |
| 82 | c12311_g1 | *Tudor staphylococcal nuclease* | NW_013581296.1.10 | -12.693 | 1.35E-07 |
| 83 | c54043_g1 | *mushroom-body expressed* | NW_013581465.1.2 | -12.667 | 1.58E-07 |
| 84 | c55085_g1 | not predicted | | -12.666 | 1.58E-07 |
| 85 | c53746_g1 | *Imaginal disc growth factor 3* | NW_013581222.1.28 | -12.659 | 1.67E-07 |
| 86 | c23095_g1 | *intronic protein 259* | NW_013581346.1.1 | -12.645 | 1.81E-07 |
| 87 | c46280_g1 | *vacuolar H[+] ATPase PPA1 subunit 1* | NW_013581485.1.9 | -12.645 | 1.81E-07 |
| 88 | c51868_g1 | *tetracycline resistance* | NW_013581225.1.87 | -12.625 | 2.04E-07 |
| 89 | c47782_g1 | *sugarless* | NW_013581551.1.6 | -12.617 | 2.16E-07 |
| 90 | c54301_g1 | *prolyl-4-hydroxylase-alpha SG1* | NW_013583024.1.4 | -12.594 | 1.77E-07 |
| 91 | c42523_g1 | *odorant-binding protein 99c* | NW_013581311.1.25 | -12.594 | 1.77E-07 |
| 92 | c50891_g1 | *Rab7* | NW_013581264.1.50 | -12.590 | 1.82E-07 |
| 93 | c57564_g1 | *Hexokinase C* | NW_013581215.1.93 | -12.563 | 2.12E-07 |
| 94 | c57765_g3 | *multiple ankyrin repeats single KH domain* | NW_013581297.1.9 | -12.553 | 2.25E-07 |
| 95 | c52177_g1 | *CG33203-like* | NW_013581264.1.62 | -12.552 | 2.28E-07 |
| 96 | c50839_g1 | *kayak* | NW_013581264.1.69 | -12.536 | 2.52E-07 |
| 97 | c52840_g1 | *Akt1* | NW_013581333.1.4 | -12.518 | 2.79E-07 |
| 98 | c36177_g1 | *Novel nucleolar protein 1* | NW_013581226.1.40 | -12.452 | 4.16E-07 |
| 99 | c54520_g1 | *lipophorin receptor 1* | NW_013581259.1.9 | -12.444 | 4.35E-07 |
| 100 | c57577_g2 | CG8839-like | NW_013581458.1.13 | -12.417 | 5.15E-07 |
| 101 | c47718_g1 | | | -12.441 | 1.24E-05 |
| 102 | c57577_g2 | | | -12.417 | 1.42E-05 |
| 103 | c57050_g2 | | | -12.386 | 1.65E-05 |
| 104 | c31514_g1 | | | -12.385 | 1.65E-05 |
| 105 | c55621_g1 | | | -12.378 | 1.71E-05 |
| 106 | c57575_g2 | | | -12.365 | 1.83E-05 |
| 107 | c47020_g1 | | | -12.346 | 1.53E-05 |
| 108 | c54013_g3 | | | -12.334 | 1.60E-05 |
| 109 | c47106_g2 | | | -12.317 | 1.72E-05 |
| 110 | c57856_g1 | | | -12.303 | 1.85E-05 |
| 111 | c52600_g1 | | | -12.278 | 2.13E-05 |
| 112 | c43569_g1 | | | -12.256 | 2.39E-05 |
| 113 | c56278_g2 | | | -12.255 | 2.39E-05 |
| 114 | c8808_g1 | | | -12.254 | 2.41E-05 |
| 115 | c53801_g1 | | | -12.247 | 2.47E-05 |
| 116 | c50743_g1 | | | -12.245 | 2.48E-05 |
| 117 | c56753_g1 | | | -12.243 | 2.48E-05 |
| 118 | c46274_g1 | | | -12.238 | 2.53E-05 |
| 119 | c8815_g1 | | | -12.233 | 2.59E-05 |
| 120 | c54746_g6 | | | -12.232 | 2.59E-05 |
| 121 | c56779_g2 | | | -12.226 | 2.66E-05 |

| | | | |
|---|---|---|---|
| 122 | c40827_g1 | -12.212 | 2.88E-05 |
| 123 | c51710_g1 | -12.211 | 2.88E-05 |
| 124 | c53296_g1 | -12.208 | 2.95E-05 |
| 125 | c47323_g1 | -12.203 | 2.99E-05 |
| 126 | c33421_g1 | -12.190 | 3.19E-05 |
| 127 | c96851_g1 | -12.158 | 3.72E-05 |
| 128 | c55329_g1 | -12.158 | 3.70E-05 |
| 129 | c50429_g1 | -12.158 | 3.72E-05 |
| 130 | c53244_g1 | -12.154 | 3.72E-05 |
| 131 | c43092_g1 | -12.153 | 3.76E-05 |
| 132 | c56594_g1 | -12.152 | 3.76E-05 |
| 133 | c26938_g1 | -12.144 | 3.87E-05 |
| 134 | c43056_g1 | -12.141 | 3.92E-05 |
| 135 | c58017_g2 | -12.137 | 4.05E-05 |
| 136 | c39949_g1 | -12.126 | 4.24E-05 |
| 137 | c51284_g1 | -12.121 | 4.38E-05 |
| 138 | c56063_g3 | -12.105 | 3.67E-05 |
| 139 | c53812_g1 | -12.103 | 3.67E-05 |
| 140 | c51907_g1 | -12.102 | 3.70E-05 |
| 141 | c48898_g1 | -12.102 | 3.67E-05 |
| 142 | c48550_g2 | -12.097 | 3.72E-05 |
| 143 | c41090_g1 | -12.095 | 3.72E-05 |
| 144 | c46953_g1 | -12.094 | 3.72E-05 |
| 145 | c44589_g1 | -12.093 | 3.72E-05 |
| 146 | c51247_g1 | -12.086 | 3.87E-05 |
| 147 | c57693_g1 | -12.070 | 4.20E-05 |
| 148 | c46698_g1 | -12.048 | 4.66E-05 |
| 149 | c34532_g1 | -12.046 | 4.73E-05 |
| 150 | c52861_g2 | -12.040 | 4.90E-05 |
| 151 | c56741_g1 | -12.031 | 5.05E-05 |
| 152 | c47790_g1 | -12.024 | 5.23E-05 |
| 153 | c37475_g1 | -12.019 | 5.42E-05 |
| 154 | c53388_g1 | -12.007 | 5.68E-05 |
| 155 | c55275_g2 | -12.006 | 5.68E-05 |
| 156 | c56555_g3 | -12.004 | 5.68E-05 |
| 157 | c57792_g1 | -12.004 | 5.68E-05 |
| 158 | c57914_g1 | -11.997 | 5.89E-05 |
| 159 | c25210_g1 | -11.990 | 6.11E-05 |
| 160 | c56205_g3 | -11.988 | 6.18E-05 |
| 161 | c10618_g1 | -11.987 | 6.18E-05 |
| 162 | c55863_g1 | -11.985 | 6.28E-05 |
| 163 | c56656_g2 | -11.980 | 6.36E-05 |
| 164 | c53758_g3 | -11.974 | 6.60E-05 |
| 165 | c51418_g1 | -11.961 | 7.00E-05 |
| 166 | c45998_g1 | -11.960 | 7.00E-05 |
| 167 | c58077_g1 | -11.959 | 7.00E-05 |
| 168 | c53741_g1 | -11.958 | 7.00E-05 |
| 169 | c55326_g1 | -11.952 | 7.23E-05 |
| 170 | c53734_g1 | -11.945 | 7.45E-05 |
| 171 | c26940_g1 | -11.945 | 7.45E-05 |
| 172 | c38209_g1 | -11.935 | 7.73E-05 |
| 173 | c52656_g2 | -11.934 | 7.73E-05 |
| 174 | c42841_g1 | -11.925 | 8.09E-05 |
| 175 | c45726_g1 | -11.917 | 8.40E-05 |
| 176 | c55000_g2 | -11.911 | 8.49E-05 |
| 177 | c16060_g1 | -11.908 | 8.56E-05 |
| 178 | c43839_g1 | -11.904 | 8.64E-05 |
| 179 | c58086_g1 | -11.901 | 8.77E-05 |
| 180 | c58369_g1 | -11.896 | 9.04E-05 |
| 181 | c56114_g2 | -11.887 | 7.22E-05 |
| 182 | c56105_g1 | -11.886 | 7.32E-05 |
| 183 | c11918_g1 | -11.881 | 7.56E-05 |

| | | | |
|---|---|---|---|
| 184 | c56435_g1 | -11.878 | 7.69E-05 |
| 185 | c57563_g2 | -11.873 | 7.73E-05 |
| 186 | c55406_g1 | -11.872 | 7.73E-05 |
| 187 | c47822_g1 | -11.871 | 7.73E-05 |
| 188 | c55697_g1 | -11.868 | 7.86E-05 |
| 189 | c53700_g1 | -11.864 | 8.00E-05 |
| 190 | c50073_g2 | -11.863 | 8.09E-05 |
| 191 | c50396_g3 | -11.862 | 8.09E-05 |
| 192 | c52661_g1 | -11.852 | 8.40E-05 |
| 193 | c52717_g2 | -11.851 | 8.49E-05 |
| 194 | c53977_g1 | -11.848 | 8.49E-05 |
| 195 | c36219_g1 | -11.848 | 8.56E-05 |
| 196 | c45281_g1 | -11.846 | 8.56E-05 |
| 197 | c23641_g1 | -11.844 | 8.56E-05 |
| 198 | c49783_g2 | -11.842 | 8.66E-05 |
| 199 | c50748_g1 | -11.840 | 8.77E-05 |
| 200 | c84144_g1 | -11.839 | 8.77E-05 |
| 201 | c40681_g2 | -11.831 | 9.04E-05 |
| 202 | c31463_g1 | -11.830 | 9.04E-05 |
| 203 | c49086_g2 | -11.829 | 9.04E-05 |
| 204 | c49625_g2 | -11.827 | 9.18E-05 |
| 205 | c38927_g1 | -11.826 | 9.18E-05 |
| 206 | c54608_g1 | -11.823 | 9.36E-05 |
| 207 | c57314_g1 | -11.819 | 9.45E-05 |
| 208 | c54078_g1 | -11.818 | 9.45E-05 |
| 209 | c44374_g1 | -11.817 | 9.45E-05 |
| 210 | c43515_g1 | -11.816 | 9.60E-05 |
| 211 | c55265_g1 | -11.810 | 9.78E-05 |
| 212 | c55179_g1 | -11.807 | 9.97E-05 |
| 213 | c54316_g1 | -11.805 | 0.0001017 |
| 214 | c55607_g1 | -11.800 | 0.0001036 |
| 215 | c46567_g1 | -11.790 | 0.0001081 |
| 216 | c55648_g1 | -11.788 | 0.0001098 |
| 217 | c57480_g1 | -11.786 | 0.0001098 |
| 218 | c51784_g1 | -11.780 | 0.0001142 |
| 219 | c58060_g1 | -11.773 | 0.0001165 |
| 220 | c49066_g1 | -11.772 | 0.0001188 |
| 221 | c57875_g1 | -11.767 | 0.0001208 |
| 222 | c24207_g1 | -11.767 | 0.0001208 |
| 223 | c71416_g1 | -11.762 | 0.0001232 |
| 224 | c45991_g5 | -11.760 | 0.0001249 |
| 225 | c25344_g1 | -11.757 | 0.0001249 |
| 226 | c55186_g1 | -11.754 | 0.0001271 |
| 227 | c51856_g1 | -11.753 | 0.0001271 |
| 228 | c53461_g1 | -11.749 | 0.0001289 |
| 229 | c50756_g1 | -11.746 | 0.0001307 |
| 230 | c56319_g1 | -11.746 | 0.0001307 |
| 231 | c54168_g1 | -11.746 | 0.0001307 |
| 232 | c54648_g1 | -11.742 | 0.0001334 |
| 233 | c49519_g2 | -11.738 | 0.000136 |
| 234 | c58165_g1 | -11.734 | 0.0001381 |
| 235 | c52129_g1 | -11.727 | 0.0001438 |
| 236 | c57877_g1 | -11.718 | 0.0001477 |
| 237 | c56786_g4 | -11.718 | 0.0001477 |
| 238 | c39536_g1 | -11.718 | 0.0001477 |
| 239 | c56891_g1 | -11.717 | 0.0001477 |
| 240 | c42721_g1 | -11.707 | 0.0001572 |
| 241 | c54044_g1 | -11.704 | 0.0001572 |
| 242 | c57563_g1 | -11.701 | 0.0001601 |
| 243 | c51362_g1 | -11.698 | 0.0001626 |
| 244 | c55522_g1 | -11.689 | 0.0001686 |
| 245 | c47397_g1 | -11.688 | 0.0001686 |

| | | | |
|---|---|---|---|
| 246 | c50433_g2 | -11.686 | 0.0001686 |
| 247 | c26676_g1 | -11.686 | 0.0001713 |
| 248 | c57241_g1 | -11.685 | 0.0001713 |
| 249 | c57125_g1 | -11.683 | 0.0001713 |
| 250 | c49411_g1 | -11.680 | 0.0001736 |
| 251 | c57386_g11 | -11.679 | 0.0001736 |
| 252 | c37540_g1 | -11.677 | 0.000177 |
| 253 | c50662_g1 | -11.662 | 0.0001426 |
| 254 | c84123_g1 | -11.659 | 0.0001448 |
| 255 | c50703_g1 | -11.658 | 0.0001448 |
| 256 | c54032_g1 | -11.655 | 0.0001477 |
| 257 | c41105_g1 | -11.640 | 0.0001571 |
| 258 | c53755_g9 | -11.631 | 0.0001623 |
| 259 | c42476_g1 | -11.630 | 0.0001623 |
| 260 | c53066_g1 | -11.622 | 0.0001686 |
| 261 | c56350_g1 | -11.613 | 0.0001736 |
| 262 | c57088_g1 | -11.610 | 0.000177 |
| 263 | c47442_g1 | -11.596 | 0.0001894 |
| 264 | c53173_g1 | -11.595 | 0.0001894 |
| 265 | c53264_g2 | -11.589 | 0.0001937 |
| 266 | c53386_g2 | -11.583 | 0.0002033 |
| 267 | c34297_g1 | -11.575 | 0.0002074 |
| 268 | c55984_g1 | -11.575 | 0.0002122 |
| 269 | c122422_g1 | -11.570 | 0.0002172 |
| 270 | c17482_g1 | -11.560 | 0.0002275 |
| 271 | c56348_g10 | -11.560 | 0.0002275 |
| 272 | c49173_g1 | -11.554 | 0.0002328 |
| 273 | c52369_g1 | -11.547 | 0.0002433 |
| 274 | c51794_g1 | -11.544 | 0.0002433 |
| 275 | c46866_g1 | -11.543 | 0.0002433 |
| 276 | c56123_g1 | -11.542 | 0.0002484 |
| 277 | c47321_g2 | -11.539 | 0.0002484 |
| 278 | c55303_g1 | -11.529 | 0.0002604 |
| 279 | c52947_g1 | -11.529 | 0.0002604 |
| 280 | c56599_g1 | -11.525 | 0.000266 |
| 281 | c34247_g1 | -11.525 | 0.000266 |
| 282 | c40256_g1 | -11.523 | 0.0002717 |
| 283 | c46951_g2 | -11.522 | 0.0002717 |
| 284 | c46152_g1 | -11.515 | 0.0002762 |
| 285 | c57488_g12 | -11.515 | 0.0002762 |
| 286 | c50551_g1 | -11.503 | 0.000293 |
| 287 | c52627_g1 | -11.501 | 0.000293 |
| 288 | c55008_g1 | -11.501 | 0.000293 |
| 289 | c40909_g1 | -11.499 | 0.000293 |
| 290 | c49237_g1 | -11.496 | 0.0002997 |
| 291 | c46283_g1 | -11.494 | 0.0003048 |
| 292 | c57907_g1 | -11.485 | 0.0003111 |
| 293 | c57131_g1 | -11.483 | 0.000317 |
| 294 | c51319_g1 | -11.478 | 0.0003226 |
| 295 | c58181_g2 | -11.476 | 0.0003226 |
| 296 | c49441_g1 | -11.473 | 0.0003299 |
| 297 | c37945_g1 | -11.467 | 0.0003358 |
| 298 | c26783_g1 | -11.466 | 0.0003358 |
| 299 | c50926_g1 | -11.463 | 0.0003426 |
| 300 | c27309_g1 | -11.462 | 0.0003426 |
| 301 | c54998_g2 | -11.458 | 0.0003481 |
| 302 | c55753_g1 | -11.458 | 0.0003481 |
| 303 | c57592_g2 | -11.456 | 0.0003481 |
| 304 | c8482_g1 | -11.453 | 0.0003553 |
| 305 | c57012_g2 | -11.447 | 0.0003619 |
| 306 | c49935_g1 | -11.444 | 0.0003711 |
| 307 | c47589_g1 | -11.443 | 0.000282 |

| | | | |
|---|---|---|---|
| 308 | c55705_g1 | -11.436 | 0.0002892 |
| 309 | c55342_g1 | -11.431 | 0.000293 |
| 310 | c8759_g1 | -11.431 | 0.000293 |
| 311 | c55710_g1 | -11.429 | 0.0002999 |
| 312 | c57925_g1 | -11.419 | 0.0003111 |
| 313 | c56109_g1 | -11.418 | 0.0003111 |
| 314 | c50784_g1 | -11.418 | 0.0003111 |
| 315 | c34064_g1 | -11.415 | 0.0003111 |
| 316 | c37474_g1 | -11.413 | 0.0003177 |
| 317 | c32781_g1 | -11.410 | 0.0003177 |
| 318 | c25331_g1 | -11.405 | 0.0003245 |
| 319 | c39556_g1 | -11.403 | 0.0003315 |
| 320 | c49699_g1 | -11.401 | 0.0003315 |
| 321 | c27562_g2 | -11.394 | 0.0003387 |
| 322 | c44842_g1 | -11.390 | 0.0003453 |
| 323 | c54175_g1 | -11.388 | 0.0003512 |
| 324 | c57041_g1 | -11.387 | 0.0003512 |
| 325 | c55306_g1 | -11.383 | 0.0003582 |
| 326 | c57655_g1 | -11.379 | 0.0003582 |
| 327 | c24824_g1 | -11.378 | 0.0003582 |
| 328 | c54880_g1 | -11.371 | 0.0003743 |
| 329 | c57861_g1 | -11.370 | 0.0003743 |
| 330 | c96827_g1 | -11.367 | 0.0003743 |
| 331 | c58149_g1 | -11.363 | 0.0003836 |
| 332 | c51106_g1 | -11.362 | 0.0003836 |
| 333 | c56898_g1 | -11.359 | 0.0003922 |
| 334 | c35296_g1 | -11.359 | 0.0003922 |
| 335 | c51285_g1 | -11.351 | 0.0004029 |
| 336 | c55683_g1 | -11.350 | 0.0004129 |
| 337 | c53305_g1 | -11.348 | 0.0004129 |
| 338 | c30806_g1 | -11.340 | 0.0004331 |
| 339 | c57588_g1 | -11.336 | 0.0004331 |
| 340 | c55188_g1 | -11.336 | 0.0004331 |
| 341 | c53740_g1 | -11.335 | 0.0004331 |
| 342 | c26900_g1 | -11.335 | 0.0004422 |
| 343 | c49817_g1 | -11.332 | 0.0004422 |
| 344 | c55226_g1 | -11.332 | 0.0004422 |
| 345 | c56902_g3 | -11.329 | 0.0004422 |
| 346 | c56045_g1 | -11.324 | 0.000466 |
| 347 | c57966_g1 | -11.320 | 0.000466 |
| 348 | c46488_g2 | -11.319 | 0.000466 |
| 349 | c39708_g1 | -11.311 | 0.0004913 |
| 350 | c47734_g2 | -11.310 | 0.0004913 |
| 351 | c46428_g1 | -11.309 | 0.0004913 |
| 352 | c30202_g1 | -11.301 | 0.0005171 |
| 353 | c55662_g1 | -11.299 | 0.0005171 |
| 354 | c57651_g2 | -11.295 | 0.0005273 |
| 355 | c56400_g1 | -11.292 | 0.0005273 |
| 356 | c51717_g1 | -11.292 | 0.0005273 |
| 357 | c42567_g1 | -11.289 | 0.0005389 |
| 358 | c56595_g2 | -11.287 | 0.0005389 |
| 359 | c48604_g2 | -11.287 | 0.0005389 |
| 360 | c122672_g1 | -11.284 | 0.0005475 |
| 361 | c54861_g1 | -11.280 | 0.0005475 |
| 362 | c10705_g1 | -11.276 | 0.000562 |
| 363 | c53851_g2 | -11.272 | 0.0005747 |
| 364 | c52935_g1 | -11.268 | 0.0005747 |
| 365 | c55026_g1 | -11.265 | 0.0005854 |
| 366 | c56893_g1 | -11.263 | 0.0005854 |
| 367 | c55462_g1 | -11.260 | 0.0005976 |
| 368 | c54836_g2 | -11.254 | 0.000609 |
| 369 | c57217_g1 | -11.253 | 0.000609 |

| | | | |
|---|---|---|---|
| 370 | c54488_g2 | -11.253 | 0.000609 |
| 371 | c48819_g1 | -11.252 | 0.000609 |
| 372 | c52120_g1 | -11.249 | 0.0006195 |
| 373 | c58342_g1 | -11.246 | 0.0006195 |
| 374 | c51973_g1 | -11.245 | 0.0006195 |
| 375 | c46160_g1 | -11.245 | 0.0006352 |
| 376 | c56002_g1 | -11.244 | 0.0006352 |
| 377 | c55680_g1 | -11.238 | 0.0006453 |
| 378 | c53091_g1 | -11.236 | 0.0006453 |
| 379 | c55917_g1 | -11.234 | 0.0006453 |
| 380 | c10659_g1 | -11.233 | 0.0006594 |
| 381 | c71595_g1 | -11.232 | 0.0006594 |
| 382 | c54971_g3 | -11.232 | 0.0006594 |
| 383 | c56575_g1 | -11.227 | 0.0006764 |
| 384 | c46423_g1 | -11.226 | 0.0006764 |
| 385 | c56036_g1 | -11.225 | 0.0005189 |
| 386 | c46636_g1 | -11.224 | 0.0005189 |
| 387 | c71437_g1 | -11.213 | 0.0005413 |
| 388 | c57968_g1 | -11.210 | 0.0005413 |
| 389 | c57555_g1 | -11.203 | 0.0005531 |
| 390 | c46396_g1 | -11.200 | 0.0005675 |
| 391 | c25201_g1 | -11.199 | 0.0005675 |
| 392 | c55859_g1 | -11.198 | 0.0005789 |
| 393 | c55357_g1 | -11.194 | 0.0005789 |
| 394 | c57713_g1 | -11.192 | 0.0005918 |
| 395 | c48374_g1 | -11.190 | 0.0005918 |
| 396 | c57759_g1 | -11.184 | 0.0006063 |
| 397 | c44934_g1 | -11.182 | 0.0005918 |
| 398 | c42889_g1 | -11.180 | 0.0006063 |
| 399 | c46536_g1 | -11.179 | 0.0006165 |
| 400 | c45802_g1 | -11.178 | 0.0006165 |
| 401 | c37520_g1 | -11.176 | 0.0006165 |
| 402 | c57648_g1 | -11.174 | 0.0006165 |
| 403 | c55540_g3 | -11.167 | 0.000644 |
| 404 | c50533_g1 | -11.167 | 0.000644 |
| 405 | c55965_g3 | -11.167 | 0.000644 |
| 406 | c56485_g2 | -11.166 | 0.000644 |
| 407 | c35181_g1 | -11.165 | 0.000644 |
| 408 | c49648_g2 | -11.159 | 0.0006591 |
| 409 | c53718_g2 | -11.157 | 0.0006591 |
| 410 | c51611_g2 | -11.153 | 0.0006758 |
| 411 | c54910_g1 | -11.146 | 0.0006918 |
| 412 | c57211_g1 | -11.146 | 0.0006918 |
| 413 | c51653_g1 | -11.144 | 0.0006918 |
| 414 | c48353_g1 | -11.142 | 0.000711 |
| 415 | c55792_g1 | -11.142 | 0.000711 |
| 416 | c44620_g1 | -11.136 | 0.000732 |
| 417 | c48009_g1 | -11.132 | 0.000732 |
| 418 | c58058_g1 | -11.131 | 0.0007511 |
| 419 | c58017_g1 | -11.129 | 0.0007511 |
| 420 | c51017_g1 | -11.127 | 0.0007511 |
| 421 | c56751_g5 | -11.125 | 0.0007511 |
| 422 | c14616_g1 | -11.124 | 0.0007749 |
| 423 | c53952_g1 | -11.111 | 0.0008207 |
| 424 | c57065_g2 | -11.110 | 0.0008207 |
| 425 | c54090_g2 | -11.107 | 0.0008207 |
| 426 | c55095_g1 | -11.107 | 0.0008207 |
| 427 | c24239_g1 | -11.104 | 0.0008455 |
| 428 | c57583_g1 | -11.097 | 0.0008727 |
| 429 | c52100_g1 | -11.089 | 0.0008993 |
| 430 | c34110_g1 | -11.085 | 0.0009252 |
| 431 | c55691_g1 | -11.080 | 0.0009252 |

| | | | |
|---|---|---|---|
| 432 | c50386_g1 | -11.080 | 0.0009252 |
| 433 | c58170_g1 | -11.077 | 0.0009454 |
| 434 | c52816_g1 | -11.076 | 0.0009454 |
| 435 | c52905_g1 | -11.075 | 0.0009454 |
| 436 | c50190_g1 | -11.075 | 0.0009454 |
| 437 | c53947_g1 | -11.075 | 0.0009454 |
| 438 | c46469_g1 | -11.071 | 0.0009597 |
| 439 | c51782_g1 | -11.070 | 0.0009597 |
| 440 | c52891_g1 | -11.068 | 0.0009597 |
| 441 | c49895_g1 | -11.062 | 0.0009829 |
| 442 | c54340_g1 | -11.060 | 0.0009829 |
| 443 | c57237_g4 | -11.057 | 0.0010119 |
| 444 | c51693_g1 | -11.053 | 0.0010367 |
| 445 | c57329_g1 | -11.049 | 0.0010367 |
| 446 | c44309_g1 | -11.048 | 0.0010367 |
| 447 | c15579_g1 | -11.048 | 0.0010367 |
| 448 | c53693_g1 | -11.046 | 0.0010589 |
| 449 | c52391_g1 | -11.045 | 0.0010589 |
| 450 | c49404_g1 | -11.045 | 0.0010589 |
| 451 | c44902_g1 | -11.041 | 0.0010589 |
| 452 | c57366_g1 | -11.041 | 0.0010589 |
| 453 | c48540_g1 | -11.039 | 0.0010836 |
| 454 | c54170_g1 | -11.038 | 0.0010836 |
| 455 | c58064_g4 | -11.036 | 0.0010836 |
| 456 | c57508_g1 | -11.035 | 0.0010836 |
| 457 | c45020_g1 | -11.034 | 0.0010836 |
| 458 | c54685_g1 | -11.033 | 0.0011163 |
| 459 | c55320_g1 | -11.030 | 0.0011163 |
| 460 | c55929_g1 | -11.026 | 0.0011446 |
| 461 | c47502_g1 | -11.017 | 0.001172 |
| 462 | c51926_g1 | -11.015 | 0.001172 |
| 463 | c55175_g1 | -11.013 | 0.0012052 |
| 464 | c55853_g1 | -11.011 | 0.0012052 |
| 465 | c54696_g1 | -11.005 | 0.0009347 |
| 466 | c54018_g2 | -11.004 | 0.0009347 |
| 467 | c55959_g1 | -10.997 | 0.0009471 |
| 468 | c57988_g9 | -10.997 | 0.0009471 |
| 469 | c57776_g4 | -10.996 | 0.0009471 |
| 470 | c50372_g1 | -10.995 | 0.0009471 |
| 471 | c42532_g1 | -10.995 | 0.0009471 |
| 472 | c48551_g1 | -10.995 | 0.0009471 |
| 473 | c56900_g1 | -10.993 | 0.0009471 |
| 474 | c56988_g1 | -10.990 | 0.0009716 |
| 475 | c49064_g1 | -10.989 | 0.0009716 |
| 476 | c57100_g1 | -10.987 | 0.0009716 |
| 477 | c57764_g2 | -10.980 | 0.0010003 |
| 478 | c47034_g1 | -10.978 | 0.00103 |
| 479 | c50831_g1 | -10.978 | 0.0009471 |
| 480 | c57783_g1 | -10.972 | 0.00103 |
| 481 | c45303_g1 | -10.971 | 0.0010555 |
| 482 | c53055_g1 | -10.966 | 0.0010555 |
| 483 | c39573_g1 | -10.962 | 0.001082 |
| 484 | c54576_g1 | -10.960 | 0.001082 |
| 485 | c39584_g1 | -10.951 | 0.0011111 |
| 486 | c51997_g1 | -10.949 | 0.0011395 |
| 487 | c56792_g1 | -10.948 | 0.0011395 |
| 488 | c56024_g2 | -10.947 | 0.0011395 |
| 489 | c57178_g1 | -10.944 | 0.0011395 |
| 490 | c57944_g1 | -10.941 | 0.0011708 |
| 491 | c71475_g1 | -10.938 | 0.0011708 |
| 492 | c57311_g1 | -10.938 | 0.0011708 |
| 493 | c47027_g1 | -10.935 | 0.0012052 |

| | | | |
|---|---|---|---|
| 494 | c45371_g1 | -10.931 | 0.0012052 |
| 495 | c29222_g1 | -10.923 | 0.0012444 |
| 496 | c44190_g1 | -10.918 | 0.0012816 |
| 497 | c56503_g1 | -10.918 | 0.0012816 |
| 498 | c49748_g1 | -10.917 | 0.0012816 |
| 499 | c52471_g1 | -10.917 | 0.0012816 |
| 500 | c55692_g1 | -10.914 | 0.0013216 |
| 501 | c54225_g1 | -10.913 | 0.0013216 |
| 502 | c54671_g9 | -10.909 | 0.0013216 |
| 503 | c35070_g1 | -10.907 | 0.0013216 |
| 504 | c55293_g5 | -10.898 | 0.0014153 |
| 505 | c55825_g1 | -10.894 | 0.0014153 |
| 506 | c53684_g1 | -10.893 | 0.0014153 |
| 507 | c52162_g1 | -10.891 | 0.0014538 |
| 508 | c44570_g1 | -10.890 | 0.0014538 |
| 509 | c41097_g1 | -10.888 | 0.0014538 |
| 510 | c57608_g1 | -10.888 | 0.0014538 |
| 511 | c54268_g1 | -10.886 | 0.0014538 |
| 512 | c57227_g1 | -10.886 | 0.0014538 |
| 513 | c84044_g1 | -10.877 | 0.0015582 |
| 514 | c57140_g1 | -10.874 | 0.0015582 |
| 515 | c55565_g1 | -10.873 | 0.0015582 |
| 516 | c56336_g2 | -10.866 | 0.0016064 |
| 517 | c57728_g2 | -10.863 | 0.0016064 |
| 518 | c53447_g1 | -10.862 | 0.0016393 |
| 519 | c58237_g1 | -10.858 | 0.0016393 |
| 520 | c48980_g1 | -10.855 | 0.0016393 |
| 521 | c50238_g1 | -10.854 | 0.0016834 |
| 522 | c41895_g1 | -10.854 | 0.0016834 |
| 523 | c57540_g1 | -10.850 | 0.0016834 |
| 524 | c56270_g1 | -10.849 | 0.0016834 |
| 525 | c51849_g1 | -10.849 | 0.0016834 |
| 526 | c51916_g1 | -10.842 | 0.0017316 |
| 527 | c14157_g2 | -10.835 | 0.0017892 |
| 528 | c41081_g1 | -10.833 | 0.0017892 |
| 529 | c55667_g1 | -10.832 | 0.0018465 |
| 530 | c54679_g1 | -10.831 | 0.0018465 |
| 531 | c42211_g1 | -10.822 | 0.0019059 |
| 532 | c54578_g1 | -10.817 | 0.0019059 |
| 533 | c55509_g4 | -10.815 | 0.0019486 |
| 534 | c56041_g1 | -10.815 | 0.0019486 |
| 535 | c34573_g1 | -10.812 | 0.0019486 |
| 536 | c71560_g1 | -10.810 | 0.0019486 |
| 537 | c41096_g1 | -10.810 | 0.0019486 |
| 538 | c33815_g1 | -10.807 | 0.0019877 |
| 539 | c25365_g1 | -10.807 | 0.0019877 |
| 540 | c53824_g1 | -10.805 | 0.0019877 |
| 541 | c71380_g1 | -10.803 | 0.0019877 |
| 542 | c44257_g1 | -10.801 | 0.002056 |
| 543 | c55492_g4 | -10.790 | 0.00211 |
| 544 | c54547_g1 | -10.790 | 0.00211 |
| 545 | c57715_g4 | -10.789 | 0.00211 |
| 546 | c56474_g1 | -10.789 | 0.00211 |
| 547 | c44265_g1 | -10.788 | 0.0015723 |
| 548 | c45882_g1 | -10.786 | 0.0015723 |
| 549 | c55001_g1 | -10.785 | 0.0015723 |
| 550 | c54844_g2 | -10.785 | 0.00161 |
| 551 | c55059_g1 | -10.783 | 0.00161 |
| 552 | c42893_g1 | -10.783 | 0.00161 |
| 553 | c44418_g1 | -10.782 | 0.00161 |
| 554 | c47490_g1 | -10.782 | 0.00161 |
| 555 | c52055_g1 | -10.782 | 0.00161 |

| | | | |
|---|---|---|---|
| 556 | c31518_g1 | -10.781 | 0.00161 |
| 557 | c71454_g1 | -10.780 | 0.00161 |
| 558 | c58204_g1 | -10.774 | 0.0016614 |
| 559 | c48495_g1 | -10.772 | 0.0016614 |
| 560 | c47345_g1 | -10.771 | 0.0016614 |
| 561 | c55820_g1 | -10.767 | 0.0017026 |
| 562 | c50340_g1 | -10.767 | 0.0017026 |
| 563 | c52485_g2 | -10.765 | 0.0017026 |
| 564 | c43349_g1 | -10.763 | 0.0017026 |
| 565 | c47895_g1 | -10.763 | 0.0017026 |
| 566 | c40062_g1 | -10.761 | 0.0017026 |
| 567 | c56130_g1 | -10.758 | 0.0017654 |
| 568 | c41582_g1 | -10.756 | 0.0017654 |
| 569 | c43718_g1 | -10.751 | 0.0018256 |
| 570 | c48317_g1 | -10.746 | 0.0018256 |
| 571 | c33443_g1 | -10.745 | 0.0018256 |
| 572 | c24109_g1 | -10.739 | 0.0018883 |
| 573 | c44798_g1 | -10.736 | 0.0018883 |
| 574 | c54266_g1 | -10.734 | 0.0019437 |
| 575 | c54698_g1 | -10.734 | 0.0019437 |
| 576 | c52923_g1 | -10.733 | 0.0019437 |
| 577 | c50788_g1 | -10.732 | 0.0019437 |
| 578 | c57286_g1 | -10.731 | 0.0019437 |
| 579 | c57640_g4 | -10.728 | 0.0019875 |
| 580 | c56199_g3 | -10.727 | 0.0019875 |
| 581 | c49739_g1 | -10.726 | 0.0019875 |
| 582 | c54871_g1 | -10.726 | 0.0019875 |
| 583 | c56889_g1 | -10.725 | 0.0019875 |
| 584 | c54728_g1 | -10.723 | 0.0019875 |
| 585 | c47358_g2 | -10.723 | 0.0019875 |
| 586 | c84246_g1 | -10.722 | 0.0019875 |
| 587 | c51579_g1 | -10.716 | 0.0020471 |
| 588 | c56909_g3 | -10.716 | 0.0020471 |
| 589 | c55516_g1 | -10.714 | 0.0020471 |
| 590 | c56454_g1 | -10.707 | 0.00211 |
| 591 | c55624_g1 | -10.707 | 0.00211 |
| 592 | c49571_g2 | -10.704 | 0.00211 |
| 593 | c46416_g1 | -10.701 | 0.0021833 |
| 594 | c54194_g1 | -10.701 | 0.0021833 |
| 595 | c34700_g1 | -10.694 | 0.0022585 |
| 596 | c45456_g1 | -10.693 | 0.0022585 |
| 597 | c57477_g3 | -10.693 | 0.0022585 |
| 598 | c71326_g1 | -10.693 | 0.0022585 |
| 599 | c38436_g1 | -10.687 | 0.00211 |
| 600 | c47533_g1 | -10.687 | 0.0022585 |
| 601 | c53545_g1 | -10.687 | 0.0022585 |
| 602 | c26904_g1 | -10.685 | 0.002343 |
| 603 | c48525_g1 | -10.685 | 0.002343 |
| 604 | c39814_g1 | -10.676 | 0.002428 |
| 605 | c55072_g1 | -10.675 | 0.002343 |
| 606 | c46153_g1 | -10.675 | 0.002428 |
| 607 | c15869_g1 | -10.674 | 0.002428 |
| 608 | c47762_g1 | -10.674 | 0.002428 |
| 609 | c43425_g1 | -10.672 | 0.002428 |
| 610 | c40374_g1 | -10.668 | 0.0025101 |
| 611 | c49752_g1 | -10.667 | 0.0025101 |
| 612 | c50327_g3 | -10.666 | 0.0025101 |
| 613 | c56729_g1 | -10.665 | 0.0025101 |
| 614 | c48034_g1 | -10.664 | 0.0025101 |
| 615 | c31661_g1 | -10.661 | 0.0025101 |
| 616 | c50643_g1 | -10.660 | 0.0025101 |
| 617 | c54408_g2 | -10.652 | 0.0026144 |

| | | | |
|---|---|---|---|
| 618 | c58033_g1 | -10.649 | 0.0027053 |
| 619 | c33324_g1 | -10.648 | 0.0027053 |
| 620 | c55276_g1 | -10.643 | 0.0027053 |
| 621 | c46942_g1 | -10.637 | 0.0028023 |
| 622 | c44028_g1 | -10.632 | 0.0028023 |
| 623 | c56585_g1 | -10.631 | 0.0028925 |
| 624 | c51587_g1 | -10.630 | 0.0028925 |
| 625 | c32004_g1 | -10.630 | 0.0028925 |
| 626 | c57363_g1 | -10.628 | 0.0028925 |
| 627 | c55935_g1 | -10.625 | 0.0030013 |
| 628 | c40796_g1 | -10.622 | 0.0030013 |
| 629 | c52060_g1 | -10.621 | 0.0030013 |
| 630 | c57289_g3 | -10.614 | 0.0031227 |
| 631 | c57984_g2 | -10.604 | 0.0032177 |
| 632 | c51652_g1 | -10.603 | 0.0032177 |
| 633 | c55394_g2 | -10.601 | 0.0032177 |
| 634 | c49669_g1 | -10.594 | 0.0033209 |
| 635 | c55906_g1 | -10.589 | 0.0033209 |
| 636 | c53574_g1 | -10.588 | 0.0034326 |
| 637 | c47443_g1 | -10.587 | 0.0033209 |
| 638 | c52988_g1 | -10.586 | 0.0034326 |
| 639 | c47750_g1 | -10.585 | 0.0034326 |
| 640 | c57784_g1 | -10.583 | 0.0034326 |
| 641 | c71422_g1 | -10.579 | 0.0032177 |
| 642 | c36576_g1 | -10.577 | 0.0035491 |
| 643 | c54053_g5 | -10.571 | 0.0036792 |
| 644 | c51676_g1 | -10.570 | 0.0036792 |
| 645 | c50866_g2 | -10.569 | 0.0036792 |
| 646 | c50070_g1 | -10.568 | 0.0036792 |
| 647 | c52223_g1 | -10.566 | 0.0026292 |
| 648 | c55559_g1 | -10.562 | 0.0026292 |
| 649 | c57509_g1 | -10.562 | 0.0026292 |
| 650 | c53192_g1 | -10.560 | 0.0027264 |
| 651 | c57144_g1 | -10.557 | 0.0027264 |
| 652 | c47367_g1 | -10.557 | 0.0027264 |
| 653 | c58082_g3 | -10.553 | 0.0027264 |
| 654 | c57539_g1 | -10.551 | 0.0028315 |
| 655 | c42527_g1 | -10.546 | 0.0028315 |
| 656 | c46282_g1 | -10.545 | 0.0028315 |
| 657 | c58084_g2 | -10.538 | 0.0029377 |
| 658 | c48978_g1 | -10.532 | 0.0029377 |
| 659 | c55822_g1 | -10.529 | 0.0030601 |
| 660 | c56756_g1 | -10.524 | 0.0031727 |
| 661 | c56763_g2 | -10.522 | 0.0031727 |
| 662 | c39572_g1 | -10.519 | 0.0031727 |
| 663 | c57262_g1 | -10.518 | 0.0031727 |
| 664 | c56719_g1 | -10.517 | 0.0031727 |
| 665 | c44151_g1 | -10.515 | 0.0031727 |
| 666 | c53538_g1 | -10.513 | 0.0032744 |
| 667 | c57187_g5 | -10.511 | 0.0032744 |
| 668 | c58662_g1 | -10.511 | 0.0032744 |
| 669 | c45428_g1 | -10.508 | 0.0032744 |
| 670 | c56728_g1 | -10.507 | 0.0032744 |
| 671 | c26742_g1 | -10.504 | 0.0033969 |
| 672 | c29913_g1 | -10.504 | 0.0032744 |
| 673 | c55400_g2 | -10.504 | 0.0033969 |
| 674 | c29658_g1 | -10.501 | 0.0033969 |
| 675 | c51933_g1 | -10.498 | 0.0033969 |
| 676 | c47222_g2 | -10.497 | 0.0033969 |
| 677 | c54000_g1 | -10.493 | 0.0035126 |
| 678 | c49378_g1 | -10.492 | 0.0035126 |
| 679 | c84189_g1 | -10.491 | 0.0035126 |

| | | | |
|---|---|---|---|
| 680 | c50882_g1 | -10.490 | 0.0035126 |
| 681 | c55183_g1 | -10.489 | 0.0035126 |
| 682 | c59210_g1 | -10.488 | 0.0035126 |
| 683 | c55405_g1 | -10.488 | 0.0035126 |
| 684 | c35218_g1 | -10.483 | 0.0036593 |
| 685 | c54451_g6 | -10.483 | 0.0036593 |
| 686 | c26619_g1 | -10.481 | 0.0036593 |
| 687 | c55812_g2 | -10.475 | 0.0037952 |
| 688 | c52727_g1 | -10.473 | 0.0037952 |
| 689 | c54372_g1 | -10.472 | 0.0037952 |
| 690 | c49094_g1 | -10.471 | 0.0037952 |
| 691 | c52261_g1 | -10.469 | 0.0037952 |
| 692 | c53705_g1 | -10.466 | 0.0039511 |
| 693 | c50646_g1 | -10.465 | 0.0039511 |
| 694 | c52111_g1 | -10.464 | 0.0039511 |
| 695 | c49565_g1 | -10.462 | 0.0039511 |
| 696 | c57023_g2 | -10.462 | 0.0039511 |
| 697 | c52773_g1 | -10.456 | 0.0039511 |
| 698 | c56392_g1 | -10.452 | 0.0041334 |
| 699 | c55322_g1 | -10.450 | 0.0041334 |
| 700 | c55000_g1 | -10.444 | 0.0042905 |
| 701 | c57631_g2 | -10.441 | 0.0042905 |
| 702 | c12847_g1 | -10.441 | 0.0042905 |
| 703 | c50574_g1 | -10.440 | 0.0042905 |
| 704 | c52009_g1 | -10.440 | 0.0042905 |
| 705 | c53327_g1 | -10.440 | 0.0042905 |
| 706 | c43797_g1 | -10.439 | 0.0042905 |
| 707 | c36358_g1 | -10.438 | 0.0042905 |
| 708 | c9297_g1 | -10.433 | 0.0044761 |
| 709 | c53605_g1 | -10.428 | 0.0044761 |
| 710 | c56584_g5 | -10.424 | 0.0044761 |
| 711 | c42941_g2 | -10.424 | 0.004669 |
| 712 | c49409_g3 | -10.420 | 0.004669 |
| 713 | c54297_g1 | -10.417 | 0.004669 |
| 714 | c47608_g1 | -10.413 | 0.0048507 |
| 715 | c45727_g1 | -10.413 | 0.0048507 |
| 716 | c53334_g1 | -10.413 | 0.0048507 |
| 717 | c55081_g1 | -10.412 | 0.0048507 |
| 718 | c52946_g1 | -10.410 | 0.0048507 |
| 719 | c57460_g2 | -10.406 | 0.0048507 |
| 720 | c42741_g1 | -10.406 | 0.0048507 |
| 721 | c34184_g1 | -10.404 | 0.0050608 |
| 722 | c57152_g1 | -10.399 | 0.0050608 |
| 723 | c51680_g1 | -10.398 | 0.0050608 |
| 724 | c57939_g1 | -10.391 | 0.0052289 |
| 725 | c55168_g5 | -10.387 | 0.0052289 |
| 726 | c34387_g1 | -10.386 | 0.0052289 |
| 727 | c31547_g1 | -10.385 | 0.0052289 |
| 728 | c39794_g1 | -10.384 | 0.0053851 |
| 729 | c56944_g1 | -10.381 | 0.0053851 |
| 730 | c38990_g2 | -10.380 | 0.0053851 |
| 731 | c52465_g1 | -10.379 | 0.0053851 |
| 732 | c52188_g2 | -10.379 | 0.0053851 |
| 733 | c49299_g1 | -10.378 | 0.0053851 |
| 734 | c57727_g1 | -10.376 | 0.0053851 |
| 735 | c42943_g1 | -10.376 | 0.0053851 |
| 736 | c54774_g2 | -10.376 | 0.0053851 |
| 737 | c46603_g1 | -10.375 | 0.0053851 |
| 738 | c19475_g1 | -10.373 | 0.0055492 |
| 739 | c52058_g3 | -10.372 | 0.0055492 |
| 740 | c49323_g1 | -10.370 | 0.0055492 |
| 741 | c46209_g1 | -10.368 | 0.0055492 |

| | | | |
|---|---|---|---|
| 742 | c55101_g2 | -10.364 | 0.0055492 |
| 743 | c55389_g3 | -10.364 | 0.0055492 |
| 744 | c52312_g1 | -10.363 | 0.0055492 |
| 745 | c40654_g1 | -10.361 | 0.0057455 |
| 746 | c58057_g8 | -10.359 | 0.0057455 |
| 747 | c56076_g1 | -10.357 | 0.0057455 |
| 748 | c55685_g1 | -10.354 | 0.0057455 |
| 749 | c52176_g1 | -10.353 | 0.0057455 |
| 750 | c49628_g2 | -10.352 | 0.005933 |
| 751 | c46473_g1 | -10.350 | 0.005933 |
| 752 | c57745_g1 | -10.350 | 0.005933 |
| 753 | c39759_g1 | -10.349 | 0.005933 |
| 754 | c40487_g1 | -10.348 | 0.005933 |
| 755 | c122703_g1 | -10.348 | 0.005933 |
| 756 | c56842_g1 | -10.346 | 0.005933 |
| 757 | c55162_g1 | -10.345 | 0.005933 |
| 758 | c58344_g1 | -10.343 | 0.005933 |
| 759 | c51285_g2 | -10.343 | 0.005933 |
| 760 | c52929_g1 | -10.342 | 0.005933 |
| 761 | c55687_g1 | -10.333 | 0.0044761 |
| 762 | c36028_g1 | -10.330 | 0.0044761 |
| 763 | c55027_g1 | -10.329 | 0.0044761 |
| 764 | c30138_g1 | -10.321 | 0.0046842 |
| 765 | c52291_g1 | -10.319 | 0.0046842 |
| 766 | c45909_g1 | -10.316 | 0.0048869 |
| 767 | c55540_g1 | -10.307 | 0.0050608 |
| 768 | c51752_g1 | -10.307 | 0.0050608 |
| 769 | c49176_g1 | -10.307 | 0.0050608 |
| 770 | c54695_g1 | -10.306 | 0.0050608 |
| 771 | c32489_g1 | -10.302 | 0.0050608 |
| 772 | c51285_g3 | -10.301 | 0.0050608 |
| 773 | c18689_g1 | -10.300 | 0.0050608 |
| 774 | c39363_g1 | -10.299 | 0.0050608 |
| 775 | c46987_g1 | -10.298 | 0.0050608 |
| 776 | c48323_g1 | -10.298 | 0.0050608 |
| 777 | c58072_g3 | -10.298 | 0.0050608 |
| 778 | c109723_g1 | -10.297 | 0.0050608 |
| 779 | c43148_g1 | -10.295 | 0.0052607 |
| 780 | c52764_g1 | -10.295 | 0.0052607 |
| 781 | c44191_g1 | -10.294 | 0.0052607 |
| 782 | c53403_g1 | -10.293 | 0.0052607 |
| 783 | c45365_g1 | -10.293 | 0.0052607 |
| 784 | c44312_g1 | -10.293 | 0.0052607 |
| 785 | c49409_g2 | -10.292 | 0.0052607 |
| 786 | c109567_g1 | -10.289 | 0.0052607 |
| 787 | c56094_g1 | -10.284 | 0.0054301 |
| 788 | c37338_g1 | -10.284 | 0.0054301 |
| 789 | c51389_g1 | -10.283 | 0.0054301 |
| 790 | c56235_g1 | -10.283 | 0.0054301 |
| 791 | c49462_g1 | -10.283 | 0.0054301 |
| 792 | c54683_g1 | -10.279 | 0.0054301 |
| 793 | c57701_g1 | -10.278 | 0.0054301 |
| 794 | c42347_g1 | -10.277 | 0.0054301 |
| 795 | c54395_g1 | -10.275 | 0.0054301 |
| 796 | c57826_g1 | -10.275 | 0.0056203 |
| 797 | c38825_g1 | -10.270 | 0.0056203 |
| 798 | c56684_g2 | -10.270 | 0.0056203 |
| 799 | c51148_g1 | -10.269 | 0.0056203 |
| 800 | c39922_g1 | -10.268 | 0.0056203 |
| 801 | c50991_g1 | -10.268 | 0.0056203 |
| 802 | c15074_g1 | -10.266 | 0.0056203 |
| 803 | c57034_g1 | -10.265 | 0.0056203 |

| | | | |
|---|---|---|---|
| 804 | c28798_g1 | -10.262 | 0.0058505 |
| 805 | c46169_g1 | -10.261 | 0.0058505 |
| 806 | c47667_g1 | -10.258 | 0.0058505 |
| 807 | c57501_g1 | -10.257 | 0.0058505 |
| 808 | c49671_g1 | -10.254 | 0.0058505 |
| 809 | c51859_g2 | -10.249 | 0.0060556 |
| 810 | c51490_g1 | -10.247 | 0.0060556 |
| 811 | c58057_g2 | -10.247 | 0.0060556 |
| 812 | c27264_g1 | -10.246 | 0.0060556 |
| 813 | c41739_g1 | -10.245 | 0.0060556 |
| 814 | c55488_g1 | -10.241 | 0.0060556 |
| 815 | c47130_g1 | -10.240 | 0.0063219 |
| 816 | c57259_g2 | -10.239 | 0.0063219 |
| 817 | c15618_g1 | -10.238 | 0.0063219 |
| 818 | c56356_g1 | -10.232 | 0.0063219 |
| 819 | c52305_g1 | -10.232 | 0.0063219 |
| 820 | c23697_g1 | -10.232 | 0.0063219 |
| 821 | c56979_g1 | -10.230 | 0.0063219 |
| 822 | c56948_g1 | -10.226 | 0.0066351 |
| 823 | c53776_g1 | -10.225 | 0.0066351 |
| 824 | c56746_g1 | -10.224 | 0.0066351 |
| 825 | c24327_g2 | -10.221 | 0.0063219 |
| 826 | c48422_g1 | -10.220 | 0.0066351 |
| 827 | c47341_g2 | -10.215 | 0.0069247 |
| 828 | c55121_g1 | -10.214 | 0.0066351 |
| 829 | c46703_g1 | -10.213 | 0.0069247 |
| 830 | c48854_g1 | -10.213 | 0.0069247 |
| 831 | c12732_g1 | -10.212 | 0.0069247 |
| 832 | c53842_g1 | -10.212 | 0.0069247 |
| 833 | c45505_g1 | -10.211 | 0.0069247 |
| 834 | c48531_g1 | -10.211 | 0.0069247 |
| 835 | c45624_g1 | -10.210 | 0.0069247 |
| 836 | c43077_g1 | -10.209 | 0.0069247 |
| 837 | c41928_g1 | -10.207 | 0.0069247 |
| 838 | c55037_g1 | -10.204 | 0.0072155 |
| 839 | c57640_g2 | -10.203 | 0.0072155 |
| 840 | c52987_g1 | -10.200 | 0.0072155 |
| 841 | c57577_g8 | -10.197 | 0.0072155 |
| 842 | c42653_g1 | -10.194 | 0.0072155 |
| 843 | c54844_g1 | -10.192 | 0.0075291 |
| 844 | c47898_g1 | -10.192 | 0.0075291 |
| 845 | c36726_g1 | -10.189 | 0.0075291 |
| 846 | c50637_g1 | -10.189 | 0.0075291 |
| 847 | c23145_g1 | -10.188 | 0.0075291 |
| 848 | c55117_g1 | -10.186 | 0.0075291 |
| 849 | c54799_g1 | -10.186 | 0.0075291 |
| 850 | c39257_g2 | -10.183 | 0.0075291 |
| 851 | c45950_g1 | -10.178 | 0.0078 |
| 852 | c40625_g1 | -10.178 | 0.0078 |
| 853 | c50608_g1 | -10.177 | 0.0078 |
| 854 | c55993_g1 | -10.176 | 0.0078 |
| 855 | c58154_g3 | -10.173 | 0.0078 |
| 856 | c51979_g1 | -10.172 | 0.0078 |
| 857 | c47346_g1 | -10.171 | 0.0078 |
| 858 | c42822_g1 | -10.169 | 0.0081345 |
| 859 | c122430_g1 | -10.168 | 0.0081345 |
| 860 | c50832_g1 | -10.167 | 0.0078 |
| 861 | c45382_g1 | -10.167 | 0.0081345 |
| 862 | c53532_g2 | -10.167 | 0.0081345 |
| 863 | c54908_g1 | -10.165 | 0.0081345 |
| 864 | c57811_g1 | -10.159 | 0.0081345 |
| 865 | c45782_g1 | -10.156 | 0.0085061 |

| | | | |
|---|---|---|---|
| 866 | c50872_g1 | -10.151 | 0.0085061 |
| 867 | c54688_g2 | -10.149 | 0.0085061 |
| 868 | c55042_g1 | -10.149 | 0.0085061 |
| 869 | c43854_g1 | -10.148 | 0.0085061 |
| 870 | c55554_g2 | -10.144 | 0.0088757 |
| 871 | c55743_g1 | -10.143 | 0.0088757 |
| 872 | c57176_g2 | -10.140 | 0.0088757 |
| 873 | c58152_g1 | -10.139 | 0.0088757 |
| 874 | c53172_g1 | -10.138 | 0.0088757 |
| 875 | c49418_g2 | -10.135 | 0.0088757 |
| 876 | c56571_g1 | -10.133 | 0.0088757 |
| 877 | c54036_g1 | -10.132 | 0.0092683 |
| 878 | c54746_g5 | -10.131 | 0.0092683 |
| 879 | c56997_g1 | -10.129 | 0.0092683 |
| 880 | c56879_g1 | -10.128 | 0.0092683 |
| 881 | c55658_g1 | -10.127 | 0.0088757 |
| 882 | c52416_g1 | -10.125 | 0.0092683 |
| 883 | c57599_g1 | -10.124 | 0.0092683 |
| 884 | c36507_g1 | -10.121 | 0.0092683 |
| 885 | c122420_g1 | -10.117 | 0.0096268 |
| 886 | c56205_g4 | -10.115 | 0.0096268 |
| 887 | c56939_g1 | -10.113 | 0.0096268 |
| 888 | c32709_g1 | -10.113 | 0.0096268 |
| 889 | c15554_g1 | -10.112 | 0.0096268 |
| 890 | c56733_g1 | -10.112 | 0.0096268 |
| 891 | c57187_g3 | -10.111 | 0.0096268 |
| 892 | c53558_g1 | -10.111 | 0.0096268 |
| 893 | c55076_g1 | -10.110 | 0.0096268 |
| 894 | c56722_g2 | -10.104 | 0.0071617 |
| 895 | c55677_g1 | -10.103 | 0.0071617 |
| 896 | c57187_g1 | -10.103 | 0.0071617 |
| 897 | c51292_g1 | -10.101 | 0.0071617 |
| 898 | c47850_g1 | -10.101 | 0.0071617 |
| 899 | c39218_g1 | -10.100 | 0.0071617 |
| 900 | c71442_g1 | -10.095 | 0.0071617 |
| 901 | c45175_g1 | -10.094 | 0.0071617 |
| 902 | c55686_g3 | -10.094 | 0.0075211 |
| 903 | c54308_g1 | -10.093 | 0.0075211 |
| 904 | c54371_g1 | -10.084 | 0.0075211 |
| 905 | c46084_g2 | -10.083 | 0.0075211 |
| 906 | c52371_g1 | -10.080 | 0.0078 |
| 907 | c39152_g1 | -10.080 | 0.0078 |
| 908 | c15611_g1 | -10.080 | 0.0078 |
| 909 | c58241_g1 | -10.078 | 0.0078 |
| 910 | c57531_g1 | -10.078 | 0.0078 |
| 911 | c27180_g1 | -10.076 | 0.0078 |
| 912 | c57100_g2 | -10.070 | 0.0078 |
| 913 | c39960_g1 | -10.070 | 0.0078 |
| 914 | c44284_g1 | -10.065 | 0.0081345 |
| 915 | c54828_g1 | -10.065 | 0.0078 |
| 916 | c50787_g1 | -10.063 | 0.0081345 |
| 917 | c51794_g2 | -10.061 | 0.0081345 |
| 918 | c44548_g1 | -10.061 | 0.0081345 |
| 919 | c23397_g1 | -10.056 | 0.0081345 |
| 920 | c55613_g6 | -10.055 | 0.0081345 |
| 921 | c41234_g1 | -10.050 | 0.0085061 |
| 922 | c56676_g1 | -10.049 | 0.0085061 |
| 923 | c27163_g1 | -10.048 | 0.0085061 |
| 924 | c55810_g1 | -10.048 | 0.0085061 |
| 925 | c51164_g1 | -10.047 | 0.0085061 |
| 926 | c56560_g1 | -10.046 | 0.0085061 |
| 927 | c41890_g1 | -10.042 | 0.0085061 |

| | | | |
|---|---|---|---|
| 928 | c36021_g1 | -10.042 | 0.0088757 |
| 929 | c56008_g1 | -10.039 | 0.0088757 |
| 930 | c53236_g1 | -10.039 | 0.0088757 |
| 931 | c55613_g3 | -10.039 | 0.0088757 |
| 932 | c54242_g1 | -10.038 | 0.0088757 |
| 933 | c55614_g1 | -10.035 | 0.0088757 |
| 934 | c40857_g1 | -10.034 | 0.0088757 |
| 935 | c24730_g1 | -10.031 | 0.0088757 |
| 936 | c53163_g1 | -10.030 | 0.0088757 |
| 937 | c34548_g1 | -10.029 | 0.0088757 |
| 938 | c41146_g1 | -10.026 | 0.0092999 |
| 939 | c49709_g1 | -10.025 | 0.0092999 |
| 940 | c52327_g1 | -10.024 | 0.0092999 |
| 941 | c56526_g1 | -10.022 | 0.0092999 |
| 942 | c57024_g2 | -10.019 | 0.0092999 |
| 943 | c58072_g6 | -10.017 | 0.0092999 |
| 944 | c54300_g2 | -10.015 | 0.0092999 |
| 945 | c57740_g4 | -10.009 | 0.0096976 |
| 946 | c43034_g1 | -10.009 | 0.0096976 |
| 947 | c32002_g1 | -10.009 | 0.0096976 |
| 948 | c24475_g1 | -10.006 | 0.0096976 |
| 949 | c31400_g1 | -10.005 | 0.0096976 |
| 950 | c57338_g4 | -10.005 | 0.0096976 |
| 951 | c49231_g1 | -10.004 | 0.0096976 |
| 952 | c45608_g1 | -9.998 | 0.0102336 |
| 953 | c53816_g3 | -9.996 | 0.0102336 |
| 954 | c50901_g1 | -9.995 | 0.0102336 |
| 955 | c54736_g1 | -9.990 | 0.0102336 |
| 956 | c53769_g1 | -9.990 | 0.0102336 |
| 957 | c57969_g2 | -9.989 | 0.0102336 |
| 958 | c54905_g1 | -9.979 | 0.0107657 |
| 959 | c44672_g1 | -9.978 | 0.0107657 |
| 960 | c54801_g1 | -9.977 | 0.0107657 |
| 961 | c54953_g1 | -9.975 | 0.0107657 |
| 962 | c54222_g1 | -9.975 | 0.0107657 |
| 963 | c50560_g1 | -9.975 | 0.0107657 |
| 964 | c52157_g1 | -9.975 | 0.0107657 |
| 965 | c54808_g2 | -9.971 | 0.0113302 |
| 966 | c56238_g1 | -9.968 | 0.0113302 |
| 967 | c40865_g1 | -9.967 | 0.0113302 |
| 968 | c56069_g1 | -9.965 | 0.0113302 |
| 969 | c53355_g1 | -9.965 | 0.0113302 |
| 970 | c56982_g1 | -9.964 | 0.0113302 |
| 971 | c29766_g1 | -9.960 | 0.0118789 |
| 972 | c55632_g1 | -9.958 | 0.0118789 |
| 973 | c49740_g1 | -9.958 | 0.0118789 |
| 974 | c26797_g1 | -9.957 | 0.0118789 |
| 975 | c50640_g1 | -9.947 | 0.0118789 |
| 976 | c52108_g1 | -9.946 | 0.0118789 |
| 977 | c32403_g1 | -9.943 | 0.0124088 |
| 978 | c15658_g1 | -9.943 | 0.0124088 |
| 979 | c45373_g1 | -9.942 | 0.0124088 |
| 980 | c43369_g1 | -9.940 | 0.0124088 |
| 981 | c45565_g1 | -9.938 | 0.0124088 |
| 982 | c47134_g1 | -9.937 | 0.0124088 |
| 983 | c47237_g1 | -9.936 | 0.0124088 |
| 984 | c50775_g1 | -9.936 | 0.0124088 |
| 985 | c57014_g2 | -9.936 | 0.0124088 |
| 986 | c71411_g1 | -9.934 | 0.0124088 |
| 987 | c51178_g1 | -9.932 | 0.0124088 |
| 988 | c53368_g2 | -9.932 | 0.0124088 |
| 989 | c54954_g1 | -9.928 | 0.0129074 |

| | | | |
|---|---|---|---|
| 990 | c53038_g1 | -9.928 | 0.0129074 |
| 991 | c56234_g1 | -9.926 | 0.0129074 |
| 992 | c57238_g1 | -9.923 | 0.0129074 |
| 993 | c36278_g1 | -9.920 | 0.0129074 |
| 994 | c56636_g1 | -9.920 | 0.0129074 |
| 995 | c57935_g3 | -9.916 | 0.0134691 |
| 996 | c54902_g1 | -9.915 | 0.0129074 |
| 997 | c57822_g1 | -9.915 | 0.0134691 |
| 998 | c34913_g1 | -9.910 | 0.0134691 |
| 999 | c97179_g1 | -9.908 | 0.0134691 |
| 1000 | c109627_g1 | -9.908 | 0.0134691 |
| 1001 | c55512_g3 | -9.905 | 0.0134691 |
| 1002 | c56768_g6 | -9.905 | 0.0134691 |
| 1003 | c57411_g1 | -9.903 | 0.0134691 |
| 1004 | c36479_g1 | -9.903 | 0.0134691 |
| 1005 | c49256_g3 | -9.901 | 0.0141427 |
| 1006 | c43198_g1 | -9.900 | 0.0141427 |
| 1007 | c39193_g1 | -9.899 | 0.0141427 |
| 1008 | c46072_g1 | -9.897 | 0.0141427 |
| 1009 | c51136_g1 | -9.896 | 0.0141427 |
| 1010 | c53620_g1 | -9.891 | 0.0141427 |
| 1011 | c39648_g1 | -9.889 | 0.0141427 |
| 1012 | c46841_g1 | -9.889 | 0.0141427 |
| 1013 | c52075_g1 | -9.885 | 0.0147433 |
| 1014 | c35155_g1 | -9.884 | 0.0147433 |
| 1015 | c42270_g1 | -9.884 | 0.0147433 |
| 1016 | c56000_g1 | -9.879 | 0.0147433 |
| 1017 | c56584_g4 | -9.878 | 0.0147433 |
| 1018 | c57758_g1 | -9.874 | 0.0147433 |
| 1019 | c10749_g1 | -9.874 | 0.0147433 |
| 1020 | c54014_g2 | -9.874 | 0.0147433 |
| 1021 | c71420_g1 | -9.874 | 0.0147433 |
| 1022 | c57589_g1 | -9.873 | 0.0147433 |
| 1023 | c54157_g1 | -9.872 | 0.0155187 |
| 1024 | c71344_g1 | -9.870 | 0.0155187 |
| 1025 | c48549_g1 | -9.867 | 0.0155187 |
| 1026 | c31286_g1 | -9.860 | 0.0109679 |
| 1027 | c56307_g2 | -9.860 | 0.0109679 |
| 1028 | c54309_g1 | -9.859 | 0.0109679 |
| 1029 | c49902_g1 | -9.858 | 0.0109679 |
| 1030 | c55151_g1 | -9.857 | 0.0115466 |
| 1031 | c39500_g1 | -9.857 | 0.0115466 |
| 1032 | c56071_g1 | -9.857 | 0.0115466 |
| 1033 | c52695_g1 | -9.856 | 0.0115466 |
| 1034 | c49932_g2 | -9.856 | 0.0109679 |
| 1035 | c54951_g1 | -9.851 | 0.0115466 |
| 1036 | c58347_g1 | -9.847 | 0.0115466 |
| 1037 | c50861_g1 | -9.846 | 0.0115466 |
| 1038 | c56129_g1 | -9.845 | 0.0115466 |
| 1039 | c25307_g1 | -9.842 | 0.0115466 |
| 1040 | c49121_g2 | -9.840 | 0.0115466 |
| 1041 | c53384_g1 | -9.840 | 0.0121831 |
| 1042 | c53426_g5 | -9.839 | 0.0121831 |
| 1043 | c55223_g1 | -9.838 | 0.0121831 |
| 1044 | c55496_g1 | -9.834 | 0.0121831 |
| 1045 | c53306_g2 | -9.832 | 0.0121831 |
| 1046 | c54643_g1 | -9.832 | 0.0121831 |
| 1047 | c56063_g4 | -9.832 | 0.0121831 |
| 1048 | c30098_g1 | -9.827 | 0.0121831 |
| 1049 | c50591_g1 | -9.826 | 0.0126618 |
| 1050 | c46475_g1 | -9.826 | 0.0126618 |
| 1051 | c22439_g1 | -9.826 | 0.0126618 |

| | | | |
|---|---|---|---|
| 1052 | c49278_g1 | -9.823 | 0.0126618 |
| 1053 | c58082_g4 | -9.823 | 0.0126618 |
| 1054 | c57699_g1 | -9.823 | 0.0126618 |
| 1055 | c51421_g1 | -9.821 | 0.0126618 |
| 1056 | c57573_g2 | -9.821 | 0.0126618 |
| 1057 | c32766_g1 | -9.821 | 0.0126618 |
| 1058 | c51476_g1 | -9.820 | 0.0126618 |
| 1059 | c31551_g2 | -9.819 | 0.0126618 |
| 1060 | c49420_g1 | -9.818 | 0.0126618 |
| 1061 | c47492_g1 | -9.818 | 0.0126618 |
| 1062 | c57128_g3 | -9.817 | 0.0126618 |
| 1063 | c49608_g1 | -9.816 | 0.0126618 |
| 1064 | c55678_g1 | -9.816 | 0.0126618 |
| 1065 | c42178_g1 | -9.816 | 0.0126618 |
| 1066 | c42395_g1 | -9.813 | 0.0126618 |
| 1067 | c54276_g1 | -9.811 | 0.0126618 |
| 1068 | c52666_g1 | -9.811 | 0.013301 |
| 1069 | c48575_g2 | -9.810 | 0.013301 |
| 1070 | c52349_g1 | -9.809 | 0.0126618 |
| 1071 | c84298_g1 | -9.808 | 0.013301 |
| 1072 | c57615_g2 | -9.808 | 0.013301 |
| 1073 | c24295_g1 | -9.807 | 0.013301 |
| 1074 | c45563_g1 | -9.806 | 0.013301 |
| 1075 | c55735_g1 | -9.806 | 0.013301 |
| 1076 | c55077_g1 | -9.806 | 0.013301 |
| 1077 | c49160_g1 | -9.805 | 0.013301 |
| 1078 | c40045_g1 | -9.804 | 0.013301 |
| 1079 | c57922_g1 | -9.803 | 0.013301 |
| 1080 | c57725_g1 | -9.801 | 0.013301 |
| 1081 | c54324_g2 | -9.801 | 0.013301 |
| 1082 | c57544_g1 | -9.799 | 0.013301 |
| 1083 | c36360_g1 | -9.794 | 0.0140039 |
| 1084 | c40071_g1 | -9.793 | 0.0140039 |
| 1085 | c48948_g1 | -9.790 | 0.0140039 |
| 1086 | c53635_g1 | -9.790 | 0.0140039 |
| 1087 | c42438_g1 | -9.786 | 0.0140039 |
| 1088 | c34502_g1 | -9.785 | 0.0140039 |
| 1089 | c24986_g1 | -9.781 | 0.0140039 |
| 1090 | c55911_g2 | -9.781 | 0.0140039 |
| 1091 | c57114_g1 | -9.781 | 0.0140039 |
| 1092 | c56922_g1 | -9.777 | 0.0146959 |
| 1093 | c55345_g1 | -9.776 | 0.0146959 |
| 1094 | c8189_g1 | -9.775 | 0.0146959 |
| 1095 | c57149_g1 | -9.775 | 0.0146959 |
| 1096 | c36529_g1 | -9.773 | 0.0146959 |
| 1097 | c57276_g5 | -9.772 | 0.0146959 |
| 1098 | c34861_g1 | -9.772 | 0.0146959 |
| 1099 | c29347_g1 | -9.771 | 0.0146959 |
| 1100 | c56324_g1 | -9.769 | 0.0146959 |
| 1101 | c33415_g1 | -9.769 | 0.0146959 |
| 1102 | c15576_g1 | -9.766 | 0.0146959 |
| 1103 | c39441_g1 | -9.765 | 0.0146959 |
| 1104 | c54682_g5 | -9.764 | 0.0146959 |
| 1105 | c55907_g1 | -9.762 | 0.0154454 |
| 1106 | c37516_g1 | -9.761 | 0.0154454 |
| 1107 | c46405_g1 | -9.761 | 0.0154454 |
| 1108 | c50002_g1 | -9.760 | 0.0154454 |
| 1109 | c40821_g1 | -9.760 | 0.0154454 |
| 1110 | c47692_g1 | -9.758 | 0.0146959 |
| 1111 | c57054_g1 | -9.755 | 0.0154454 |
| 1112 | c55049_g1 | -9.754 | 0.0154454 |
| 1113 | c55213_g1 | -9.752 | 0.0154454 |

| | | | |
|---|---|---|---|
| 1114 | c51484_g1 | -9.752 | 0.0154454 |
| 1115 | c58230_g1 | -9.752 | 0.0154454 |
| 1116 | c55256_g1 | -9.752 | 0.0154454 |
| 1117 | c47601_g1 | -9.745 | 0.0154454 |
| 1118 | c52776_g1 | -9.742 | 0.0162934 |
| 1119 | c57828_g1 | -9.742 | 0.0162934 |
| 1120 | c34968_g1 | -9.741 | 0.0162934 |
| 1121 | c43974_g1 | -9.741 | 0.0162934 |
| 1122 | c54951_g2 | -9.740 | 0.0162934 |
| 1123 | c56473_g1 | -9.739 | 0.0162934 |
| 1124 | c43307_g1 | -9.738 | 0.0162934 |
| 1125 | c53340_g1 | -9.738 | 0.0162934 |
| 1126 | c52826_g1 | -9.735 | 0.0162934 |
| 1127 | c96838_g1 | -9.733 | 0.0162934 |
| 1128 | c122472_g1 | -9.733 | 0.0162934 |
| 1129 | c32453_g1 | -9.732 | 0.0162934 |
| 1130 | c52261_g4 | -9.731 | 0.0162934 |
| 1131 | c57306_g2 | -9.727 | 0.0172305 |
| 1132 | c55170_g12 | -9.726 | 0.0172305 |
| 1133 | c47712_g1 | -9.726 | 0.0172305 |
| 1134 | c57655_g3 | -9.725 | 0.0172305 |
| 1135 | c53617_g1 | -9.723 | 0.0172305 |
| 1136 | c57571_g4 | -9.718 | 0.0172305 |
| 1137 | c45092_g1 | -9.717 | 0.0172305 |
| 1138 | c56198_g1 | -9.716 | 0.0172305 |
| 1139 | c53555_g1 | -9.715 | 0.0172305 |
| 1140 | c55150_g1 | -9.712 | 0.0181507 |
| 1141 | c55990_g3 | -9.712 | 0.0162934 |
| 1142 | c57276_g1 | -9.712 | 0.0181507 |
| 1143 | c44810_g1 | -9.707 | 0.0181507 |
| 1144 | c36265_g1 | -9.705 | 0.0181507 |
| 1145 | c26471_g1 | -9.705 | 0.0181507 |
| 1146 | c52193_g1 | -9.703 | 0.0181507 |
| 1147 | c53020_g1 | -9.703 | 0.0181507 |
| 1148 | c46566_g1 | -9.703 | 0.0181507 |
| 1149 | c57638_g1 | -9.702 | 0.0181507 |
| 1150 | c46174_g1 | -9.701 | 0.0181507 |
| 1151 | c55764_g1 | -9.700 | 0.0181507 |
| 1152 | c54176_g1 | -9.700 | 0.0181507 |
| 1153 | c58030_g1 | -9.698 | 0.0181507 |
| 1154 | c53377_g1 | -9.698 | 0.0191281 |
| 1155 | c39309_g1 | -9.696 | 0.0191281 |
| 1156 | c53094_g7 | -9.694 | 0.0191281 |
| 1157 | c50542_g1 | -9.692 | 0.0191281 |
| 1158 | c52101_g1 | -9.691 | 0.0191281 |
| 1159 | c47196_g1 | -9.689 | 0.0191281 |
| 1160 | c53375_g1 | -9.688 | 0.0191281 |
| 1161 | c54254_g1 | -9.687 | 0.0191281 |
| 1162 | c51736_g1 | -9.686 | 0.0191281 |
| 1163 | c56722_g3 | -9.685 | 0.0191281 |
| 1164 | c51193_g1 | -9.685 | 0.0191281 |
| 1165 | c38269_g1 | -9.684 | 0.0191281 |
| 1166 | c50904_g1 | -9.684 | 0.0191281 |
| 1167 | c52313_g1 | -9.682 | 0.0191281 |
| 1168 | c56254_g1 | -9.681 | 0.0202282 |
| 1169 | c30304_g1 | -9.680 | 0.0202282 |
| 1170 | c30953_g1 | -9.680 | 0.0202282 |
| 1171 | c53527_g1 | -9.678 | 0.0202282 |
| 1172 | c55809_g5 | -9.676 | 0.0202282 |
| 1173 | c53808_g2 | -9.676 | 0.0202282 |
| 1174 | c50138_g1 | -9.675 | 0.0202282 |
| 1175 | c58181_g1 | -9.671 | 0.0202282 |

| | | | |
|---|---|---|---|
| 1176 | c52403_g1 | -9.667 | 0.0202282 |
| 1177 | c52467_g1 | -9.666 | 0.0202282 |
| 1178 | c55912_g1 | -9.665 | 0.0202282 |
| 1179 | c56692_g1 | -9.663 | 0.0213894 |
| 1180 | c57208_g1 | -9.663 | 0.0213894 |
| 1181 | c47378_g1 | -9.663 | 0.0213894 |
| 1182 | c36438_g1 | -9.662 | 0.0202282 |
| 1183 | c57778_g4 | -9.659 | 0.0213894 |
| 1184 | c30784_g2 | -9.658 | 0.0213894 |
| 1185 | c40016_g1 | -9.656 | 0.0213894 |
| 1186 | c49206_g1 | -9.650 | 0.0213894 |
| 1187 | c39135_g1 | -9.648 | 0.0213894 |
| 1188 | c50501_g1 | -9.645 | 0.0225845 |
| 1189 | c84745_g1 | -9.644 | 0.0213894 |
| 1190 | c47662_g1 | -9.642 | 0.0225845 |
| 1191 | c23731_g1 | -9.642 | 0.0225845 |
| 1192 | c57257_g1 | -9.641 | 0.0225845 |
| 1193 | c52737_g1 | -9.639 | 0.0225845 |
| 1194 | c54488_g1 | -9.631 | 0.0225845 |
| 1195 | c54611_g1 | -9.631 | 0.0225845 |
| 1196 | c45117_g1 | -9.631 | 0.0225845 |
| 1197 | c52880_g2 | -9.630 | 0.0225845 |
| 1198 | c52241_g1 | -9.629 | 0.0225845 |
| 1199 | c109657_g1 | -9.626 | 0.0239111 |
| 1200 | c42425_g1 | -9.625 | 0.0239111 |
| 1201 | c42588_g1 | -9.618 | 0.0239111 |
| 1202 | c27304_g1 | -9.617 | 0.0239111 |
| 1203 | c54978_g1 | -9.613 | 0.0239111 |
| 1204 | c48768_g1 | -9.612 | 0.0225845 |
| 1205 | c54765_g3 | -9.611 | 0.0239111 |
| 1206 | c55659_g1 | -9.610 | 0.0252971 |
| 1207 | c42232_g1 | -9.607 | 0.0252971 |
| 1208 | c15822_g1 | -9.605 | 0.0172305 |
| 1209 | c57559_g1 | -9.604 | 0.0172305 |
| 1210 | c44831_g2 | -9.604 | 0.0162934 |
| 1211 | c46562_g1 | -9.604 | 0.0172305 |
| 1212 | c54339_g3 | -9.601 | 0.0172305 |
| 1213 | c51358_g4 | -9.598 | 0.0172305 |
| 1214 | c53117_g1 | -9.597 | 0.0172305 |
| 1215 | c50650_g1 | -9.597 | 0.0172305 |
| 1216 | c54247_g1 | -9.593 | 0.0172305 |
| 1217 | c24805_g1 | -9.593 | 0.0181507 |
| 1218 | c43052_g1 | -9.593 | 0.0181507 |
| 1219 | c57993_g1 | -9.593 | 0.0181507 |
| 1220 | c53439_g1 | -9.592 | 0.0181507 |
| 1221 | c32858_g1 | -9.591 | 0.0181507 |
| 1222 | c49559_g1 | -9.590 | 0.0181507 |
| 1223 | c33261_g1 | -9.587 | 0.0181507 |
| 1224 | c55023_g1 | -9.587 | 0.0181507 |
| 1225 | c13933_g1 | -9.583 | 0.0181507 |
| 1226 | c54013_g1 | -9.582 | 0.0181507 |
| 1227 | c43343_g2 | -9.581 | 0.0181507 |
| 1228 | c51845_g2 | -9.581 | 0.0181507 |
| 1229 | c55990_g1 | -9.580 | 0.0181507 |
| 1230 | c52069_g1 | -9.580 | 0.0181507 |
| 1231 | c52030_g1 | -9.580 | 0.0181507 |
| 1232 | c34702_g1 | -9.572 | 0.0192477 |
| 1233 | c56510_g1 | -9.570 | 0.0192477 |
| 1234 | c33638_g1 | -9.570 | 0.0192477 |
| 1235 | c53862_g1 | -9.569 | 0.0192477 |
| 1236 | c51316_g1 | -9.565 | 0.0192477 |
| 1237 | c53479_g1 | -9.563 | 0.0192477 |

| | | | |
|---|---|---|---|
| 1238 | c46120_g1 | -9.559 | 0.0192477 |
| 1239 | c45690_g1 | -9.550 | 0.0204116 |
| 1240 | c55094_g2 | -9.549 | 0.0204116 |
| 1241 | c55939_g3 | -9.549 | 0.0204116 |
| 1242 | c55038_g1 | -9.545 | 0.0204116 |
| 1243 | c52815_g1 | -9.544 | 0.0204116 |
| 1244 | c50029_g1 | -9.544 | 0.0204116 |
| 1245 | c46894_g1 | -9.541 | 0.0204116 |
| 1246 | c53476_g1 | -9.540 | 0.0204116 |
| 1247 | c26573_g1 | -9.539 | 0.0204116 |
| 1248 | c34714_g1 | -9.538 | 0.0204116 |
| 1249 | c19430_g1 | -9.537 | 0.0216921 |
| 1250 | c42021_g1 | -9.532 | 0.0216921 |
| 1251 | c38351_g1 | -9.532 | 0.0216921 |
| 1252 | c53337_g1 | -9.532 | 0.0216921 |
| 1253 | c50986_g2 | -9.531 | 0.0216921 |
| 1254 | c55474_g1 | -9.530 | 0.0216921 |
| 1255 | c57420_g1 | -9.528 | 0.0216921 |
| 1256 | c56727_g1 | -9.526 | 0.0216921 |
| 1257 | c122580_g1 | -9.525 | 0.0216921 |
| 1258 | c57042_g1 | -9.525 | 0.0204116 |
| 1259 | c10879_g1 | -9.519 | 0.0216921 |
| 1260 | c56282_g1 | -9.519 | 0.0216921 |
| 1261 | c48536_g1 | -9.518 | 0.0230376 |
| 1262 | c50915_g1 | -9.513 | 0.0230376 |
| 1263 | c56989_g1 | -9.508 | 0.0230376 |
| 1264 | c53726_g1 | -9.506 | 0.0230376 |
| 1265 | c56772_g1 | -9.505 | 0.0230376 |
| 1266 | c55348_g1 | -9.505 | 0.0230376 |
| 1267 | c26695_g1 | -9.503 | 0.0230376 |
| 1268 | c53478_g1 | -9.499 | 0.0244192 |
| 1269 | c33554_g2 | -9.498 | 0.0244192 |
| 1270 | c57244_g1 | -9.498 | 0.0244192 |
| 1271 | c42490_g1 | -9.496 | 0.0244192 |
| 1272 | c49452_g1 | -9.494 | 0.0230376 |
| 1273 | c49458_g1 | -9.494 | 0.0244192 |
| 1274 | c54341_g1 | -9.494 | 0.0244192 |
| 1275 | c44767_g1 | -9.493 | 0.0244192 |
| 1276 | c40268_g1 | -9.493 | 0.0244192 |
| 1277 | c55715_g1 | -9.492 | 0.0244192 |
| 1278 | c52105_g1 | -9.491 | 0.0244192 |
| 1279 | c55514_g1 | -9.490 | 0.0244192 |
| 1280 | c57045_g1 | -9.487 | 0.0244192 |
| 1281 | c50007_g1 | -9.487 | 0.0244192 |
| 1282 | c49617_g1 | -9.486 | 0.0244192 |
| 1283 | c45487_g1 | -9.484 | 0.0244192 |
| 1284 | c51359_g1 | -9.481 | 0.0260766 |
| 1285 | c53019_g1 | -9.480 | 0.0260766 |
| 1286 | c57236_g1 | -9.478 | 0.0244192 |
| 1287 | c30880_g1 | -9.478 | 0.0260766 |
| 1288 | c52057_g1 | -9.478 | 0.0260766 |
| 1289 | c47485_g2 | -9.477 | 0.0260766 |
| 1290 | c57592_g3 | -9.474 | 0.0260766 |
| 1291 | c56458_g1 | -9.472 | 0.0260766 |
| 1292 | c55628_g2 | -9.472 | 0.0260766 |
| 1293 | c56341_g1 | -9.471 | 0.0260766 |
| 1294 | c53204_g1 | -9.470 | 0.0260766 |
| 1295 | c24345_g1 | -9.469 | 0.0260766 |
| 1296 | c39162_g1 | -9.466 | 0.0260766 |
| 1297 | c54326_g1 | -9.466 | 0.0260766 |
| 1298 | c50903_g1 | -9.458 | 0.0276647 |
| 1299 | c10085_g1 | -9.457 | 0.0276647 |

| | | | |
|---|---|---|---|
| 1300 | c42539_g2 | -9.454 | 0.0276647 |
| 1301 | c57796_g2 | -9.450 | 0.0276647 |
| 1302 | c51282_g3 | -9.450 | 0.0276647 |
| 1303 | c46442_g1 | -9.450 | 0.0276647 |
| 1304 | c34124_g1 | -9.446 | 0.0276647 |
| 1305 | c54050_g1 | -9.446 | 0.0244192 |
| 1306 | c56461_g1 | -9.445 | 0.0276647 |
| 1307 | c48586_g1 | -9.445 | 0.0276647 |
| 1308 | c56931_g1 | -9.442 | 0.0276647 |
| 1309 | c31076_g1 | -9.441 | 0.0276647 |
| 1310 | c44533_g1 | -9.441 | 0.0276647 |
| 1311 | c57181_g1 | -9.438 | 0.0292992 |
| 1312 | c51159_g1 | -9.435 | 0.0292992 |
| 1313 | c31842_g1 | -9.434 | 0.0292992 |
| 1314 | c57800_g1 | -9.429 | 0.0292992 |
| 1315 | c51458_g1 | -9.428 | 0.0292992 |
| 1316 | c52428_g1 | -9.427 | 0.0292992 |
| 1317 | c47521_g1 | -9.426 | 0.0292992 |
| 1318 | c54872_g2 | -9.424 | 0.0292992 |
| 1319 | c55066_g1 | -9.424 | 0.0292992 |
| 1320 | c58111_g1 | -9.423 | 0.0292992 |
| 1321 | c57565_g3 | -9.422 | 0.0292992 |
| 1322 | c7777_g1 | -9.422 | 0.0292992 |
| 1323 | c23688_g1 | -9.422 | 0.0292992 |
| 1324 | c46019_g1 | -9.421 | 0.0292992 |
| 1325 | c55766_g1 | -9.420 | 0.0292992 |
| 1326 | c53942_g1 | -9.418 | 0.0311564 |
| 1327 | c54701_g1 | -9.416 | 0.0292992 |
| 1328 | c54616_g1 | -9.409 | 0.0311564 |
| 1329 | c57094_g2 | -9.408 | 0.0311564 |
| 1330 | c55989_g2 | -9.407 | 0.0311564 |
| 1331 | c57233_g1 | -9.403 | 0.0311564 |
| 1332 | c57039_g2 | -9.402 | 0.0311564 |
| 1333 | c10746_g1 | -9.401 | 0.0311564 |
| 1334 | c56106_g3 | -9.401 | 0.0311564 |
| 1335 | c45294_g1 | -9.400 | 0.0311564 |
| 1336 | c57887_g1 | -9.399 | 0.0330762 |
| 1337 | c54437_g1 | -9.399 | 0.0330762 |
| 1338 | c34719_g1 | -9.397 | 0.0330762 |
| 1339 | c57624_g4 | -9.395 | 0.0330762 |
| 1340 | c45412_g1 | -9.394 | 0.0330762 |
| 1341 | c52378_g1 | -9.391 | 0.0330762 |
| 1342 | c23207_g1 | -9.390 | 0.0330762 |
| 1343 | c56445_g1 | -9.390 | 0.0330762 |
| 1344 | c44560_g1 | -9.388 | 0.0330762 |
| 1345 | c33984_g1 | -9.385 | 0.0330762 |
| 1346 | c57083_g1 | -9.385 | 0.0330762 |
| 1347 | c56031_g1 | -9.384 | 0.0330762 |
| 1348 | c54493_g2 | -9.384 | 0.0330762 |
| 1349 | c57525_g2 | -9.383 | 0.0330762 |
| 1350 | c52884_g2 | -9.382 | 0.0330762 |
| 1351 | c49777_g1 | -9.382 | 0.0330762 |
| 1352 | c52993_g1 | -9.381 | 0.0330762 |
| 1353 | c55968_g1 | -9.379 | 0.0351876 |
| 1354 | c46969_g1 | -9.375 | 0.0351876 |
| 1355 | c54870_g1 | -9.368 | 0.0351876 |
| 1356 | c22919_g1 | -9.367 | 0.0351876 |
| 1357 | c39254_g1 | -9.367 | 0.0351876 |
| 1358 | c54049_g1 | -9.366 | 0.0351876 |
| 1359 | c42321_g1 | -9.364 | 0.0351876 |
| 1360 | c46027_g1 | -9.360 | 0.0351876 |
| 1361 | c43968_g1 | -9.359 | 0.0351876 |

| | | | |
|---|---|---|---|
| 1362 | c25322_g1 | -9.359 | 0.0351876 |
| 1363 | c57338_g3 | -9.357 | 0.0375986 |
| 1364 | c27232_g1 | -9.357 | 0.0375986 |
| 1365 | c55160_g1 | -9.356 | 0.0375986 |
| 1366 | c50571_g1 | -9.354 | 0.0375986 |
| 1367 | c53541_g1 | -9.353 | 0.0375986 |
| 1368 | c42623_g1 | -9.351 | 0.0351876 |
| 1369 | c47541_g1 | -9.350 | 0.0375986 |
| 1370 | c57313_g1 | -9.349 | 0.0375986 |
| 1371 | c48858_g1 | -9.349 | 0.0375986 |
| 1372 | c57466_g2 | -9.347 | 0.0375986 |
| 1373 | c57530_g1 | -9.347 | 0.0375986 |
| 1374 | c56322_g1 | -9.342 | 0.0375986 |
| 1375 | c84178_g1 | -9.337 | 0.0375986 |
| 1376 | c57841_g1 | -9.333 | 0.0404152 |
| 1377 | c19925_g1 | -9.331 | 0.0269849 |
| 1378 | c58103_g2 | -9.329 | 0.0269849 |
| 1379 | c55043_g1 | -9.329 | 0.0269849 |
| 1380 | c39594_g1 | -9.329 | 0.0269849 |
| 1381 | c98332_g1 | -9.329 | 0.0269849 |
| 1382 | c35450_g1 | -9.327 | 0.0269849 |
| 1383 | c56080_g1 | -9.325 | 0.0269849 |
| 1384 | c34576_g1 | -9.320 | 0.0269849 |
| 1385 | c55673_g1 | -9.320 | 0.0269849 |
| 1386 | c54525_g1 | -9.319 | 0.0269849 |
| 1387 | c40934_g1 | -9.319 | 0.0237176 |
| 1388 | c36605_g1 | -9.317 | 0.0269849 |
| 1389 | c27926_g1 | -9.316 | 0.0269849 |
| 1390 | c53292_g1 | -9.315 | 0.0269849 |
| 1391 | c55699_g1 | -9.314 | 0.028834 |
| 1392 | c51209_g1 | -9.313 | 0.028834 |
| 1393 | c55000_g3 | -9.305 | 0.028834 |
| 1394 | c52130_g1 | -9.304 | 0.028834 |
| 1395 | c54926_g1 | -9.303 | 0.028834 |
| 1396 | c26857_g1 | -9.302 | 0.028834 |
| 1397 | c52884_g1 | -9.302 | 0.028834 |
| 1398 | c56386_g1 | -9.300 | 0.028834 |
| 1399 | c56382_g1 | -9.298 | 0.028834 |
| 1400 | c42456_g1 | -9.298 | 0.028834 |
| 1401 | c55586_g1 | -9.296 | 0.028834 |
| 1402 | c96939_g1 | -9.294 | 0.0252971 |
| 1403 | c57957_g1 | -9.293 | 0.028834 |
| 1404 | c51118_g1 | -9.292 | 0.028834 |
| 1405 | c47829_g2 | -9.290 | 0.0307605 |
| 1406 | c53806_g1 | -9.289 | 0.0307605 |
| 1407 | c57561_g1 | -9.289 | 0.028834 |
| 1408 | c38329_g1 | -9.288 | 0.0307605 |
| 1409 | c56272_g1 | -9.287 | 0.0307605 |
| 1410 | c55343_g1 | -9.287 | 0.0307605 |
| 1411 | c10553_g1 | -9.285 | 0.0307605 |
| 1412 | c53176_g1 | -9.279 | 0.0307605 |
| 1413 | c57126_g1 | -9.278 | 0.0307605 |
| 1414 | c48817_g1 | -9.278 | 0.0307605 |
| 1415 | c57392_g1 | -9.276 | 0.0307605 |
| 1416 | c40234_g1 | -9.275 | 0.0307605 |
| 1417 | c56299_g1 | -9.274 | 0.0269849 |
| 1418 | c96846_g1 | -9.272 | 0.0307605 |
| 1419 | c53354_g1 | -9.271 | 0.0307605 |
| 1420 | c31243_g1 | -9.270 | 0.0307605 |
| 1421 | c55069_g1 | -9.270 | 0.0307605 |
| 1422 | c56499_g1 | -9.267 | 0.0329919 |
| 1423 | c26950_g1 | -9.265 | 0.0329919 |

| | | | |
|---|---|---|---|
| 1424 | c33943_g1 | -9.264 | 0.0329919 |
| 1425 | c56392_g2 | -9.263 | 0.0329919 |
| 1426 | c46899_g1 | -9.263 | 0.0329919 |
| 1427 | c50344_g1 | -9.263 | 0.0329919 |
| 1428 | c46214_g1 | -9.262 | 0.0329919 |
| 1429 | c44722_g1 | -9.254 | 0.0329919 |
| 1430 | c52265_g1 | -9.252 | 0.0329919 |
| 1431 | c37891_g1 | -9.252 | 0.0329919 |
| 1432 | c48410_g1 | -9.249 | 0.0329919 |
| 1433 | c55142_g5 | -9.248 | 0.0329919 |
| 1434 | c54824_g1 | -9.244 | 0.0351786 |
| 1435 | c54431_g1 | -9.242 | 0.0351786 |
| 1436 | c23742_g1 | -9.242 | 0.0351786 |
| 1437 | c54430_g1 | -9.240 | 0.0351786 |
| 1438 | c47317_g1 | -9.239 | 0.0351786 |
| 1439 | c10568_g1 | -9.235 | 0.0351786 |
| 1440 | c56117_g3 | -9.234 | 0.0351786 |
| 1441 | c43737_g1 | -9.234 | 0.0351786 |
| 1442 | c53395_g1 | -9.233 | 0.0351786 |
| 1443 | c47506_g1 | -9.232 | 0.0351786 |
| 1444 | c53671_g1 | -9.232 | 0.0351786 |
| 1445 | c52161_g1 | -9.232 | 0.0351786 |
| 1446 | c53513_g1 | -9.227 | 0.0351786 |
| 1447 | c41696_g1 | -9.227 | 0.0351786 |
| 1448 | c55075_g2 | -9.226 | 0.0351786 |
| 1449 | c57339_g1 | -9.226 | 0.0351786 |
| 1450 | c54886_g1 | -9.225 | 0.0351786 |
| 1451 | c47291_g1 | -9.224 | 0.0351786 |
| 1452 | c16562_g1 | -9.218 | 0.0375986 |
| 1453 | c41380_g1 | -9.216 | 0.0375986 |
| 1454 | c49606_g1 | -9.213 | 0.0375986 |
| 1455 | c56204_g1 | -9.210 | 0.0375986 |
| 1456 | c48799_g1 | -9.209 | 0.0375986 |
| 1457 | c51228_g2 | -9.202 | 0.0375986 |
| 1458 | c50153_g1 | -9.202 | 0.0375986 |
| 1459 | c84052_g1 | -9.201 | 0.0375986 |
| 1460 | c39173_g1 | -9.201 | 0.0375986 |
| 1461 | c49504_g1 | -9.201 | 0.0375986 |
| 1462 | c56866_g1 | -9.199 | 0.0404152 |
| 1463 | c71604_g1 | -9.199 | 0.0375986 |
| 1464 | c45720_g1 | -9.197 | 0.0404152 |
| 1465 | c56292_g1 | -9.195 | 0.0404152 |
| 1466 | c41198_g1 | -9.193 | 0.0404152 |
| 1467 | c54424_g4 | -9.192 | 0.0404152 |
| 1468 | c57781_g1 | -9.190 | 0.0404152 |
| 1469 | c41973_g1 | -9.188 | 0.0404152 |
| 1470 | c56087_g1 | -9.186 | 0.0404152 |
| 1471 | c53085_g1 | -9.185 | 0.0404152 |
| 1472 | c56397_g3 | -9.183 | 0.0404152 |
| 1473 | c54485_g1 | -9.180 | 0.0404152 |
| 1474 | c56686_g3 | -9.178 | 0.0404152 |
| 1475 | c56800_g1 | -9.178 | 0.0404152 |
| 1476 | c34140_g1 | -9.173 | 0.0433703 |
| 1477 | c41767_g1 | -9.173 | 0.0404152 |
| 1478 | c53583_g2 | -9.171 | 0.0433703 |
| 1479 | c55132_g1 | -9.170 | 0.0433703 |
| 1480 | c71328_g1 | -9.170 | 0.0433703 |
| 1481 | c54641_g2 | -9.169 | 0.0404152 |
| 1482 | c55655_g4 | -9.169 | 0.0433703 |
| 1483 | c38325_g1 | -9.168 | 0.0433703 |
| 1484 | c57075_g3 | -9.167 | 0.0433703 |
| 1485 | c48910_g1 | -9.167 | 0.0433703 |

| | | | |
|---|---|---|---|
| 1486 | c56712_g2 | -9.166 | 0.0433703 |
| 1487 | c52174_g1 | -9.165 | 0.0433703 |
| 1488 | c55778_g1 | -9.164 | 0.0433703 |
| 1489 | c42772_g1 | -9.164 | 0.0433703 |
| 1490 | c56865_g1 | -9.164 | 0.0433703 |
| 1491 | c53508_g1 | -9.160 | 0.0433703 |
| 1492 | c55987_g1 | -9.160 | 0.0433703 |
| 1493 | c54398_g1 | -9.159 | 0.0404152 |
| 1494 | c57531_g2 | -9.159 | 0.0433703 |
| 1495 | c55407_g3 | -9.159 | 0.0433703 |
| 1496 | c57649_g1 | -9.158 | 0.0433703 |
| 1497 | c45595_g1 | -9.156 | 0.0433703 |
| 1498 | c55890_g1 | -9.151 | 0.0433703 |
| 1499 | c52307_g1 | -9.149 | 0.0433703 |
| 1500 | c50085_g1 | -9.149 | 0.0463549 |
| 1501 | c56028_g1 | -9.148 | 0.0463549 |
| 1502 | c32558_g1 | -9.146 | 0.0463549 |
| 1503 | c46715_g1 | -9.146 | 0.0463549 |
| 1504 | c54013_g4 | -9.144 | 0.0463549 |
| 1505 | c50448_g1 | -9.143 | 0.0463549 |
| 1506 | c56643_g1 | -9.140 | 0.0463549 |
| 1507 | c49448_g1 | -9.140 | 0.0463549 |
| 1508 | c47095_g1 | -9.138 | 0.0463549 |
| 1509 | c55224_g1 | -9.138 | 0.0463549 |
| 1510 | c36776_g1 | -9.138 | 0.0463549 |
| 1511 | c36339_g1 | -9.138 | 0.0463549 |
| 1512 | c52463_g1 | -9.136 | 0.0463549 |
| 1513 | c10628_g1 | -9.135 | 0.0463549 |
| 1514 | c49153_g1 | -9.134 | 0.0463549 |
| 1515 | c46230_g1 | -9.132 | 0.0463549 |
| 1516 | c26497_g1 | -9.131 | 0.0463549 |
| 1517 | c49417_g1 | -9.127 | 0.0463549 |
| 1518 | c54862_g1 | -9.126 | 0.049328 |
| 1519 | c51814_g1 | -9.123 | 0.049328 |
| 1520 | c55513_g3 | -9.122 | 0.049328 |
| 1521 | c47867_g1 | -9.122 | 0.0463549 |
| 1522 | c54218_g2 | -9.121 | 0.0463549 |
| 1523 | c54210_g1 | -9.120 | 0.049328 |
| 1524 | c23655_g1 | -9.120 | 0.049328 |
| 1525 | c51115_g1 | -9.119 | 0.049328 |
| 1526 | c34100_g1 | -9.119 | 0.049328 |
| 1527 | c51138_g1 | -9.118 | 0.049328 |
| 1528 | c56213_g1 | -9.118 | 0.0463549 |
| 1529 | c55362_g1 | -9.118 | 0.049328 |
| 1530 | c47969_g1 | -9.118 | 0.049328 |
| 1531 | c52143_g1 | -9.117 | 0.049328 |
| 1532 | c53657_g2 | -9.116 | 0.049328 |
| 1533 | c57145_g1 | -9.115 | 0.049328 |
| 1534 | c53545_g3 | -9.114 | 0.049328 |
| 1535 | c45033_g1 | -9.113 | 0.049328 |
| 1536 | c47182_g1 | -9.113 | 0.049328 |
| 1537 | c52862_g1 | -9.113 | 0.049328 |
| 1538 | c57386_g1 | -9.108 | 0.049328 |
| 1539 | c45387_g1 | -9.107 | 0.049328 |
| 1540 | c40964_g1 | -9.106 | 0.049328 |
| 1541 | c53490_g1 | -9.102 | 0.049328 |
| 1542 | c44053_g1 | -9.096 | 0.049328 |
| 1543 | c41007_g1 | -9.091 | 0.049328 |
| 1544 | c44892_g1 | -9.034 | 0.0404152 |
| 1545 | c56120_g1 | -9.034 | 0.0404152 |
| 1546 | c48399_g1 | -9.033 | 0.0404152 |
| 1547 | c49720_g1 | -9.029 | 0.0404152 |

| | | | |
|---|---|---|---|
| 1548 | c48555_g1 | -9.028 | 0.0404152 |
| 1549 | c45362_g1 | -9.027 | 0.0404152 |
| 1550 | c52969_g1 | -9.027 | 0.0404152 |
| 1551 | c45708_g1 | -9.025 | 0.0404152 |
| 1552 | c37106_g1 | -9.024 | 0.0404152 |
| 1553 | c34305_g1 | -9.024 | 0.0404152 |
| 1554 | c55277_g1 | -9.023 | 0.0404152 |
| 1555 | c32215_g1 | -9.016 | 0.0435221 |
| 1556 | c54228_g2 | -9.015 | 0.0435221 |
| 1557 | c56371_g1 | -9.012 | 0.0435221 |
| 1558 | c44904_g1 | -9.011 | 0.0435221 |
| 1559 | c55035_g1 | -9.009 | 0.0435221 |
| 1560 | c29083_g1 | -9.008 | 0.0435221 |
| 1561 | c47897_g1 | -9.007 | 0.0435221 |
| 1562 | c54070_g1 | -9.007 | 0.0435221 |
| 1563 | c50878_g1 | -9.006 | 0.0435221 |
| 1564 | c32455_g1 | -9.003 | 0.0435221 |
| 1565 | c55220_g3 | -9.003 | 0.0435221 |
| 1566 | c32480_g1 | -9.001 | 0.0435221 |
| 1567 | c23833_g1 | -9.001 | 0.0435221 |
| 1568 | c37504_g1 | -8.997 | 0.0435221 |
| 1569 | c52367_g1 | -8.996 | 0.0435221 |
| 1570 | c52004_g1 | -8.996 | 0.0435221 |
| 1571 | c42127_g1 | -8.995 | 0.046692 |
| 1572 | c57306_g4 | -8.994 | 0.046692 |
| 1573 | c51733_g1 | -8.991 | 0.046692 |
| 1574 | c48513_g1 | -8.990 | 0.046692 |
| 1575 | c47404_g1 | -8.990 | 0.046692 |
| 1576 | c52831_g1 | -8.987 | 0.046692 |
| 1577 | c23039_g1 | -8.987 | 0.046692 |
| 1578 | c57624_g1 | -8.985 | 0.046692 |
| 1579 | c56291_g2 | -8.983 | 0.046692 |
| 1580 | c46524_g1 | -8.982 | 0.046692 |
| 1581 | c40022_g1 | -8.982 | 0.046692 |
| 1582 | c57734_g1 | -8.981 | 0.046692 |
| 1583 | c57653_g1 | -8.978 | 0.046692 |
| 1584 | c39927_g1 | -8.976 | 0.046692 |
| 1585 | c51189_g1 | -8.976 | 0.046692 |
| 1586 | c46050_g1 | -8.974 | 0.046692 |
| 1587 | c16657_g1 | -8.974 | 0.046692 |
| 1588 | c44092_g1 | -8.974 | 0.046692 |
| 1589 | c57382_g1 | -8.974 | 0.046692 |
| 1590 | c49610_g1 | -8.971 | 0.046692 |
| 1591 | c10169_g1 | -8.971 | 0.046692 |
| 1592 | c26673_g1 | -8.970 | 0.046692 |
| 1593 | c47847_g1 | -8.969 | 0.046692 |
| 1594 | c43940_g1 | -8.968 | 0.046692 |
| 1595 | c48671_g1 | -8.968 | 0.046692 |
| 1596 | c44737_g1 | -8.960 | 0.046692 |
| 1597 | c31616_g1 | -7.926 | 3.09E-12 |
| 1598 | c26772_g1 | -6.425 | 3.25E-08 |
| 1599 | c36796_g1 | -5.728 | 1.26E-06 |
| 1600 | c11124_g1 | -5.694 | 1.52E-06 |
| 1601 | c50761_g3 | -5.321 | 1.09E-05 |
| 1602 | c46036_g2 | -5.188 | 9.13E-07 |
| 1603 | c56113_g1 | -5.165 | 2.41E-05 |
| 1604 | c96847_g1 | -5.158 | 2.47E-05 |
| 1605 | c57332_g3 | -5.154 | 2.50E-05 |
| 1606 | c54405_g1 | -5.140 | 2.36E-05 |
| 1607 | c26670_g1 | -5.053 | 4.47E-07 |
| 1608 | c53364_g1 | -4.928 | 3.69E-06 |
| 1609 | c49675_g1 | -4.927 | 6.30E-05 |

| | | | |
|---|---|---|---|
| 1610 | c49801_g1 | -4.908 | 6.92E-05 |
| 1611 | c52007_g1 | -4.775 | 8.15E-06 |
| 1612 | c47303_g1 | -4.686 | 1.34E-05 |
| 1613 | c41162_g1 | -4.659 | 0.000172 |
| 1614 | c10660_g1 | -4.641 | 1.66E-05 |
| 1615 | c14192_g1 | -4.582 | 2.07E-07 |
| 1616 | c56877_g1 | -4.563 | 0.0002736 |
| 1617 | c51321_g1 | -4.559 | 0.0002787 |
| 1618 | c56973_g1 | -4.502 | 3.77E-07 |
| 1619 | c31126_g1 | -4.420 | 2.56E-06 |
| 1620 | c34906_g1 | -4.397 | 2.92E-06 |
| 1621 | c33259_g1 | -4.386 | 0.0005789 |
| 1622 | c109563_g1 | -4.362 | 5.68E-05 |
| 1623 | c57566_g4 | -4.253 | 0.000887 |
| 1624 | c19814_g1 | -4.225 | 1.34E-05 |
| 1625 | c122464_g1 | -4.077 | 1.79E-06 |
| 1626 | c57020_g1 | -4.068 | 0.0018625 |
| 1627 | c55495_g3 | -4.008 | 1.05E-05 |
| 1628 | c47094_g1 | -3.963 | 6.55E-06 |
| 1629 | c53904_g1 | -3.898 | 0.0032177 |
| 1630 | c71341_g1 | -3.896 | 0.0001249 |
| 1631 | c37675_g1 | -3.835 | 1.11E-05 |
| 1632 | c57171_g1 | -3.825 | 2.02E-05 |
| 1633 | c53094_g4 | -3.824 | 0.0005417 |
| 1634 | c47318_g1 | -3.815 | 3.70E-05 |
| 1635 | c57475_g1 | -3.801 | 5.68E-05 |
| 1636 | c40774_g1 | -3.764 | 2.66E-05 |
| 1637 | c54734_g1 | -3.759 | 7.21E-06 |
| 1638 | c39277_g1 | -3.754 | 0.0001118 |
| 1639 | c50896_g1 | -3.708 | 0.000136 |
| 1640 | c52782_g1 | -3.619 | 0.0081345 |
| 1641 | c50545_g1 | -3.589 | 5.03E-05 |
| 1642 | c96868_g1 | -3.554 | 8.64E-05 |
| 1643 | c43517_g1 | -3.519 | 6.98E-05 |
| 1644 | c46691_g2 | -3.419 | 0.0004993 |
| 1645 | c56779_g4 | -3.397 | 0.0155187 |
| 1646 | c42820_g1 | -3.349 | 0.0012052 |
| 1647 | c47729_g4 | -3.245 | 0.000128 |
| 1648 | c53966_g2 | -3.233 | 0.0010037 |
| 1649 | c46386_g1 | -3.228 | 0.0242045 |
| 1650 | c15133_g1 | -3.217 | 0.0005054 |
| 1651 | c57540_g2 | -3.166 | 0.0062667 |
| 1652 | c10634_g1 | -3.161 | 0.0001381 |
| 1653 | c29272_g1 | -3.152 | 0.0014153 |
| 1654 | c47773_g1 | -3.121 | 0.0028786 |
| 1655 | c44943_g1 | -3.118 | 0.031145 |
| 1656 | c56762_g4 | -3.104 | 0.0030608 |
| 1657 | c51296_g1 | -3.093 | 9.45E-05 |
| 1658 | c71319_g1 | -3.092 | 0.0330762 |
| 1659 | c58095_g1 | -3.091 | 0.0075571 |
| 1660 | c41358_g1 | -3.066 | 0.0012816 |
| 1661 | c46881_g1 | -3.018 | 0.0043717 |
| 1662 | c50253_g1 | -3.006 | 0.0394999 |
| 1663 | c37438_g1 | -2.960 | 0.0003723 |
| 1664 | c44410_g1 | -2.935 | 0.000595 |
| 1665 | c51845_g4 | -2.933 | 0.0055492 |
| 1666 | c57516_g2 | -2.919 | 0.0006978 |
| 1667 | c52641_g1 | -2.895 | 0.0014538 |
| 1668 | c109566_g1 | -2.894 | 0.0003002 |
| 1669 | c10543_g1 | -2.812 | 0.0005412 |
| 1670 | c41220_g1 | -2.803 | 0.0082834 |
| 1671 | c51099_g2 | -2.760 | 0.0003922 |

| | | | |
|---|---|---|---|
| 1672 | c43584_g1 | -2.718 | 0.0230376 |
| 1673 | c53380_g2 | -2.679 | 0.0008362 |
| 1674 | c56418_g1 | -2.669 | 0.0133738 |
| 1675 | c38993_g1 | -2.665 | 0.002129 |
| 1676 | c46700_g1 | -2.603 | 0.0098386 |
| 1677 | c43581_g2 | -2.583 | 0.0014631 |
| 1678 | c52990_g1 | -2.581 | 0.0057455 |
| 1679 | c42642_g1 | -2.561 | 0.0181507 |
| 1680 | c27278_g1 | -2.551 | 0.0019437 |
| 1681 | c35005_g2 | -2.471 | 0.0228912 |
| 1682 | c55006_g2 | -2.451 | 0.0244192 |
| 1683 | c36468_g5 | -2.450 | 0.0041969 |
| 1684 | c39630_g1 | -2.433 | 0.0059808 |
| 1685 | c24424_g1 | -2.426 | 0.0020848 |
| 1686 | c109579_g1 | -2.294 | 0.0386624 |
| 1687 | c52011_g1 | -2.281 | 0.0403739 |
| 1688 | c71688_g1 | -2.253 | 0.0439429 |
| 1689 | c54058_g1 | -2.232 | 0.0230146 |
| 1690 | c54274_g1 | -2.214 | 0.0143954 |
| 1691 | c23686_g1 | -2.203 | 0.0055315 |
| 1692 | c45178_g1 | -2.164 | 0.009313 |
| 1693 | c57583_g2 | -2.132 | 0.0415549 |
| 1694 | c47898_g2 | -2.099 | 0.0104891 |
| 1695 | c56770_g1 | -2.065 | 0.0267115 |
| 1696 | c43562_g1 | -2.046 | 0.0093258 |
| 1697 | c25467_g1 | -1.943 | 0.0093721 |
| 1698 | c47992_g1 | -1.933 | 0.023326 |
| 1699 | c56133_g1 | -1.927 | 0.0124088 |
| 1700 | c54123_g3 | -1.875 | 0.0142615 |
| 1701 | c50790_g2 | -1.842 | 0.0178985 |
| 1702 | c42746_g1 | -1.777 | 0.0227728 |
| 1703 | c53698_g2 | -1.732 | 0.0281783 |
| 1704 | c58238_g3 | -1.618 | 0.0307998 |
| 1705 | c43857_g2 | -1.543 | 0.043474 |
| 1706 | c54742_g1 | 1.410 | 0.031145 |
| 1707 | c55046_g1 | 1.566 | 0.0260766 |
| 1708 | c51812_g1 | 1.568 | 0.0292992 |
| 1709 | c55881_g1 | 1.578 | 0.0182258 |
| 1710 | c57979_g1 | 1.628 | 0.031509 |
| 1711 | c52989_g1 | 1.684 | 0.0106309 |
| 1712 | c56584_g3 | 1.760 | 0.010319 |
| 1713 | c57747_g1 | 1.801 | 0.038702 |
| 1714 | c37995_g1 | 1.801 | 0.0094271 |
| 1715 | c24300_g1 | 1.827 | 0.0107089 |
| 1716 | c54779_g3 | 1.902 | 0.0063219 |
| 1717 | c56952_g1 | 1.963 | 0.0026144 |
| 1718 | c42736_g1 | 1.972 | 0.0057707 |
| 1719 | c58134_g1 | 2.000 | 0.0028184 |
| 1720 | c23656_g1 | 2.016 | 0.0270781 |
| 1721 | c57605_g1 | 2.195 | 0.0032718 |
| 1722 | c54682_g3 | 2.216 | 0.007639 |
| 1723 | c51910_g1 | 2.246 | 0.0004146 |
| 1724 | c53268_g1 | 2.260 | 0.0009829 |
| 1725 | c54463_g2 | 2.356 | 0.0005279 |
| 1726 | c42361_g1 | 2.360 | 0.0008207 |
| 1727 | c54468_g2 | 2.411 | 0.0067386 |
| 1728 | c42782_g1 | 2.430 | 0.00161 |
| 1729 | c57484_g1 | 2.431 | 0.00211 |
| 1730 | c39298_g1 | 2.476 | 0.0002717 |
| 1731 | c49219_g1 | 2.482 | 0.0075571 |
| 1732 | c51652_g2 | 2.488 | 0.0003037 |
| 1733 | c39199_g1 | 2.528 | 9.60E-05 |

| | | | |
|---|---|---|---|
| 1734 | c31940_g1 | 2.542 | 0.0001824 |
| 1735 | c57284_g1 | 2.550 | 0.0003453 |
| 1736 | c49653_g1 | 2.551 | 0.0006306 |
| 1737 | c56315_g3 | 2.642 | 0.0023408 |
| 1738 | c84190_g1 | 2.681 | 0.0001289 |
| 1739 | c38882_g1 | 2.760 | 0.0002073 |
| 1740 | c27267_g1 | 2.792 | 5.68E-05 |
| 1741 | c51872_g1 | 2.871 | 1.07E-05 |
| 1742 | c55776_g1 | 2.941 | 0.0147433 |
| 1743 | c19787_g1 | 2.955 | 6.83E-06 |
| 1744 | c50415_g1 | 2.962 | 1.59E-05 |
| 1745 | c71572_g1 | 2.975 | 2.95E-05 |
| 1746 | c53266_g1 | 3.005 | 3.56E-06 |
| 1747 | c52251_g1 | 3.023 | 5.63E-05 |
| 1748 | c55656_g1 | 3.067 | 0.0005475 |
| 1749 | c46247_g1 | 3.086 | 0.001042 |
| 1750 | c53438_g1 | 3.104 | 9.80E-07 |
| 1751 | c53257_g1 | 3.143 | 0.0005789 |
| 1752 | c35299_g1 | 3.146 | 1.78E-06 |
| 1753 | c53083_g1 | 3.203 | 4.48E-07 |
| 1754 | c53144_g1 | 3.238 | 1.56E-05 |
| 1755 | c54457_g1 | 3.253 | 4.28E-07 |
| 1756 | c54646_g1 | 3.256 | 3.54E-07 |
| 1757 | c50577_g1 | 3.261 | 1.70E-07 |
| 1758 | c53629_g1 | 3.330 | 1.57E-06 |
| 1759 | c51080_g1 | 3.368 | 2.50E-05 |
| 1760 | c51692_g1 | 3.401 | 7.28E-07 |
| 1761 | c14005_g2 | 3.437 | 3.13E-08 |
| 1762 | c45564_g1 | 3.452 | 1.62E-07 |
| 1763 | c84079_g1 | 3.483 | 7.32E-08 |
| 1764 | c54144_g1 | 3.494 | 2.22E-05 |
| 1765 | c54397_g1 | 3.504 | 2.94E-07 |
| 1766 | c53532_g3 | 3.579 | 8.28E-08 |
| 1767 | c52653_g1 | 3.590 | 4.31E-07 |
| 1768 | c50506_g1 | 3.668 | 3.52E-08 |
| 1769 | c33810_g2 | 3.684 | 0.0011239 |
| 1770 | c49686_g1 | 3.695 | 0.0019138 |
| 1771 | c31520_g1 | 3.700 | 4.02E-06 |
| 1772 | c51512_g1 | 3.921 | 1.44E-08 |
| 1773 | c52696_g1 | 3.923 | 1.56E-05 |
| 1774 | c54042_g1 | 3.923 | 3.25E-08 |
| 1775 | c57790_g3 | 3.957 | 2.74E-09 |
| 1776 | c55244_g1 | 3.957 | 3.51E-10 |
| 1777 | c51276_g18 | 4.090 | 1.37E-10 |
| 1778 | c36300_g1 | 4.150 | 3.27E-05 |
| 1779 | c39444_g1 | 4.292 | 1.52E-06 |
| 1780 | c10762_g2 | 4.364 | 1.10E-11 |
| 1781 | c48522_g1 | 4.381 | 6.86E-11 |
| 1782 | c43382_g2 | 4.489 | 0.0080393 |
| 1783 | c52058_g2 | 4.499 | 8.04E-11 |
| 1784 | c49047_g1 | 4.551 | 2.23E-12 |
| 1785 | c56291_g1 | 4.694 | 8.14E-13 |
| 1786 | c46847_g1 | 4.821 | 4.66E-13 |
| 1787 | c56959_g1 | 5.169 | 0.0390317 |
| 1788 | c56893_g2 | 5.300 | 1.18E-15 |
| 1789 | c32378_g1 | 5.395 | 7.52E-15 |
| 1790 | c84143_g1 | 5.410 | 8.39E-16 |
| 1791 | c47324_g1 | 5.445 | 0.0390317 |
| 1792 | c26502_g2 | 5.468 | 6.33E-10 |
| 1793 | c46661_g1 | 6.038 | 0.0019875 |
| 1794 | c18631_g1 | 6.141 | 1.71E-17 |
| 1795 | c55650_g1 | 6.203 | 1.06E-13 |

| | | | |
|---|---|---|---|
| 1796 | c51276_g6 | 6.303 | 1.13E-14 |
| 1797 | c52890_g1 | 6.409 | 0.0019875 |
| 1798 | c57109_g1 | 6.866 | 3.50E-22 |
| 1799 | c54365_g1 | 7.085 | 3.36E-14 |
| 1800 | c52517_g1 | 7.145 | 0.0001588 |
| 1801 | c39339_g1 | 7.631 | 7.28E-07 |
| 1802 | c50124_g1 | 8.094 | 9.74E-28 |
| 1803 | c46260_g1 | 8.140 | 8.61E-26 |
| 1804 | c49026_g1 | 8.438 | 2.42E-27 |
| 1805 | c55419_g1 | 8.757 | 2.42E-27 |
| 1806 | c56514_g1 | 8.820 | 2.81E-34 |
| 1807 | c58545_g1 | 9.004 | 0.0215887 |
| 1808 | c58307_g1 | 9.049 | 1.89E-31 |
| 1809 | c134138_g1 | 9.588 | 0.0035389 |
| 1810 | c14276_g1 | 9.593 | 5.37E-25 |
| 1811 | c38226_g1 | 9.868 | 5.37E-25 |
| 1812 | c55028_g1 | 10.003 | 0.0011562 |
| 1813 | c31740_g1 | 10.003 | 0.0011562 |
| 1814 | c46433_g1 | 10.059 | 1.95E-30 |
| 1815 | c37120_g1 | 10.095 | 6.19E-39 |
| 1816 | c84017_g1 | 10.423 | 4.68E-22 |
| 1817 | c50147_g1 | 10.454 | 1.16E-40 |
| 1818 | c103700_g1 | 10.757 | 4.28E-48 |
| 1819 | c10794_g1 | 10.981 | 7.07E-40 |
| 1820 | c58232_g1 | 11.324 | 3.74E-07 |
| 1821 | c114314_g1 | 11.467 | 1.43E-45 |
| 1822 | c77614_g1 | 12.250 | 4.66E-13 |
| 1823 | c129600_g1 | 13.250 | 1.95E-23 |
| 1824 | c53455_g1 | 13.493 | 1.93E-25 |
| 1825 | c47749_g1 | 13.545 | 1.04E-38 |
