## Supplementary material for "Decoding the reproductive system of the olive fruit fly, *Bactrocera oleae*"

| No | B. oleae | Scaffold | Functional class |
|---|---|---|---|
| 1 | alphaTub84B | NW_013581225.1.2 | sperm protein |
| 2 | betaTub85D | NW_013581217.1.128 | sperm protein |
| 3 | Ccp84Ad | NW_013583085.1.13 | chitin binding |
| 4 | Cdlc2 | NW_013581488.1.1 | sperm protein |
| 5 | CG10407-like | NW_013581506.1.10 | unknown function |
| 6 | CG10433-like | NW_013581250.1.1 | defense/immunity |
| 7 | CG10730-like | NW_013581262.1.37 | unknown function |
| 8 | CG11598-PB-like | NW_013581987.1.8 | lipid metabolism |
| 9 | CG11864-like | NW_013581224.1.5 | protease |
| 10 | CG13340-like | NW_013581355.1.7 | protease |
| 11 | CG15031-like | NW_013581935.1.1 | unknown function |
| 12 | CG15116-like | NW_013581511.1.8 | defense/immunity |
| 13 | CG15117-like | NW_013582880.1.1 | carbohydrate metabolism |
| 14 | CG17843-like | NW_013581762.1.2 | oxidative stress response |
| 15 | CG17919-like | NW_013581353.1.17 | signal transduction |
| 16 | CG18135-like | NW_013581267.1.22 | unknown function |
| 17 | CG18284-like | NW_013581453.1.14 | lipid metabolism |
| 18 | CG18628-like | NW_013581453.1.14 | unknown function |
| 19 | CG2852-like | NW_013581924.1.1 | protein modification |
| 20 | CG3153-like | NW_013581262.1.25 | unknown function |
| 21 | CG31704-like | NW_013582946.1.1 | protease inhibitor |
| 22 | CG31758-like | NW_013599804.1.1 | protease inhibitor |
| 23 | CG4847-like | NW_013581323.1.30 | protease |
| 24 | CG5162-like | NW_013586638.1.2 | lipid metabolism |
| 25 | CG6426-like | NW_013581234.1.7 | defense/immunity |
| 26 | CG6461-like | NW_013581236.1.27 | protease |
| 27 | CG8102-like | NW_013582722.1.1 | sperm protein |
| 28 | CG9168-like | NW_013581262.1.39 | unknown function |
| 29 | CG9975-like | NW_013582087.1.1 | unknown function |
| 30 | Cpr51A | NW_013581314.1.29 | chitin binding |
| 31 | Cpr67Fb | NW_013584326.1.1 | chitin binding |
| 32 | Egm | NW_013581265.1.12 | oxidative stress response |
| 33 | Est-6 | NW_013581551.1.20 | lipid metabolism |
| 34 | Hexo2 | NW_013581256.1.14 | carbohydrate metabolism |
| 35 | mfas-PB | NW_013581231.1.21 | signal transduction |
| 36 | NUCB1 | NW_013581534.1.9 | calcium binding |
| 37 | Peb | NW_013581247.1.21 | post-mating behavior |
| 38 | Peritrophin-A | NW_013581236.1.13 | chitin binding |
| 39 | Phm | NW_013583124.1.1 | protein modification |
| 40 | regucalcin | NW_013581216.1.74 | defense/immunity |
| 41 | Spn1 | NW_013581585.1.8 | protease inhibitor |
| 42 | trx | NW_013581231.1.1 | DNA interactions |
| 43 | Or82a | NW_013581351.1.3 | odorant binding |
| 44 | alphaTub84B | NW_013581225.1.2 | sperm protein |
