## Supplementary material for "Decoding the reproductive system of the olive fruit fly, *Bactrocera oleae*"

| Gene | Primers |
|---|---|
| Lower female reproductive tract | |
| *troponin C* | 5'-AAAACCAAGCCCATCCACC-3' |
| | 5'-GCGATTTGTTCGGGAGTCAG-3' |
| *yolk protein-2* | 5'-CGCGTATAGCCTAAAACCCAC-3' |
| | 5'-TGCAGGGTGATATCCTCCAC-3' |
| *lingerer* | 5'-CGCGTATAACTCGAGCGACTCC-3' |
| | 5'-GCGGCAGCTAATCGTCAATGC-3' |
| *glutathione S-transferase epsilon class* | 5'-ATGGCTTACCTGTCTAAATG-3' |
| | 5'-GTTTATTCCTCACTTTCACC-3' |
| *bestrophin 2* | 5'-AGGACATCCGACAACAACGGC-3' |
| | 5'-ATATTTGTGGTGACGGGCGCAG-3' |
| *ornithine decarboxylase antizyme* | 5'-ACGTTGCAATGCCTGACAAG-3' |
| | 5'-AACAACTGCGTCGACATCCA-3' |
| Male accessory glands/ ejaculatory bulb | |
| *brunelleschi* | 5'-AAGCGAGGTAACACTACGGC-3' |
| | 5'-GATTACCGTTTGTGGCAGCG-3' |
| *CG2254-like* | 5'-TATGCACATATGTATGCACATGAAA-3' |
| | 5'-ATGTTGCGCGCGTCTTTAG-3' |
| *timeless* | 5'-TGGCGGCGGACGTATAATAG-3' |
| | 5'-AAGTGCTCCCGTAGTTGGTG-3' |
| *c52416* | 5'-CGCTGTCACCACTGACTATGGC-3' |
| | 5'-TCCTCTGTCACCAGCTCAGAAAC-3' |
| *c53574* | 5'-GCATTTGCTGGCGCTTATCA-3' |
| | 5'-GCACAAACGGAAAGATGGCA-3' |
| *yellow-g* | 5'-TTGCGTGTTGGACAGGGTGC-3' |
| | 5'-AATTCGTGCCACCATCGGCG-3' |
| **Testes** | |
| *c15699* | 5'-CGAGAATATAAACGAACCTG-3' |
| | 5'-ATCACTTCAACTCTCTCTGTC-3' |
| *c58283* | 5'-AGTGAGTGATCCTGTACTGTC-3' |
| | 5'-TCGGTATACTCTACCTATCCAC-3' |
| *mucin* | 5'-CCAACCGACACAACGAAAGG-3' |
| | 5'-TGGCAAAGCCGCCAAAATAC-3' |
| *hemolectin* | 5'-CCAAATGCACAATTCACCAC-3; |
| | 5'-GCATCGTTCAGCACATATCC-3' |
| *c37552* | 5'-AGCGAAATAGTCCAGTTAGGTG-3' |
| | 5'-CCACACCAAACGATTACGGC-3' |
| *cation transporter* | 5'-ACTAAGTTTGGGTGTAACCG-3' |
| | 5'-GTGATACTTTCCGTAGTTTG-3' |
| *c42518* | 5'-GGCACCACATAAACTCTAAC-3' |
| | 5'-TGCACTCCGCTAATTGCC-3' |
| *scribbler isoform J* | 5'-GGTTTACTCCTTGCGTTGCC-3' |
| | 5'-CGGACCTCAAAACGATGCAC-3' |
| *c52071* | 5'-GCGCTTCATCATCCACAGAC-3' |
| | 5'-CGCTGTTAATACGCCACGC-3' |
| Housekeeping genes | |
| *RPL19* | 5'-CTTCACGTACTTTATGCCTTC-3' |
| | 5'-GCAAGGGTAATGTGTTCAA-3' |
| *GAPDH* | 5'-ATGAAGGTCGTATCTAATGC-3' |

| | |
|---|---|
| *GAPDH* | 5'-TAGTTGCGTGAACAGTAGTC-3' |

| Product size |
| --- |
| 98 |
| 80 |
| 124 |
| 114 |
| 105 |
| 101 |
| 75 |
| 73 |
| 87 |
| 111 |
| 112 |
| 97 |
| 150 |
| 98 |
| 125 |
| 140 |
| 82 |
| 120 |
| 100 |
| 83 |
| 135 |
| 126 |
| 115 |

[illegible]
